# Protein Editing using a Concerted Transposition Reaction

**DOI:** 10.1101/2024.06.03.597171

**Authors:** Yi Hua, Nicholas E. S. Tay, Xuanjia Ye, Jeremy A. Owen, Hengyuan Liu, Robert E. Thompson, Tom W. Muir

## Abstract

Protein engineering through the chemical or enzymatic ligation of polypeptide fragments has proven enormously powerful for studying countless biochemical processes *in vitro*. In general, this strategy necessitates a protein folding step following ligation of the unstructured fragments, a requirement that constrains the types of systems amenable to the approach. Here, we report an *in vitro* strategy that allows internal regions of target proteins to be replaced in a single operation. Conceptually, our system is analogous to a DNA transposition reaction, but employs orthogonal pairs of split inteins to swap out a designated region of a host protein with an exogenous molecular cassette. We show using isotopic labeling experiments that this ‘protein transposition’ reaction is concerted when the kinetics for the embedded intein pairs are suitably matched. Critically, this feature allows for efficient manipulation of protein primary structure in the context of a native fold. The utility of this method is illustrated using several protein systems including the multi-subunit chromatin remodeling complex, ACF, where we also show protein transposition can occur *in situ* within the cell nucleus. By carrying out a molecular ‘cut and paste’ on a protein or protein complex under native folding conditions, our approach dramatically expands the scope of protein semisynthesis.

---

The ability to assemble proteins from synthetic and recombinant fragments – i.e., semisynthesis – has had a tremendous impact on the biomedical sciences. Through the introduction of unnatural amino acids, post-translational modifications (PTMs), spectroscopic and biochemical probes, as well as combinations thereof, protein semisynthesis provides the means to tackle mechanistic problems that are difficult to address using other approaches(*1*). Despite these attributes, not every protein is a good candidate for *in vitro* semisynthesis using current strategies. While the size of the protein of interest (POI) is usually not a deciding factor, the location of the desired modification site within the primary sequence often can be. The vast majority of studies have focused on the installation of modifications within the N- or C-terminal proximal regions of the protein since this requires a single ligation step employing a short, synthetically accessible peptide fragment. By contrast, the introduction of modifications into the interior of a protein through semisynthesis is much more challenging, requiring multiple ligation steps employing three or more fragments (Fig. 1A). As with any multistep process, the overall yield can be modest due to accumulated losses during each successive ligation and purification step. Moreover, generation of the necessary fragments can be non-trivial. This is especially true for the flanking recombinant protein segments which, lacking the ability to adopt a native fold, can be hard to isolate in useful amounts (*2, 3*). Irrespective of the number of ligation steps employed, the final protein product must be able to efficiently fold into its native state following assembly of the fragments. This can be challenging for many proteins under *in vitro* conditions(*4, 5*). Furthermore, many proteins are part of large multi-subunit complexes, the reconstitution of which may not be feasible in a test-tube. Thus, there remain numerous systems where *in vitro* semisynthesis could be used to address specific functional or structural questions, but where the above constraints preclude such an undertaking.

**Fig 1.**
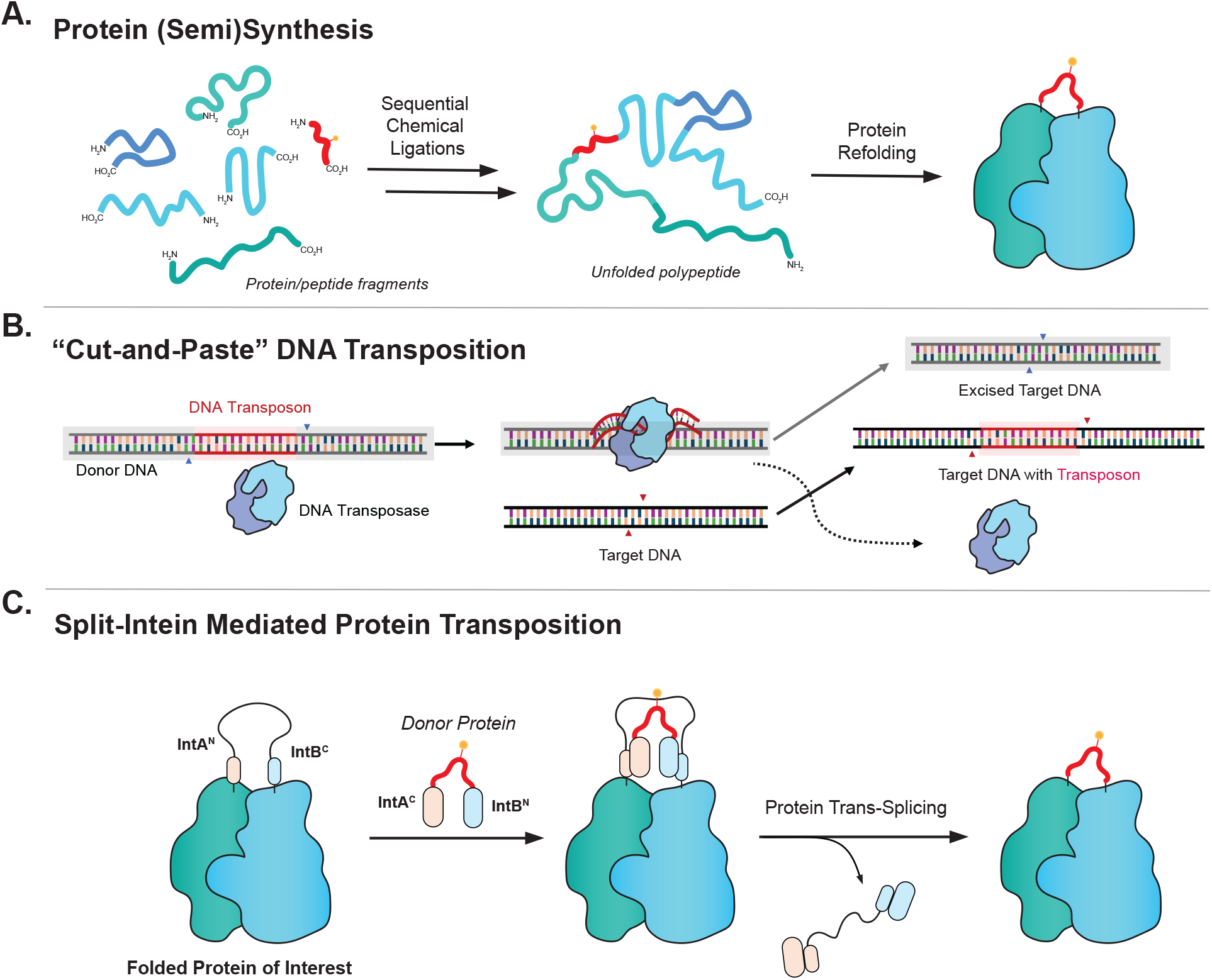
Design of a split intein-mediated protein transposition reaction. (a) Traditional protein semi(synthesis) using the stepwise assembly of multiple peptide fragments. Accumulated losses during the multi-step synthesis combined with an obligate final protein folding step limit the scope of this approach. (b) Schematic showing the convenience of the ‘Cut and Paste’ mechanism of DNA transposition. (c) A concerted protein transposition reaction carried out by a pair of orthogonal split inteins flanking the modification site. The approach circumvents the need to generate and manipulate protein fragments and does not require a folding step.

Motivated by the need for a more streamlined and broadly applicable *in vitro* method for performing semisynthesis on internal regions of a protein, including within complexes, we report here a strategy that allows such editing to be carried out in a concerted manner and without the need for a dedicated folding step. Our approach represents a conceptual departure from previous *in vitro* semisynthesis efforts(*1, 6-8*), which have involved stepwise assembly of individually prepared and purified protein fragments, thereby necessitating a final folding step (Fig. 1A). Rather, the current method takes inspiration from DNA transposition in which an exogenous transposon sequence is inserted into a recipient locus through the action of a transposase enzyme (*9*) (Fig. 1B). To adapt transposition to a protein context, we envisioned employing orthogonal pairs (i.e., they do not cross-react) of split inteins which undergo protein trans splicing (PTS) reactions(*10*) (Fig. S1). While PTS has been widely applied in the protein semisynthesis area (*1*), there have been only scattered reports of using this process to modify internal regions of proteins(*8, 11, 12*). Previous work from our lab employed a combination of naturally split and artificially split inteins to perform a one-pot protein assembly from three purified building blocks (*8*). However, the yield of this tandem PTS process was modest, making it of limited practical value. More recently, Pless and co-workers have used pairs of split inteins to modify, via a microinjection process, ion channels in individual Xenopus oocyte cells(*12, 13*). Again, the estimated yields of these tandem PTS reactions are low (<5%). Building on these earlier efforts, we wondered whether the efficiency of such insertion processes could be improved by using pairs of engineered split inteins with suitably tuned and balanced properties. This would allow for a more robust ‘protein transposition’ reaction in which a recipient protein containing strategically embedded pairs of split inteins is reacted with a ‘transposon’ construct containing the insert of interest flanked by the complementary split intein fragments (Fig. 1C). In principle, this *in vitro* reaction could be carried out in the context of a natively folded host protein or protein complex.

## Efficient Protein Transposition Requires Matched Split Intein Splicing Kinetics

Conceivably, the proposed transposition reaction could occur through one of two mechanisms: (i) a stepwise process, in which one intein pair splices before the other, or (ii) a concerted-type reaction whereby the two reactions occur concurrently (Fig. 2A). The latter scenario is more desirable since it would avoid the buildup intermediates which, by definition, constitute fragments of the recipient protein that could proceed down a misfolding pathway. Two factors must be considered here, namely the binding kinetics of the N- and C-terminal intein fragments (Int^N^ and Int^C^, respectively) and the rate of the ensuing protein trans-splicing reactions. Since both binding and splicing kinetics can vary significantly for different split inteins(*14-20*) the choice of which pair to use in the transposition reaction was likely to be critical for optimizing the system. Thus, our initial efforts were channeled towards identifying suitable split intein pairs.

**Fig 2.**
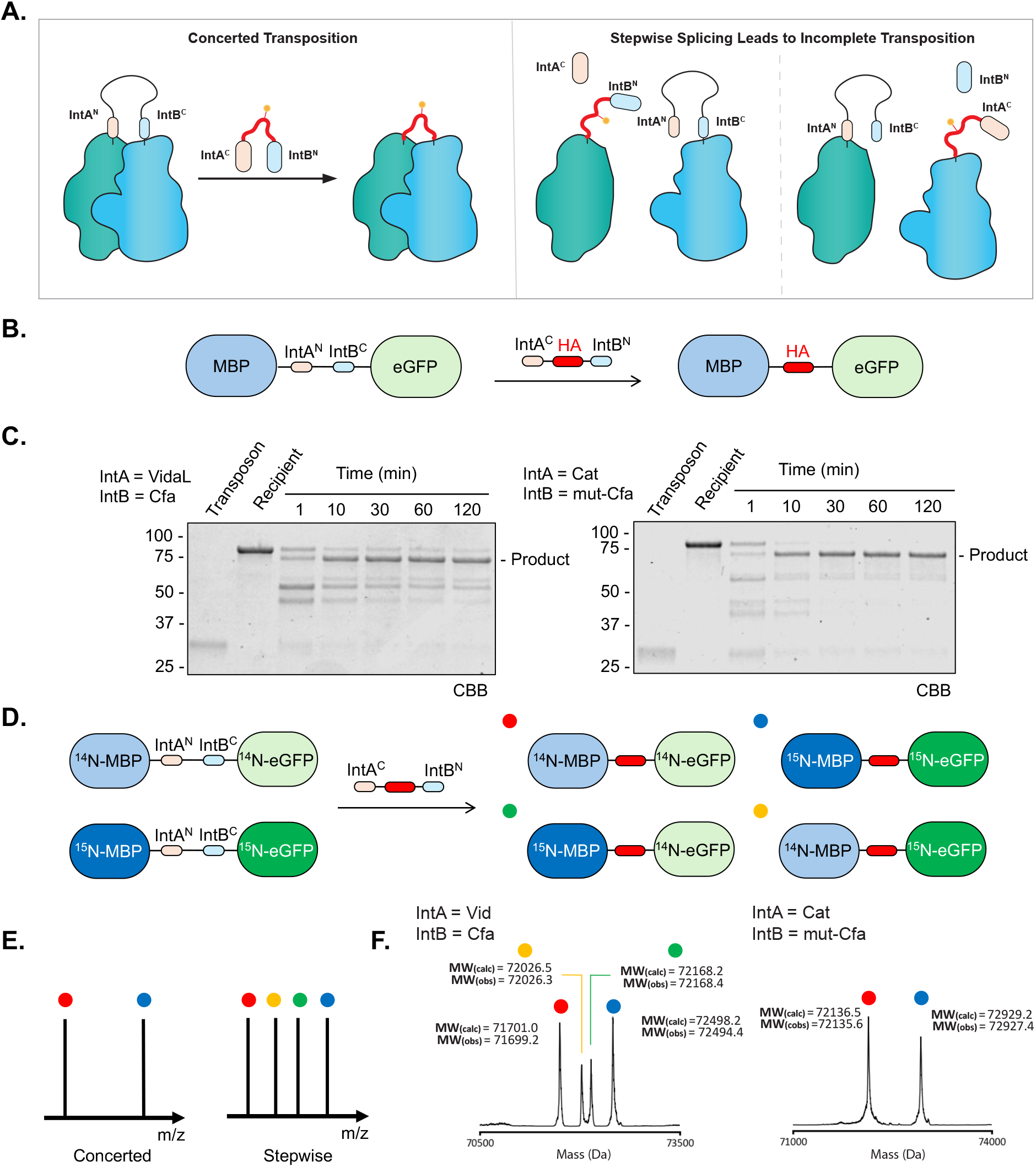
Efficient Protein Transposition Requires Matched Split-Intein Splicing Kinetics. (a) Product outcomes for protein transposition depend on split-intein splicing kinetics. Mismatched splicing kinetics for the two split inteins can lead to a build-up of unwanted reaction intermediates. (b) The design of MBP-eGFP model system for optimizing the protein transposition reaction. (c) Characterization of transposition reactions using the VidaL/Cfa split intein pair (left) and Cat/mut-Cfa pair (right). MBP-eGFP recipient and transposon donor constructs (2 μM of each) were reacted for the indicated times in 100 mM phosphate, 150 mM NaCl, 1 mM EDTA, 1 mM TCEP, pH 7.2. Reactions mixtures were analyzed by SDS-PAGE with Coomassie staining (CBB). The expected transposition product is indicated. Lower bands correspond to incomplete splicing intermediates. (d) The design of an isotopic labeling experiment to differentiate between stepwise and concerted processes. A 1:1 mixture of uniformly ^14^N or ^15^N labeled MBP-eGFP recipient proteins are reacted with the transposon construct. Up to four different isotopic compositions in the transposition product are possible depending on the reaction pathway. These kinetic outcomes are denoted by colored balls. (e) Anticipated product mass spectrometry readout distinguishing the concerted versus stepwise transposition pathways. (f) Mass spectrometry characterization of transposition products in an isotopic labeling experiment for different intein pairs. Each isotopically labeled recipient (2 μM total) is reacted with an equimolar amount of HA tag transposon (2 μM) for 2 hours in 100 mM phosphate, 150 mM NaCl, 1 mM EDTA, 1 mM TCEP, pH 7.2. After the transposition, the products were isolated by RP-HPLC and the isotopic compositions were detected by ESI-MS and deconvoluted. These experiments reveal that the VidaL/Cfa split intein pair proceeds via a stepwise process whereas the Cat/mut-Cfa pair enables almost fully concerted transposition.

To test the feasibility of the proposed transposition reaction, as well as the mechanism by which it might proceed, we designed a test system in which two orthogonal intein fragments, separated by a short linker containing a TEV protease cut site, were inserted between the model proteins, maltose binding protein (MBP) and enhanced green fluorescent protein (eGFP) (Fig. 2B, Fig. S2). Importantly, MBP and eGFP have no affinity for one another, minimizing any biasing effect that an interaction between the flanking regions of the recipient protein might have on the reaction pathway. This modular system allowed us to vary the split inteins employed in the transposition reaction with the goal of finding an optimal pair. To reduce the size of the embedded transposition apparatus within the recipient construct, we elected to use an atypically split intein on the N-terminal side (either Cat (*17*) or VidaL (*18*)) and a canonically split intein on the C-terminal side (Cfa(*15*)); the former have short Int^N^ sequences, whereas the opposite is true for the latter. A series of these MBP-IntA^N^-linker-IntB^C^-eGFP proteins were generated along with the corresponding transposon constructs, IntA^C^-insert-IntB^N^, containing the model insert (HA epitope tag) flanked by the complementary split inteins (Fig. 2B, Fig. S3). The matching constructs were then mixed in a 1:1 ratio and the reactions monitored over time by SDS-PAGE and LC-MS (Fig. 2C, Figs. S4-S5). In all cases, we observed generation of the expected transposition products, however, the efficiency of the reaction varied as a function of the split intein pairs employed. For instance, in the reaction involving the VidaL and Cfa split inteins, we observed a build-up of intermediates formed by single intein splicing (Figs. 2C and S4B); by contrast, these intermediates were less pronounced when we used Cat and Cfa (Fig. S5B).

We speculated that the differences in overall efficiency as function of split intein pairing might reflect alternate reaction pathways, stepwise *vs*. concerted. An experiment based on isotopic labeling and mass spectrometry was conducted to investigate this possibility (Fig. 2D). We generated two versions of the MBP-IntA^N^-linker-IntB^C^-eGFP recipient protein, one containing the natural nitrogen isotope (^14^N) and the other uniformly labeled with the heavier ^15^N stable isotope (Fig. 2E, Fig. S6). These two proteins were co-mixed at a 1:1 ratio and then reacted with corresponding transposon construct. Assuming a completely concerted pathway is taken, then the transposition product should retain either ^15^N or ^14^N labels in both MBP and eGFP, i.e. no isotopic mixing should occur and only two peaks should be observed in the mass spectrum at a 1:1 ratio. Conversely, if a purely stepwise pathway is operational then four products should be obtained, two with uniform isotopic labeling and two with mixed labeling (Fig. 2E). Control studies that invoked the TEV cleavage site within the linker region of the recipient protein indicated that the ratio of the four isotopic products in a purely stepwise process is 1:1:1:1 (Fig. S7). In the case of the reaction involving the VidaL and Cfa inteins, we observed four distinct masses within the product fraction. The ratio of these MS peaks was approximately 2:1:1:2, suggesting that a combination of concerted and stepwise pathways was operational (Fig. 2F, Fig. S8). When pairing Cat and Cfa together, we observed fewer products generated from the stepwise pathway (Fig. S9). To improve the concerted-to-stepwise ratio, we introduced a mutant into Cfa (mut-Cfa) predicted to have slower splicing kinetics (Cfa^N^ M75L, M81L – designated Cfa^N^_m_) compared to the wild-type (*15*), which would synchronize the splicing rates to that of both Cat and VidaL (Fig. S10) (*17, 18*). We observed that the transposition reactions involving mut-Cfa and either VidaL or Cat gave a very different isotopic distribution in the product (Fig. 2F, Figs. S8-9). In both cases, the concerted reaction dominated as reflected by the observation of two major peaks in the mass spectrum corresponding to the uniformly labeled species. We estimate that in the case of the mut-Cfa and Cat pairing that almost all reactions occurred by a concerted pathway, indicated that this pairing is well matched in terms of binding and splicing kinetics. Characterization by SDS-PAGE also showed reduced build-up of protein splicing intermediates when using mut-Cfa and Cat (Fig. 2C).

Notably, varying the ratio of reactants had minimal effect on conversion efficiency when using mut-Cfa and Cat (Fig. S11). Based on these results, we elected to use the Cat/mut-Cfa pairing in our subsequent studies.

### Protein Transposition Occurs Seamlessly on Folded Proteins

Next, we asked whether the transposition reaction can occur in the context of a folded recipient protein, as opposed to between two different folded proteins as in the previous example. To explore this, we inserted the Cat/mut-Cfa apparatus into a surface exposed loop in eGFP, specifically at position 173-174 (Fig. 3A). This site has previously been used in split eGFP constructs(*21*), and thus we anticipated it would be permissive to polypeptide insertions (Fig. S12A). This recipient protein was treated with the HA-embedded transposon with flanking complementary split inteins. Gratifyingly, we observed efficient protein transposition, with the embedded splicing apparatus being cleanly removed and replaced by the exogenous insert (Fig. 3B, Fig. S12B). The fluorescence properties of eGFP, which are highly sensitive to the native 3D structure of the protein(*22*), provided a convenient means with which to monitor the folded state of the protein during the reaction. We observed no change in the eGFP emission spectrum during the reaction, which is consistent with the transposition occurring in a concerted manner on the folded protein (Fig. 3C, Fig. S12C).

**Fig 3.**
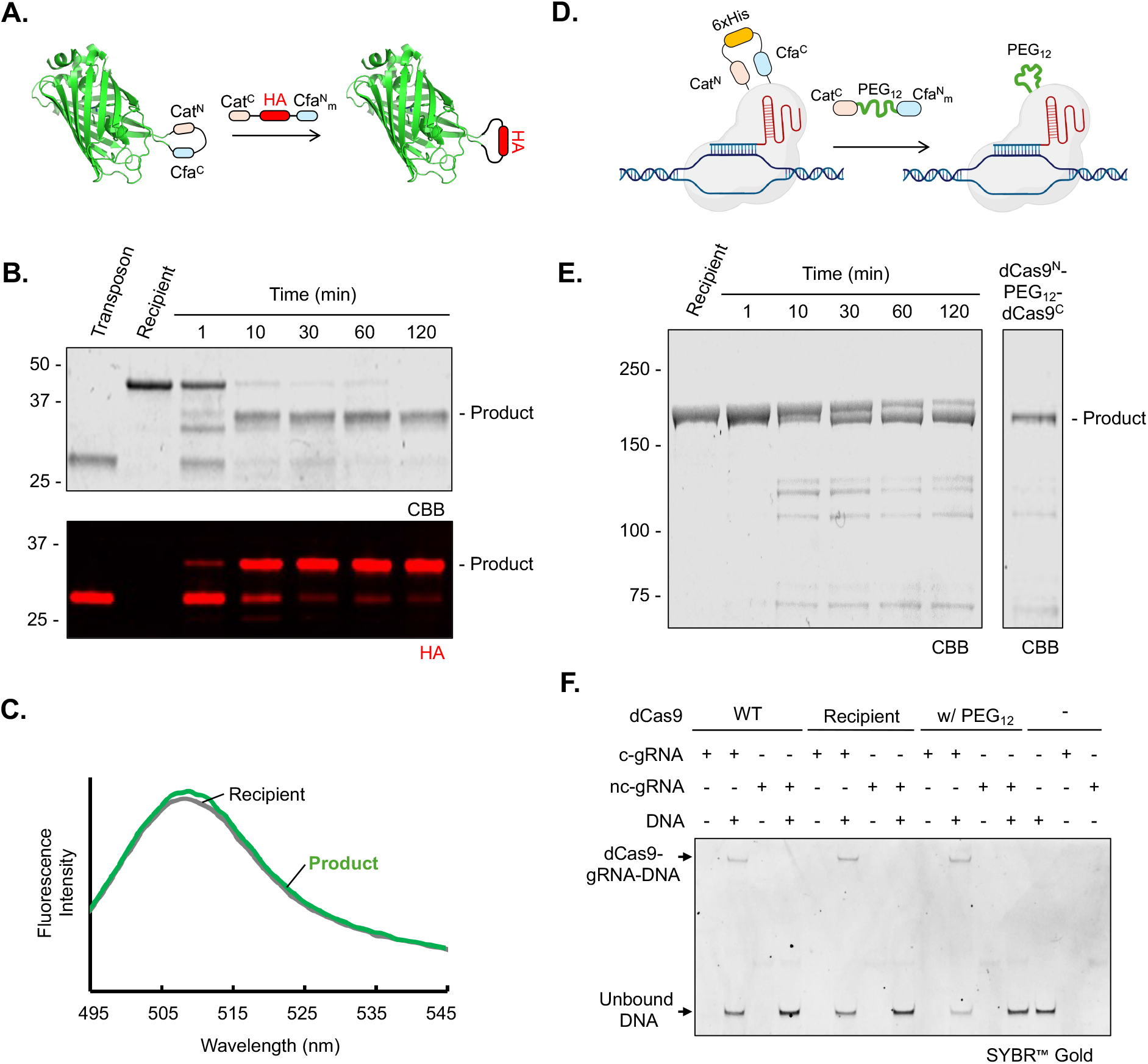
Protein transposition enables traceless installation of protein tags and chimeric cargo on folded proteins. (a) Scheme illustrating the transposition reaction on an internal loop of eGFP. (b) eGFP recipient and HA tag transposon constructs (2 μM each) were reacted for the indicated times in 100 mM phosphate, 150 mM NaCl, 1 mM EDTA, 1 mM TCEP, pH 7.2. Reactions mixtures were analyzed by SDS-PAGE with Coomassie staining (CBB) and western blotting with anti-HA antibody. The expected transposition product is noted in both analyses. (c) The fluorescence emission spectra (λ_ex_ = 488 nm) of the recipient eGFP protein before and after the transposition reaction. (d) Schematic of the transposition reaction within the dCas9 fold using Cat/mut-Cfa orthogonal split intein pair. In the example shown, the 6x His tag inside the recipient dCas9 gets replaced by a PEG_12_ abiotic polymer enabling purification of the product from the unreacted recipient. (e) Analysis of the dCas9 transposition reaction involving a PEG_12_-containing transposon construct. Recipient dCas9 and PEG_12_ transposon constructs (1.5 μM of each) were reacted for the indicated times in 50 mM Tris, 250 mM NaCl, 1 mM TCEP, 10% v/v glycerol, pH 7.5. Reactions mixtures were further purified by reverse nickel affinity chromatography. Both the reaction progress (left) and purified product (right) were analyzed by SDS-PAGE with Coomassie staining and the expected product is indicated. (f) DNA binding activity of dCas9 constructs. A wild-type dCas9 without any transposition construct, the recipient dCas9 and the purified dCas9 containing PEG_12_ after transposition were mixed with DNA in the presence of the complementary guide RNA (c-gRNA) or non-complementary guide RNA (nc-gRNA) and their DNA binding capability characterized by 5% native TBE gel electrophoresis with SYBR Gold staining.

Encouraged by the above results, the transposition system was next applied to a much larger recipient protein. For this, we chose the endonuclease deficient version of *Streptococcus pyogenes* CRISPR associated protein 9 (dCas9), a ∼160 kDa globular protein that has been extensively engineered for genomics applications(*23-25*). Specifically, we examined whether an exogenous cassette could be transposed into a permissive site (residues 573-574 located within a loop) in the middle of the protein (Fig. 3D) (*26*). Accordingly, a chimeric version of the protein was generated in which the splicing apparatus, separated by a His-tag, was embedded at this position (Fig. S13A). Treatment of this recipient dCas9 protein with a reactive transposon construct leads to the efficient generation of the expected transposition product which can be purified using a reverse–nickel affinity chromatography procedure, exploiting the fact that the embedded His-tag is removed during the process (Fig. S13B). This engineered version of dCas9 (recipient dCas9) retained its biological activity as demonstrated by the ability to bind a target DNA sequence in the presence of a cognate guide RNA (Fig. 3F) when compared to its unmodified counterpart. In initial studies, we showed that this recipient dCas9 is capable of undergoing a successful transposition reaction using the transposon containing an HA tag (Fig. S14).

In principle, our approach provides the means to alter both the side chains and amide backbone of a target protein. To illustrate this point, we set about introducing a completely abiotic polymer into the dCas9 protein. Specifically, the dCas9 recipient was reacted with a transposon construct in which a polyethylene glycol (PEG) chain was nested between the Cat^C^ and Cfa^N^_m_ fragments (Fig. 3D, Figs. S15-S16). Notably, this necessitated development of an efficient modular strategy for generation of transposon constructs containing synthetic inserts of choice (Fig. S15). We first validated the incorporation of the PEG chain into our model MBP-eGFP fusion protein by SDS-PAGE and ESI-MS (Fig. S17), which gave us confidence that this abiotic polymer could be inserted into protein backbones. As in the previous examples, the transposition reaction proceeded smoothly, affording a backbone engineered version of dCas9 that, importantly, retained its DNA binding activity in the presence of complementary guide RNA (Figs. 3E-F). The success of this example underscores the remarkable synthetic flexibility granted by our transposition approach. Indeed, to our knowledge, this is the first time a polymer of this type has been introduced into the backbone of an otherwise recombinant protein.

### Protein Transposition Enables Insertion of Functional Probes for the Biochemical Studies of Protein Complexes

Next, we asked whether our system could be used to directly engineer a protein complex. As noted earlier, proteins that reside within multi-subunit complexes represent a significant blind spot for *in vitro* semisynthesis (*1*). Our strategy should help alleviate this limitation by allowing direct manipulation of a suitable engineered pre-formed recipient complex. To test this, we inserted the transposition apparatus into a key regulatory region of SMARCA5, the core ATPase subunit of the ATP-dependent chromatin remodeling complex, ACF (*27-29*). Previous studies from our group identified a central basic domain with the sequence KRERK, termed the acidic patch binding (APB) motif, in this ∼120 kDa protein required for chromatin remodeling activity (*30*). The SMARCA5 recipient protein used in our studies contained the pre-transposition apparatus in place of this basic domain, which ablates APB-dependent functions of SMARCA5 (Fig. 4A). Thus, we predicted that successful reaction with a transposon construct containing this missing regulatory domain would restore the chromatin remodeling activity of the multi-subunit ACF complex, i.e. transposition should act as an ‘on’ switch. Co-incubation of the transposon constructs (Figs. S18-20) with the pre-formed ACF complex harboring the transposition apparatus within the SMARCA5 subunit led to the efficient generation of the expected products, which were confirmed by SDS-PAGE and western blotting (Fig. 4B, Figs. S21-22). Importantly, co-immunoprecipitation studies indicated that the ACF complex remained intact following the transposition reaction (Fig. S22A). We then conducted chromatin remodeling studies using a restriction enzyme accessibility assay (REAA) (*31*). Consistent with the reaction acting as an ‘on’ switch, we observed that transposition restored chromatin remodeling activity to the ACF complex (Fig. 4C).

**Fig 4.**
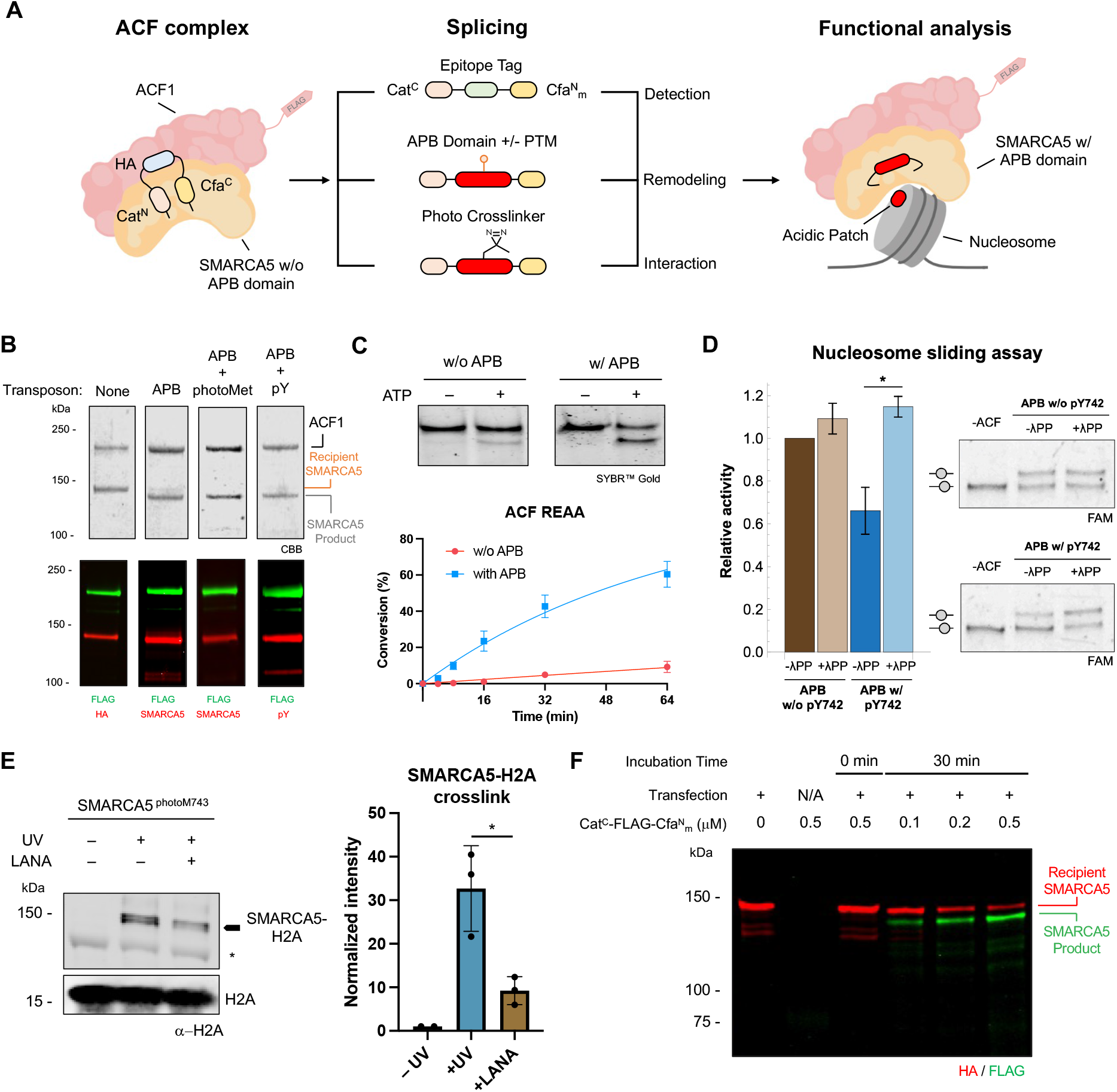
Protein transposition on SMARCA5 and ACF complex enables functional evaluation of remodeling activity and binding preferences. a) Scheme illustrating protein transposition using the Cat/mut-Cfa orthogonal split intein pair on the SMARCA5 subunit of ACF complex; various chemical modifications can be installed via the transposon to enable different biochemical experiments. (b) The pre-made ACF complex containing the recipient SMARCA5 (500 nM) was mixed with a designated transposon construct (500 nM) in 25 mM HEPES, 60 mM KCl, 10 mM MgCl_2_, 1 mM TCEP, 10% v/v glycerol, 0.02% v/v IGEPAL CA630, pH 7.75. Following overnight reaction at 4 °C the products were characterized by SDS-PAGE with Coomassie staining (CBB) and western blotting with indicated antibodies. The APB transposons used in this experiment are as follows: SMARCA5 APB1 (APB), SMARCA5 APB+photoMet (APB w/photoMet), and SMARCA5 APB+pY (APB + pY) (See Fig. S22 for further details) (c) Top: Remodeling activity of modified ACF complexes on mononucleosomes (MNs) as assayed by REAA (10 nM ACF, 10 nM MNs); reintroducing the APB domain into a catalytically inactive ACF complex via protein transposition rescues its activity. Bottom: Kinetics of remodeling for the indicated ACF complexes as measured by REAA, Errors = s.e.m. (*n* = 3 independent experiments). Representative native gel analyses of the remodeling reactions are shown in Fig. S31. (d) Introducing a phosphotyrosine modification into SMARCA5 (pY742) markedly slows nucleosome sliding, as observed by electrophoretic mobility shift after repositioning (8.33 nM ACF, 10 nM MNs, 4 minutes). Upon λ protein phosphatase (λPP) treatment, ACF activity is fully restored. Off-center bead-on-a-string represents the un-remodeled MN whereas the on-center bead-on-a-string represents the remodeled MN. Quantification is by gel densitometry; error bars represent the s.e.m. from 3 replicates. * denotes P value < 0.05. See Fig. S32 for more details. (e) A photo-methionine residue (photoM743) installed within SMARCA5 via protein transposition enables photo-crosslinking between SMARCA5 (10 pmol) and a mononucleosome (10 pmol) upon UV-irradiation (20 min). Left: Immunostaining with an H2A antibody shows a positive crosslink between SMARCA5 and histone, which is significantly ablated upon the addition of excess LANA peptide (10 uM) — a competitive binder to the nucleosome acidic patch. The asterisk denotes nonspecific binding of histone antibody to SMARCA5. Right: densitometry analysis of the crosslinking signal in the immuno-blot normalized to total H2A signal. Errors represent the s.e.m. from 3 independent biological replicates. * denotes P value < 0.05. (f) Immunoblot analysis of *in nucleo* protein transposition reaction between endogenously expressed recipient SMARCA5 in HEK 293T cells containing an embedded HA tag (red signal) and exogenously added transposon construct (0.1–0.5 uM) containing a FLAG tag (green signal) over 30 min. Successful *in nucleo* transposition is denoted by the decrease in the HA signal with a concomitant increase in the FLAG signal.

We next used our transposition technology to study the effect of dynamic PTMs within SMARCA5. Phosphoproteomics analyses of cancer cell lines have identified a phosphorylated tyrosine residue (pTyr742) that is adjacent to the SMARCA5 basic APB motif(*32, 33*). Notably, this tyrosine residue is highly conserved in ISWI orthologs (Fig. S23). While the function of this PTM is not known, its location proximal to this critical domain led us to speculate that it might alter ACF remodeling activity. Indeed, structural modeling using an available cyro-EM structure of ISW1 bound to the nucleosome (*34*) suggests that this phosphotyrosine is positioned to interact with basic sidechains in the APB motif, thereby affecting its function (Fig. S24). To explore the functional impact of this PTM, we used our transposition system to successfully install a peptide transposon containing pTyr742 within the ACF complex (Fig. 4B, Fig. S19). Consistent with our hypothesis, we observed a significant phosphorylation-dependent decrease in ACF remodeling activity as readout by electrophoretic mobility shift assays (EMSA) (Fig. 4D). Notably, this represents the first direct demonstration that a PTM within an ATP-dependent chromatin remodeler affects its biochemical function. As such, this result argues for additional studies on this phospho-PTM in order to fully explore its potential role in tuning ACF remodeling in cells.

SMARCA5 also provided the opportunity to test whether our technology could be used to install biochemical probes into the interior of recipient proteins. Accordingly, we used the transposition system to incorporate a diazirine-containing photo-methionine (photo-Met) residue adjacent to the basic domain of SMARCA5 (Fig. 4A, Fig. S20). A transposon construct containing this crosslinker was generated and then reacted with the ACF recipient to give the expected semisynthetic ACF complex, which retains remodeling activity (Fig. S25). This crosslinker probe was then exploited to gain insight into how the basic domain in SMARCA5 regulates ACF activity. To that end, semisynthetic SMARCA5 was UV irradiated in the presence of a nucleosomal substrate. Immunoblotting of this reaction mixture revealed robust crosslinking to histone H2A (Fig. 4E and Fig. S26), which corroborates our earlier studies that reported an anchoring effect of the basic region of SMARCA5 to the nucleosome acidic patch (*30*). Further supporting the specificity of this interaction, significant ablation of crosslinking between SMARCA5 and histone H2A was observed in the presence of an excess of the latency-associated nuclear antigen (LANA) peptide (Fig. 4E, Fig. S27), which binds to the acidic patch of the nucleosome (*35*) .

Finally, since split inteins do not cross-react with mammalian proteins (*36*), we wondered whether the transposition reaction would work in a more complex cellular environment. To investigate this, the SMARCA5 recipient protein bearing an internal HA tag was expressed in HEK 293T cells and the isolated nuclei from these cells treated with the transposon construct containing a recombinant FLAG tag (Fig. S28). As in the *in vitro* examples, we observed highly efficient transposition as revealed by western blotting (Fig. 4F). Taken together, these experiments with SMARCA5 and the ACF complex highlight the broad applicability of protein transposition from *in vitro* biochemical studies to *in nucleo* experiments.

## Summary

In conclusion, we have developed an *in vitro* protein engineering strategy that allows internal regions of suitably engineered recipient proteins to be replaced with exogenous inserts in a single operation. Our protein transposition approach circumvents the need to generate and manipulate fragments of a target protein during *in vitro* semisynthesis and in the process obviates the need for a final folding step. Notably, the strategy allows direct engineering of protein complexes both *in vitro* and in more complex biological settings such as the cell nucleus, opening the way to a range of *in situ* applications. A key consideration for successful deployment of protein transposition will be identification of permissive sites within the recipient in which to embed the split inteins; clearly, not every location in a target protein will tolerate the insertion of the necessary transposition apparatus. Rather, we imagine that surface exposed loops, linker domains and intrinsically disordered regions (the prevalence and functional importance of which continues to grow (*37*)) are likely to be optimal sites. Given the wealth of protein structure information now available to the community, including recent advances in computational methods (*38-40*), we anticipate that many potential target systems will be identified that fulfill these design criteria. Thus, the current approach is expected to greatly expand the range of proteins accessible to *in vitro* semisynthesis.

## ACKNOWLEDGMENTS

We thank members of the Muir lab for many helpful discussions during the course of this work. We also thank Saw Kyin and Henry H. Shwe at the Princeton Proteomics Facility.

## Funding

This work was funded by NIH-GMS grant R01 GM086868 and NIH-NCI grant R01 CA240768. N.E.S.T. is supported by an NIH postdoctoral fellowship (GM149123). J.A.O. is supported by a Damon Runyon Quantitative Biology Fellowship (DRQ-19-24). X.Y. is supported by a graduate fellowship from the China Scholarship Council (CSC).

## Author contributions

Y.H., N.E.S.T., R.E.T. and T.W.M. conceived the project. All authors contributed to the experimental design and execution. The manuscript was written by Y.H., N.E.S.T. and T.W.M.

## Competing interests

The authors declare that they have no competing interests.

## Data and materials availability

All data needed to evaluate the conclusions in the paper are present in the paper or the supplementary materials.

## Notes

### Competing Interest Statement

The authors have declared no competing interest.

