## Supplementary Information for "Protein Editing using a Concerted Transposition Reaction"

### Table of Contents


Materials and Methods ........ S3
- General Methods ........ S3
- Equipment ........ S4
- Statistics and Reproducibility ........ S4
- Computational Modeling ........ S5

Molecular Cloning Protocols ........ S5

General Protein Expression and Purification Protocols ........ S16
- Expression of intein-containing constructs ........ S16
- Purification of intein-containing constructs ........ S16

Expressed Protein Ligation ........ S22
- General Strategy for Protein Semisynthesis ........ S22
- GyrA Intein Expressed Protein Ligation with MesNa ........ S23
- Split Intein Expressed Protein Ligation with MesNa ........ S24
- Peptide Synthesis ........ S24
- Native Chemical Ligation ........ S25
- Synthesis of LANA peptide ........ S27

Methods for Protein Transposition ........ S28
- General Strategy for Protein Transposition ........ S28
- Mixed Nitrogen Isotope Protein Transposition ........ S28
- Protein Transposition on eGFP and Relevant Fluorescence Measurements ........ S29
- Assembly of the Recipient ACF complex ........ S29
- Protein Transposition on the Recipient ACF Complex ........ S29
- Lambda Phosphatase Experiments for an ACF Complex with SMARCA5 (pY742) ........ S31
- Synthesis of SMARCA5 (photoMet743) by Protein Transposition ........ S31
- Transposition Reactions on Recipient dCas9 ........ S31
- LC-MS/MS Analysis of Transposition Products ........ S32

Methods for dCas9 Experiments ........ S33
- In Vitro Transcription of gRNA ........ S33
- In Vitro dCas9:gRNA:DNA Binding Gel Shift Assay ........ S33

Methods for Nucleosome Remodeling by Designer ACF Complexes ........ S34
- Nucleosome preparation ........ S34
- ACF Remodeling Assay ........ S35

Photo-crosslinking Protocol ........ S36

*In nucleo* Protein Trans-Splicing Protocol ........ S37

Antibodies and Concentrations Used ........ S38

Supplemental Figures ........ S39

Uncropped Gels and Blots Used in this Study ........ S71

Full Ion Mass List for LC-MS/MS Experiments ........ S82

References ........ S83

### Materials and Methods

#### General Materials

Common reagents and chemicals were purchased from Millipore Sigma (St. Louis, MO) unless stated otherwise and were used without further purification. Canonical Fmoc amino acids for use in solid-phase peptide synthesis (SPPS) were purchased from either Matrix Innovations (Quebec City, Canada) or Novabiochem (Darmstadt, Germany). Coupling reagent (7-Azabenzotriazol-1-yloxy)tripyrrolidinophosphonium hexafluorophosphate (PyAOP) was purchased from Oakwood (Estill, SC). Trityl ChemMatrix peptide resin was purchased from Biotage (Charlotte, NC). HO-TCP(Cl)-ProTide peptide resin from purchased from CEM (Charlotte, NC). Trifluoracetic acid (TFA) was purchased from Halocarbon (North Augusta, SC). Oligonucleotides and synthetic genes were purchased from Sigma-Aldrich (Milwaukee, WI). PrimeSTAR® HS DNA Polymerase (premixed) were purchased from Takara Bio USA, Inc. (San Jose, CA). 2x Gibson Assembly Master Mix were purchased from New England Biolabs (Ipswich, MA). "In-house" high-competency cells used for cloning and protein expression were generated from One Shot BL21 (DE3) chemically competent *E. coli* and sub-cloning efficiency DH5α competent cells purchased from Invitrogen (Calsbad, CA). Gibco™ Sf9 cells (Invitrogen, Catalog Number: 12659017) and Gibco™ Sf-900™ III SFM media (Invitrogen, Catalog Number: 12658027) were used for insect-cell based protein expression. DNA purification kits were purchased from Qiagen (Valencia, CA). All plasmids were sequenced by GENEWIZ (South Plainfield, NJ) or Plasmidsaurus (Eugene, OR). Acrylamide, TEMED, APS, and Criterion Empty Cassette were purchased from Bio-Rad. N,N-diisopropylethylamine (DIPEA), Luria Bertani (LB) media, phenylmethylsulfonyl fluoride (PMSF), and all buffering salts were purchased from Fisher Scientific (Pittsburgh, PA). Dimethylformamide (DMF), dichloromethane (DCM), Coomassie brilliant blue, triisopropylsilane (TIS), β-mercaptoethanol (BME), ammonium-$^{15}$N chloride ($^{15}NH_4Cl$), 4-Mercaptophenylacetic acid (MPAA) were purchased from Sigma-Aldrich (Milwaukee, WI) and used without further purification. Sodium 2-mercaptoethanesulfonate (MESNa) was purchased from Combi-Blocks (San Diego, CA). Tris(2-carboxyethyl)phosphine hydrochloride (TCEP) and isopropyl-β-D-thiogalaetopyranoside (IPTG) were purchased from Gold Biotechnology (St. Louis, MO). Nickel-nitrilotriacetic acid (Ni-NTA) resin and Nitrocellulose membrane (0.45 μm) was purchased from

Thermo Scientific (Rockford, IL). Fmoc-NH-$PEG_{12}$-$CH_2CH_2COOH$ was purchased from Ambeed (Arlington Hts, IL). Anti-DYKDDDDK (FLAG) G1 Affinity Resin and the DYKDDDK(FLAG) peptide were purchased from GenScript (Piscataway, NJ). Lambda Protein Phosphatase (Lambda PP) was purchased from New England Biolabs (Ipswich, MA). All reagents for cell culture and transfection were obtained from Thermo Fisher unless otherwise indicated. HEK 293T cells (ATCC) were cultured in DMEM supplemented with 10 % FBS (Sigma-Aldrich), 2 mM L-glutamine, and 500 units/ml penicillin and streptomycin. Transfections were performed using lipofectamine 2000 or lipofectamine 3000 (Invitrogen) as directed by the manufacturer.

**Equipment**

Analytical RP-HPLC was performed on Agilent 1260 Infinity with a Vydac C18 column (5 µm, 4.6 x 150 mm) at a flow rate of 1 mL/min. Semi-preparative RP-HPLC was performed on Agilent 1260 Infinity with a Zorbax 300SB-C18 column (5 µm, 9.4 x 250 mm) at a flow rate of 4 mL/min. Large scale peptide purification was performed with preparative Waters RP-HPLC with a 2535 Quartenary Gradient Module employing an XBridge Peptide BEH C18 OBD Prep Column, 300Å (10 µm, 19 × 250 mm) (Waters) at a flow rate of 20 mL/min. All runs used 0.1 % TFA (trifluoroacetic acid) in water (solvent A) and 90 % acetonitrile in water with 0.1 % TFA (solvent B) as the mobile phases. Unless otherwise stated, peptides and proteins were analyzed using the following gradients: 0% B for 2 minutes followed by 0-75% B over 40 minutes. Electrospray ionization mass spectrometric analysis (ESI-MS) was performed on a Bruker Daltonics MicroTOF-Q II mass spectrometer. Lyophilization of samples after HPLC was performed on a Millrock Technology MD85 lyophilizer (Kingston, NY). Cell disruption was performed with an Ultrasonic Liquid Processor Sonic Dismembrator (ThermoFisher). All size-exclusion chromatography (SEC) was performed on a GE Healthcare (Chicago, IL) AKTA PV-908 FPLC system or a Cytiva (Marlborough, MA) AKTA Go protein purification system with a Superdex S200 increase 10/300 column. SDS-PAGE gels and western blots were imaged on a LI-COR Odyssey Photoimager (Lincoln, NE) or a Cytiva Amersham ImageQuant 800 (Malborough, MA). Fluorescence analysis was performed on a HORIBA Jobin Yvon Fluorolog-3 spectrofluorometer.

**Statistics and Reproducibility**

All statistical analyses were conducted in GraphPad Prism v.10.2.2. P values were determined by unpaired t-test as listed in the figure legends. The statistical significances of differences (ns denotes not significant, * denotes $P < 0.05$, ** denotes $P < 0.01$, ***, denotes $P < 0.001$, **** denotes $P < 0.0001$) are specified throughout the figures and legends. All experiments analyzed by SDS-PAGE/Western blotting were repeated at least three times (independent biological replicates).

**Computational Modeling**

AlphaFold 3 (AF3) was first used to predict of the structure of ISW1a-H2A complex (*1*). With this as a starting point, ISW1a(pTyr775) was then introduced using the Vienna-PTM 2.0 package (http://vienna-ptm.univie.ac.at/) (*2-4*). Next, we mapped this modeled PTM-containing ISW1a-H2A structure onto the full nucleosome using the cryo-EM structure of the ISW1a-nucleosome complex (PDB 6JYL) (*5*) to align the APB region. The modeled structure was visualized using ChimeraX (v.1.7).

#### Molecular Cloning Protocols

All constructs for *E. coli* expression were cloned using pET (Kanamycin/Ampicilin resistance) or pTXB1 (Ampicilin resistance) vectors. Plasmids encoding for SMARCA5 and baculovirus for expressing ACF1-FLAG were obtained as previously described (*6*). Plasmids encoding for Cas9-containing proteins were cloned from pET-Cas9-NLS-6xHis, which was a gift from David Liu (Addgene plasmid #62933; http://n2t.net/addgene:62933; RRID: Addgene_62933). HA-MBP-$IntA^{N}$-linker-$IntB^{C}$-eGFP-FLAG, $IntA^{C}$-transposon-$IntB^{N}$, $IntA^{C}$-linker-GyrA and acidic patch binding region (APB region) were assembled using Gibson assembly from previously described constructs (*7-9*). The TEV protease site, HA and FLAG epitope tags and 6xHis tag within the linker were introduced using inverse PCR with overhang primers followed by blunt-end ligation. Mutations on $Cfa^{N}$ and Cas9 (for the conversion to dCas9) were carried out by site-directed mutagenesis. Insertion of $IntA^{N}$-linker-$IntB^{C}$ sequence into the recipient protein sequence (eGFP, dCas9, SMARCA5) was performed by Gibson assembly. Plasmids were transformed into DH5α competent *E. coli* cells for amplification. All plasmids were purified using mini-prep protocol (Qiagen) and validated by Sanger sequencing (GENEWIZ) or full plasmid sequencing (Plasmidsaurus).

A full list of plasmids with fully annotated amino acid sequences for protein expression in this study (here shown with unprocessed N-terminal methionine and His$_{n=5/6}$-SUMO tag when relevant) is provided below. Final protein products that are obtained after enzymatic cleavage (Ulp1 or TEV protease) are shown in bold for all constructs. Cfa, VidaL and Cat sequences were underlined and shaded.

Plasmid 1:

**pET-5xHis-SUMO-HA-MBP-ESG-Vid$^{N}$-linker-(TEV)-Cfa$^{C}$-CFN-eGFP-FLAG**

MGSSHHHHHGSGLVPRGSASMSDSEVNQEAKPEVKPEVKPETHINLKVSDGSSEIFFKIK
KTTPLRRLMEAFAKRQGKEMDSLRFLYDGIRIQADQTPEDLDMEDNDIIEAHREQIGG**Y**
**PYDVPDYAKIEEGKLVIWINGDKGYNGLAEVGKKFEKDTGIKVTVEHPDKLEEKFP**
**QVAATGDGPDIIFWAHDRFGGYAQSGLLAEITPDKAFQDKLYPFTWDAVRYNGKL**
**IAYPIAVEALSLIYNKDLLPNPPKTWEEIPALDKELKAKGKSALMFNLQEPYFTWPL**
**IAADGGYAFKYENGKYDIKDVGVDNAGAKAGLTFLVDLIKNKHMNADTDYSIAEA**
**AFNKGETAMTINGPWAWSNIDTSKVNYGVTVLPTFKGQPSKPFVGVLSAGINAASP**
**NKELAKEFLENYLLTDEGLEAVNKDKPLGAVALKSYEEELAKDPRIAATMENAQK**
**GEIMPNIPQMSAFWYAVRTAVINAASGRQTVDEAPKDAQTNESGCLPKEAVVQIRL**
**TKKGSESNKRKSNFSENLYFQGSKKKREQSNDIARGVKIISRKSLGTQNVYDIGVEK**
**DHNFLLKNGLVASNCFNMVSKGEELFTGVVPILVELDGDVNGHKFSVSGEGEGDA**
**TYGKLTLKFICTTGKLPVPWPTLVTTLTYGVQCFSRYPDHMKQHDFFKSAMPEGY**
**VQERTIFFKDDGNYKTRAEVKFEGDTLVNRIELKGIDFKEDGNILGHKLEYNYNSH**
**NVYIMADKQKNGIKVNFKIRHNIEDGSVQLADHYQQNTPIGDGPVLLPDNHYLSTQ**
**SALSKDPNEKRDHMVLLEFVTAAGITLGMDELYKDYKDDDDK**

Plasmid 2:

pET-5xHis-SUMO-**Vid$^{C}$-SGK-HA-GAEY-Cfa$^{N}$**

MGSSHHHHHGSGLVPRGSASMSDSEVNQEAKPEVKPEVKPETHINLKVSDGSSEIFFKIK
KTTPLRRLMEAFAKRQGKEMDSLRFLYDGIRIQADQTPEDLDMEDNDIIEAHREQIGG**M**

IEEKKVTVQELRELYLSGEYTIEIDTPDGYQTIGKWFDKGVLSMVRVATATYETVC
AFNHMIQLADNTWVQACELDVGVDIQTAAGIQPVMLVEDTSDAECYDFEVMHPN
HRYYGDGIVSHNSGKYPYDVPDYAGAEYCLSYDTEILTVEYGFLPIGKIVEERIECT
VYTVDKNGFVYTQPIAQWHNRGEQEVFEYCLEDGSIIRATKDHKFMTTDGQMLPI
DEIFERGLDLKQVDGLP

Plasmid 3:

**pET-5xHis-SUMO-Vid$^{C}$-SGK-HA-GAEY-Cfa$^{N}_{m}$**

MGSSHHHHHGSGLVPRGSASMSDSEVNQEAKPEVKPEVKPETHINLKVSDGSSEIFFKIK
KTTPLRRLMEAFAKRQGKEMDSLRFLYDGIRIQADQTPEDLDMEDNDIIEAHREQIGGM
IEEKKVTVQELRELYLSGEYTIEIDTPDGYQTIGKWFDKGVLSMVRVATATYETVC
AFNHMIQLADNTWVQACELDVGVDIQTAAGIQPVMLVEDTSDAECYDFEVMHPN
HRYYGDGIVSHNSGKYPYDVPDYAGAEYCLSYDTEILTVEYGFLPIGKIVEERIECT
VYTVDKNGFVYTQPIAQWHNRGEQEVFEYCLEDGSIIRATKDHKFLTTDGQLLPID
EIFERGLDLKQVDGLP

Plasmid 4:

**pET-5xHis-SUMO-HA-MBP-EFE-Cat$^{N}$-linker-(TEV)-Cfa$^{C}$-CFN-eGFP-FLAG**

MGSSHHHHHGSGLVPRGSASMSDSEVNQEAKPEVKPEVKPETHINLKVSDGSSEIFFKIK
KTTPLRRLMEAFAKRQGKEMDSLRFLYDGIRIQADQTPEDLDMEDNDIIEAHREQIGGY
PYDVPDYAKIEEGKLVIWINGDKGYNGLAEVGKKFEKDTGIKVTVEHPDKLEEKFP
QVAATGDGPDIIFWAHDRFGGYAQSGLLAEITPDKAFQDKLYPFTWDAVRYNGKL
IAYPIAVEALSLIYNKDLLPNPPKTWEEIPALDKELKAKGKSALMFNLQEPYFTWPL
IAADGGYAFKYENGKYDIKDVGVDNAGAKAGLTFLVDLIKNKHMNADTDYSIAEA
AFNKGETAMTINGPWAWSNIDTSKVNYGVTVLPTFKGQPSKPFVGVLSAGINAASP
NKELAKEFLENYLLTDEGLEAVNKDKPLGAVALKSYEEELAKDPRIAATMENAQK
GEIMPNIPQMSAFWYAVRTAVINAASGRQTVDEAPKDAQTNEFECLSGDTMIEILD
DDGIIQKISMEDLYQRLAGTSESNKRKSNFSENLYFQGSKKKREQSNDIARGVKIISR

**KSLGTQNVYDIGVEKDHNFLLKNGLVASNCFNMVSKGEELFTGVVPILVELDGDV NGHKFSVSGEGEGDATYGKLTLKFICTTGKLPVPWPTLVTTLTYGVQCFSRYPDH MKQHDFFKSAMPEGYVQERTIFFKDDGNYKTRAEVKFEGDTLVNRIELKGIDFKE DGNILGHKLEYNYNSHNVYIMADKQKNGIKVNFKIRHNIEDGSVQLADHYQQNTPI GDGPVLLPDNHYLSTQSALSKDPNEKRDHMVLLEFVTAAGITLGMDELYKDYKDD DDK**

Plasmid 5:

**pET-5xHis-SUMO-Cat$^{C}$-CEASGK-HA-GAEY-Cfa$^{N}$**

MGSSHHHHHGSGLVPRGSASMSDSEVNQEAKPEVKPEVKPETHINLKVSDGSSEIFFKIK KTTPLRRLMEAFAKRQGKEMDSLRFLYDGIRIQADQTPEDLDMEDNDIIEAHREQIGG**M FKLNTKNIKVLTPSGFKSFSGIQKVYKPFYHHIIFDDGSEIKCSDNHSFGKDKIKASTI KVGDYLQGKKVLYNEIVEEGIYLYDLLNVGEDNLYYTNGIVSHNCEASGKYPYDVP DYAGAEYCLSYDTEILTVEYGFLPIGKIVEERIECTVYTVDKNGFVYTQPIAQWHNR GEQEVFEYCLEDGSIIRATKDHKFMTTDGQMLPIDEIFERGLDLKQVDGLP**

Plasmid 6:

**pET-5xHis-SUMO-Cat$^{C}$-CEASGK-HA-GAEY-Cfa$^{N}_{m}$**

MGSSHHHHHGSGLVPRGSASMSDSEVNQEAKPEVKPEVKPETHINLKVSDGSSEIFFKIK KTTPLRRLMEAFAKRQGKEMDSLRFLYDGIRIQADQTPEDLDMEDNDIIEAHREQIGG**M FKLNTKNIKVLTPSGFKSFSGIQKVYKPFYHHIIFDDGSEIKCSDNHSFGKDKIKASTI KVGDYLQGKKVLYNEIVEEGIYLYDLLNVGEDNLYYTNGIVSHNCEASGKYPYDVP DYAGAEYCLSYDTEILTVEYGFLPIGKIVEERIECTVYTVDKNGFVYTQPIAQWHNR GEQEVFEYCLEDGSIIRATKDHKFLTTDGQLLPIDEIFERGLDLKQVDGLP**

Plasmid 7:

pET-5xHis-SUMO-**Cat$^{C}$-CEASGK-FLAG-GAEY-Cfa$^{N}_{m}$**

MGSSHHHHHGSGLVPRGSASMSDSEVNQEAKPEVKPEVKPETHINLKVSDGSSEIFFKIK
KTTPLRRLMEAFAKRQGKEMDSLRFLYDGIRIQADQTPEDLDMEDNDIIEAHREQIGG**M**
**FKLNTKNIKVLTPSGFKSFSGIQKVYKPFYHHIIFDDGSEIKCSDNHSFGKDKIKASTI**
**KVGDYLQGKKVLYNEIVEEGIYLYDLLNVGEDNLYYTNGIVSHN**CEASGK**DYKDD**
**DDK**GAEY**CLSYDTEILTVEYGFLPIGKIVEERIECTVYTVDKNGFVYTQPIAQWHNR**
**GEQEVFEYCLEDGSIIRATKDHKFLTTDGQLLPIDEIFERGLDLKQVDGLP**

Plasmid 8:

pET-5xHis-SUMO-**eGFP$^{N}$-GSSGGGFE-Cat$^{N}$-linker-(TEV)-Cfa$^{C}$-CFNGGGGSSGS-eGFP$^{C}$**

MGSSHHHHHGSGLVPRGSASMSDSEVNQEAKPEVKPEVKPETHINLKVSDGSSEIFFKIK
KTTPLRRLMEAFAKRQGKEMDSLRFLYDGIRIQADQTPEDLDMEDNDIIEAHREQIGG**M**
**VSKGEELFTGVVPILVELDGDVNGHKFSVSGEGEGDATYGKLTLKFICTTGKLPVP**
**WPTLVTTLTYGVQCFSRYPDHMKQHDFFKSAMPEGYVQERTIFFKDDGNYKTRAE**
**VKFEGDTLVNRIELKGIDFKEDGNILGHKLEYNYNSHNVYIMADKQKNGIKVNFKI**
**RHNIEDGSSGGGFECLSGDTMIEILDDDGIIQKISMEDLYQRLAGSESNKRKSNFS**EN
LYFQG**SKKKREQSNDIARGVKIISRKSLGTQNVYDIGVEKDHNFLLKNGLVASNCF**
**NGGGGSSGSGSVQLADHYQQNTPIGDGPVLLPDNHYLSTQSALSKDPNEKRDHMV**
**LLEFVTAAGITLGMDELYKDYKDDDDK**

Plasmid 9:

pET-5xHis-SUMO-**Cat$^{C}$-CEASA-GyrA**

MGSSHHHHHGSGLVPRGSASMSDSEVNQEAKPEVKPEVKPETHINLKVSDGSSEIFFKIK
KTTPLRRLMEAFAKRQGKEMDSLRFLYDGIRIQADQTPEDLDMEDNDIIEAHREQIGG**M**
**FKLNTKNIKVLTPSGFKSFSGIQKVYKPFYHHIIFDDGSEIKCSDNHSFGKDKIKASTI**
**KVGDYLQGKKVLYNEIVEEGIYLYDLLNVGEDNLYYTNGIVSHNCEASACITGDAL**
**VALPEGESVRIADIVPGARPNSDNAIDLKVLDRHGNPVLADRLFHSGEHPVYTVRTV**
**EGLRVTGTANHPLLCLVDVAGVPTLLWKLIDEIKPGDYAVIQRSAFSVDCAGFARG**

**KPEFAPTTYTVGVPGLVRFLEAHHRDPDAQAIADELTDGRFYYAKVASVTDAGVQ PVYSLRVDTADHAFITNGFVSHA**

Plasmid 10:

**pET-**5xHis-SUMO-**Cfa$^{N}_{m}$**

MGSSHHHHHGSGLVPRGSASMSDSEVNQEAKPEVKPEVKPETHINLKVSDGSSEIFFKIKKTTPLRRLMEAFAKRQGKEMDSLRFLYDGIRIQADQTPEDLDMEDNDIIEAHREQIGG**CLSYDTEILTVEYGFLPIGKIVEERIECTVYTVDKNGFVYTQPIAQWHNRGEQEVFEYCLEDGSIIRATKDHKFLTTDGQLLPIDEIFERGLDLKQVDGLP**

Plasmid 11:

**pET-dCas9$^{N}$-Cat$^{N}$-6xHis-Cfa$^{C}$-dCas9$^{C}$-NLS-VP64-HA**

**MDKKYSIGLAIGTNSVGWAVITDEYKVPSKKFKVLGNTDRHSIKKNLIGALLFDSGETAEATRLKRTARRRYTRRKNRICYLQEIFSNEMAKVDDSFFHRLEESFLVEEDKKHERHPIFGNIVDEVAYHEKYPTIYHLRKKLVDSTDKADLRLIYLALAHMIKFRGHFLIEGDLNPDNSDVDKLFIQLVQTYNQLFEENPINASGVDAKAILSARLSKSRRLENLIAQLPGEKKNGLFGNLIALSLGLTPNFKSNFDLAEDAKLQLSKDTYDDDLDNLLAQIGDQYADLFLAAKNLSDAILLSDILRVNTEITKAPLSASMIKRYDEHHQDLTLLKALVRQQLPEKYKEIFFDQSKNGYAGYIDGGASQEEFYKFIKPILEKMDGTEELLVKLNREDLLRKQRTFDNGSIPHQIHLGELHAILRRQEDFYPFLKDNREKIEKILTFRIPYYVGPLARGNSRFAWMTRKSEETITPWNFEEVVDKGASAQSFIERMTNFDKNLPNEKVLPKHSLLYEYFTVYNELTKVKYVTEGMRKPAFLSGEQKKAIVDLLFKTNRKVTVKQLKEDYFKKIECLSGDTMIEILDDDGIIQKISMEDLYQRLAGTSESNKRKSNFSHHHHHHSKKKREQSNDIARGVKIISRKSLGTQNVYDIGVEKDHNFLLKNGLVASNCFDSVEISGVEDRFNASLGTYHDLLKIIKDKDFLDNEENEDILEDIVLTLTLFEDREMIEERLKTYAHLFDDKVMKQLKRRRYTGWGRLSRKLINGIRDKQSGKTILDFLKSDGFANRNFMQLIHDDSLTFKEDIQKAQVSGQGDSLHEHIANLAGSPAIKKGILQTVKVVDELVKVMGRHKPENIVIEMARENQTTQKGQKNSRERMKRIEEGIKELGSQILKEHPVENT**

**QLQNEKLYLYYLQNGRDMYVDQELDINRLSDYDVDAIVPQSFLKDDSIDNKVLTRS DKNRGKSDNVPSEEVVKKMKNYWRQLLNAKLITQRKFDNLTKAERGGLSELDKA GFIKRQLVETRQITKHVAQILDSRMNTKYDENDKLIREVKVITLKSKLVSDFRKDF QFYKVREINNYHHAHDAYLNAVVGTALIKKYPKLESEFVYGDYKVYDVRKMIAKS EQEIGKATAKYFFYSNIMNFFKTEITLANGEIRKRPLIETNGETGEIVWDKGRDFAT VRKVLSMPQVNIVKKTEVQTGGFSKESILPKRNSDKLIARKKDWDPKKYGGFDSPT VAYSVLVVAKVEKGKSKKLKSVKELLGITIMERSSFEKNPIDFLEAKGYKEVKKDL IIKLPKYSLFELENGRKRMLASAGELQKGNELALPSKYVNFLYLASHYEKLKGSPE DNEQKQLFVEQHKHYLDEIIEQISEFSKRVILADANLDKVLSAYNKHRDKPIREQAE NIIHLFTLTNLGAPAAFKYFDTTIDRKRYTSTKEVLDATLIHQSITGLYETRIDLSQL GGDPKKKRKVMDKSRASGSGRADALDDFDLDMLGSDALDDFDLDMLGSDALDDF DLDMLGSDALDDFDLDMLINSGSRYPYDVPDYA**

Plasmid 12:

**pET-6xHis- TEV-dCas9-NLS-VP64-HA**

**MGSSHHHHHHGSGENLYFQGDKKYSIGLAIGTNSVGWAVITDEYKVPSKKFKVLG NTDRHSIKKNLIGALLFDSGETAEATRLKRTARRRYTRRKNRICYLQEIFSNEMAK VDDSFFHRLEESFLVEEDKKHERHPIFGNIVDEVAYHEKYPTIYHLRKKLVDSTDKA DLRLIYLALAHMIKFRGHFLIEGDLNPDNSDVDKLFIQLVQTYNQLFEENPINASGV DAKAILSARLSKSRRLENLIAQLPGEKKNGLFGNLIALSLGLTPNFKSNFDLAEDAK LQLSKDTYDDDLDNLLAQIGDQYADLFLAAKNLSDAILLSDILRVNTEITKAPLSAS MIKRYDEHHQDLTLLKALVRQQLPEKYKEIFFDQSKNGYAGYIDGGASQEEFYKFI KPILEKMDGTEELLVKLNREDLLRKQRTFDNGSIPHQIHLGELHAILRRQEDFYPFL KDNREKIEKILTFRIPYYVGPLARGNSRFAWMTRKSEETITPWNFEEVVDKGASAQ SFIERMTNFDKNLPNEKVLPKHSLLYEYFTVYNELTKVKYVTEGMRKPAFLSGEQ KKAIVDLLFKTNRKVTVKQLKEDYFKKIECFDSVEISGVEDRFNASLGTYHDLLKII KDKDFLDNEENEDILEDIVLTLTLFEDREMIEERLKTYAHLFDDKVMKQLKRRRYT GWGRLSRKLINGIRDKQSGKTILDFLKSDGFANRNFMQLIHDDSLTFKEDIQKAQV SGQGDSLHEHIANLAGSPAIKKGILQTVKVVDELVKVMGRHKPENIVIEMARENQT**

**TQKGQKNSRERMKRIEEGIKELGSQILKEHPVENTQLQNEKLYLYYLQNGRDMYV
DQELDINRLSDYDVDAIVPQSFLKDDSIDNKVLTRSDKNRGKSDNVPSEEVVKKMK
NYWRQLLNAKLITQRKFDNLTKAERGGLSELDKAGFIKRQLVETRQITKHVAQIL
DSRMNTKYDENDKLIREVKVITLKSKLVSDFRKDFQFYKVREINNYHHAHDAYLNA
VVGTALIKKYPKLESEFVYGDYKVYDVRKMIAKSEQEIGKATAKYFFYSNIMNFFK
TEITLANGEIRKRPLIETNGETGEIVWDKGRDFATVRKVLSMPQVNIVKKTEVQTG
GFSKESILPKRNSDKLIARKKDWDPKKYGGFDSPTVAYSVLVVAKVEKGKSKKLKS
VKELLGITIMERSSFEKNPIDFLEAKGYKEVKKDLIIKLPKYSLFELENGRKRMLAS
AGELQKGNELALPSKYVNFLYLASHYEKLKGSPEDNEQKQLFVEQHKHYLDEIIEQ
ISEFSKRVILADANLDKVLSAYNKHRDKPIREQAENIIHLFTLTNLGAPAAFKYFDTT
IDRKRYTSTKEVLDATLIHQSITGLYETRIDLSQLGGDPKKKRKVMDKSRASGSGR
ADALDDFDLDMLGSDALDDFDLDMLGSDALDDFDLDMLGSDALDDFDLDMLINSG
SRYPYDVPDYA**

Plasmid 13:

**pET-6xHis-SUMO-SMARCA5$^{N}$(1-715)-Cat$^{N}$-HA-Cfa$^{C}$-C-SMARCA5$^{C}$(747-1051)**

MGSSHHHHHHGSGLVPRGSASMSDSEVNQEAKPEVKPEVKPETHINLKVSDGSSEIFFKI
KKTTPLRRLMEAFAKRQGKEMDSLRFLYDGIRIQADQTPEDLDMEDNDIIEAHREQIGG
**AGSASSAAEPPPPPPPESAPSKPAASIASGGSNSSNKGGPEGVAAQAVASAASAGPAD
AEMEEIFDDASPGKQKEIQEPDPTYEEKMQTDRANRFEYLLKQTELFAHFIQPAAQ
KTPTSPLKMKPGRPRIKKDEKQNLLSVGDYRHRRTEQEEDEELLTESSKATNVCTR
FEDSPSYVKWGKLRDYQVRGLNWLISLYENGINGILADEMGLGKTLQTISLLGYM
KHYRNIPGPHMVLVPKSTLHNWMSEFKRWVPTLRSVCLIGDKEQRAAFVRDVLLP
GEWDVCVTSYEMLIKEKSVFKKFNWRYLVIDEAHRIKNEKSKLSEIVREFKTTNRL
LLTGTPLQNNLHELWSLLNFLLPDVFNSADDFDSWFDTNNCLGDQKLVERLHMVL
RPFLLRRIKADVEKSLPPKKEVKIYVGLSKMQREWYTRILMKDIDILNSAGKMDK
MRLLNILMQLRKCCNHPYLFDGAEPGPPYTTDMHLVTNSGKMVVLDKLLPKLKE
QGSRVLIFSQMTRVLDILEDYCMWRNYEYCRLDGQTPHDERQDSINAYNEPNSTKF
VFMLSTRAGGLGINLATADVVILYDSDWNPQVDLQAMDRAHRIGQTKTVRVFRFI**

**TDNTVEERIVERAEMKLRLDSIVIQQGRLVDQNLNKIGKDEMLQMIRHGATHVFAS
KESEITDEDIDGILERGAKKTAEMNEKLSKMGESSLRNFTMDTESSVYNFECLSGDT
MIEILDDDGIIQKISMEDLYQRLAGEDYREKQKIAFTEWIEPPYPYDVPDYANYAVD
AYVKIISRKSLGTQNVYDIGVEKDHNFLLKNGLVASNCFREALRVSEPKAPKAPRPP
KQPNVQDFQFFPPRLFELLEKEILFYRKTIGYKVPRNPELPNAAQAQKEEQLKIDEA
ESLNDEELEEKEKLLTQGFTNWNKRDFNQFIKANEKWGRDDIENIAREVEGKTPEE
VIEYSAVFWERCNELQDIEKIMAQIERGEARIQRRISIKKALDTKIGRYKAPFHQLRI
SYGTNKGKNYTEEEDRFLICMLHKLGFDKENVYDELRQCIRNSPQFRFDWFLKSR
TAMELQRRCNTLITLIERENMELEEKEKAEKKKRGPKPSTQKRKMDGAPDGRGR
KKKLKL**

Plasmid 14:

**pcDNA3.1(+)-SMARCA5$^{N}$(1-715)-Cat$^{N}$-HA-Cfa$^{C}$-C-SMARCA5$^{C}$(747-1051)**

**SSAAEPPPPPPPESAPSKPAASIASGGSNSSNKGGPEGVAAQAVASAASAGPADAEME
EIFDDASPGKQKEIQEPDPTYEEKMQTDRANRFEYLLKQTELFAHFIQPAAQKTPTS
PLKMKPGRPRIKKDEKQNLLSVGDYRHRRTEQEEDEELLTESSKATNVCTRFEDSP
SYVKWGKLRDYQVRGLNWLISLYENGINGILADEMGLGKTLQTISLLGYMKHYRN
IPGPHMVLVPKSTLHNWMSEFKRWVPTLRSVCLIGDKEQRAAFVRDVLLPGEWDV
CVTSYEMLIKEKSVFKKFNWRYLVIDEAHRIKNEKSKLSEIVREFKTTNRLLLTGTP
LQNNLHELWSLLNFLLPDVFNSADDFDSWFDTNNCLGDQKLVERLHMVLRPFLLR
RIKADVEKSLPPKKEVKIYVGLSKMQREWYTRILMKDIDILNSAGKMDKMRLLNIL
MQLRKCCNHPYLFDGAEPGPPYTTDMHLVTNSGKMVVLDKLLPKLKEQGSRVLIF
SQMTRVLDILEDYCMWRNYEYCRLDGQTPHDERQDSINAYNEPNSTKFVFMLSTR
AGGLGINLATADVVILYDSDWNPQVDLQAMDRAHRIGQTKTVRVFRFITDNTVEE
RIVERAEMKLRLDSIVIQQGRLVDQNLNKIGKDEMLQMIRHGATHVFASKESEITD
EDIDGILERGAKKTAEMNEKLSKMGESSLRNFTMDTESSVYNFECLSGDTMIEILD
DDGIIQKISMEDLYQRLAGEDYREKQKIAFTEWIEPPYPYDVPDYANYAVDAYVKII
SRKSLGTQNVYDIGVEKDHNFLLKNGLVASNCFREALRVSEPKAPKAPRPPKQPNV
QDFQFFPPRLFELLEKEILFYRKTIGYKVPRNPELPNAAQAQKEEQLKIDEAESLND**

**EELEEKEKLLTQGFTNWNKRDFNQFIKANEKWGRDDIENIAREVEGKTPEEVIEYS
AVFWERCNELQDIEKIMAQIERGEARIQRRISIKKALDTKIGRYKAPFHQLRISYGT
NKGKNYTEEEDRFLICMLHKLGFDKENVYDELRQCIRNSPQFRFDWFLKSRTAME
LQRRCNTLITLIERENMELEEKEKAEKKKRGPKPSTQKRKMDGAPDGRGRKKKL
KL**

Plasmid 15:

**pET-5xHis-SUMO-Cat$^{C}$-SMARCA5(716-747, G716C)-Cfa$^{N}_{m}$**

MGSSHHHHHGSGLVPRGSASMSDSEVNQEAKPEVKPEVKPETHINLKVSDGSSEIFFKIK
KTTPLRRLMEAFAKRQGKEMDSLRFLYDGIRIQADQTPEDLDMEDNDIIEAHREQIGG**M
FKLNTKNIKVLTPSGFKSFSGIQKVYKPFYHHIIFDDGSEIKCSDNHSFGKDKIKASTI
KVGDYLQGKKVLYNEIVEEGIYLYDLLNVGEDNLYYTNGIVSHNCEDYREKQKIAF
TEWIEPPKRERKANYAVDAYCLSYDTEILTVEYGFLPIGKIVEERIECTVYTVDKNG
FVYTQPIAQWHNRGEQEVFEYCLEDGSIIRATKDHKFLTTDGQLLPIDEIFERGLDL
KQVDGLP**

Plasmid 16:

**pET-5xHis-SUMO-Cat$^{C}$-SMARCA5(716-747, G716C, A740C)-Cfa$^{N}_{m}$**

MGSSHHHHHGSGLVPRGSASMSDSEVNQEAKPEVKPEVKPETHINLKVSDGSSEIFFKIK
KTTPLRRLMEAFAKRQGKEMDSLRFLYDGIRIQADQTPEDLDMEDNDIIEAHREQIGG**M
FKLNTKNIKVLTPSGFKSFSGIQKVYKPFYHHIIFDDGSEIKCSDNHSFGKDKIKASTI
KVGDYLQGKKVLYNEIVEEGIYLYDLLNVGEDNLYYTNGIVSHNCEDYREKQKIAF
TEWIEPPKRERKCNYAVDAYCLSYDTEILTVEYGFLPIGKIVEERIECTVYTVDKNG
FVYTQPIAQWHNRGEQEVFEYCLEDGSIIRATKDHKFLTTDGQLLPIDEIFERGLDL
KQVDGLP**

Plasmid 17:

**pET-5xHis-SUMO-Cat$^{C}$-SMARCA5(716-739, G716C)-Ava$^{N}$**

MGSSHHHHHGSGLVPRGSASMSDSEVNQEAKPEVKPEVKPETHINLKVSDGSSEIFFKIKKTTPLRRLMEAFAKRQGKEMDSLRFLYDGIRIQADQTPEDLDMEDNDIIEAHREQIGGMFKLNTKNIKVLTPSGFKSFSGIQKVYKPFYHHIIFDDGSEIKCSDNHSFGKDKIKASTIKVGDYLQGKKVLYNEIVEEGIYLYDLLNVGEDNLYYTNGIVSHNCEDYREKQKIAFTEWIEPPKRERKCLSYDTEVLTVEYGFVPIGEIVDKGIECSVFSIDSNGIVYTQPIAQWHHRGKQEVFEYCLEDGSIIKATKDHKFMTQDGKMLPIDEIFEQELDLLQVKGLPE

Plasmid 18 (*10*)

**pTXB1-Npu$^{C}$(N35A)-AAFNSGG-MxeGyrA-6xHis**

**MIKIATRKYLGKQNVYDIGVERDHNFALKNGFIASAAFNSGGCITGDALVALPEGESVRIADIVPGARPNSDNAIDLKVLDRHGNPVLADRLFHSGEHPVYTVRTVEGLRVTGTANHPLLCLVDVAGVPTLLWKLIDEIKPGDYAVIQRSAFSVDCAGFARGKPEFAPTTYTVGVPGLVRFLEAHHRDPDAQAIADELTDGRFYYAKVASVTDAGVQPVYSLRVDTADHAFITNGFVSHAHHHHHH**

Plasmid 19

**pTXB1-Npu$^{C}$(N35A)-AAFNSGG-MBP-6xHis**

**MIKIATRKYLGKQNVYDIGVERDHNFALKNGFIASAAFNSGGKIEEGKLVIWINGDKGYNGLAEVGKKFEKDTGIKVTVEHPDKLEEKFPQVAATGDGPDIIFWAHDRFGGYAQSGLLAEITPDKAFQDKLYPFTWDAVRYNGKLIAYPIAVEALSLIYNKDLLPNPPKTWEEIPALDKELKAKGKSALMFNLQEPYFTWPLIAADGGYAFKYENGKYDIKDVGVDNAGAKAGLTFLVDLIKNKHMNADTDYSIAEAAFNKGETAMTINGPWAWSNIDTSKVNYGVTVLPTFKGQPSKPFVGVLSAGINAASPNKELAKEFLENYLLTDEGLEAVNKDKPLGAVALKSYEEELAKDPRIAATMENAQKGEIMPNIPQMSAFWYAVRTAVINAASGRQTVDEAPKDAQTNHHHHHH**

#### General Protein Expression and Purification Protocols

##### Expression of intein-containing constructs

*E. coli* BL21(DE3) cells were transformed with the appropriate plasmid and grown at 37°C in 1L of Luria-Bertani media (or M9 minimal medium using $^{15}NH_4Cl$ as the only nitrogen source for the expression of $^{15}N$ labeled proteins) containing 50 μg/mL of kanamycin or 100 μg/mL of ampicillin. Once the culture reached an $OD_{600}$=0.6, isopropyl-β-D-thiogalactopyranoside (IPTG) was added to induce expression (final concentration = 0.5 mM). For $Vid^C$-containing proteins, protein expression was carried out for 3 hours at 37°C; for the rest of the constructs, protein expression was carried out for 18 hours at 18°C. The cells were pelleted via centrifugation (4000 rcf, 30 min) and the expressed proteins purified following the procedures listed below. If protein purification was not carried out on the same day, the bacterial cell pellets were flash frozen and stored at -80 °C.

##### Purification of intein-containing constructs

**HA-MBP-$IntA^N$-linker-$IntB^C$-eGFP-FLAG, $Cat^C$-X-$IntB^N$ and $eGFP^N$-GSSGGGFE-$Cat^N$-linker-(TEV)-$Cfa^C$-CFNGGGGSSGS-$eGFP^C$**

Cell pellets containing 5xHis-SUMO-HA-MBP-$IntA^N$-linker-$IntB^C$-eGFP-FLAG, 5xHis-SUMO-$Cat^C$-X-$IntB^N$ (X is a short linker region or truncated peptidic region of the full protein of interest) and 5xHis-SUMO-$eGFP^N$-GSSGGGFE-$Cat^N$-linker-(TEV)-$Cfa^C$-CFNGGGGSSGS-$eGFP^C$ were resuspended in 20 mL cell lysis buffer (100 mM phosphate, 150 mM NaCl, 20 mM imidazole, 1 mM PMSF, 5 mM BME or 1 mM DTT, pH 7.2). The resuspended cells were then lysed by sonication on ice (25% amplitude, 10x 20 sec pulses on / 40 sec off) and the soluble fraction was collected by centrifugation (35,000 rcf, 30 min). The supernatant was then mixed with 4 mL of Ni-NTA resin pre-equilibrated in lysis buffer and incubated at 4°C for 1 hr. The protein-bead mixture was loaded on a fritted column and the resin was washed with 10 CV of lysis buffer. Target proteins were eluted with 15 mL of elution buffer (100 mM phosphate, 150 mM NaCl, 250 mM imidazole, pH 7.2) and then dialyzed against dialysis buffer (100 mM phosphate, 150 mM NaCl, 1 mM TCEP) at 4°C overnight; during dialysis, 6xHis-Ulp1 protease was added into the solution to remove the 5xHis-SUMO tag. After the dialysis, the protein solution was again mixed with 4 mL of Ni-NTA resin and incubated at 4°C for 30 minutes to remove the cleaved 5xHis-SUMO tag and 6xHis-Ulp1 protease. The flow through was collected and concentrated to ~1 mL. The crude protein was then filtered and purified by size exclusion chromatography on a S200 Increase 10/300 gel filtration column with degassed storage buffer (100 mM phosphate, 150 mM NaCl, 1 mM TCEP, 10% v/v glycerol, pH 7.2) as the mobile phase. Fractions were analyzed by SDS-PAGE and fractions containing pure protein were flash-frozen and stored at -80°C (Figs. S2-3, S6, S10, S12, S18, S28).

**Vid$^{C}$-HA-IntB$^{N}$**

Cell pellets containing 5xHis-SUMO-Vid$^{C}$-HA-IntB$^{N}$ were resuspended in 20 mL cell lysis buffer (50 mM Tris, 150 mM NaCl, 1 mM PMSF, 5 mM BME, pH 7.5). The resuspended cells were then lysed by sonication on ice (25% amplitude, 10x 20 sec pulses on / 40 sec off). Lysate was cleared by centrifugation (35,000 rcf, 30 min, 4 ºC) to pellet inclusion bodies. Inclusion bodies were resuspended with 20 mL of extraction buffer (6M guanidine HCl, 50 mM Tris, 20 mM imidazole, 5 mM BME, pH 7.5) and incubated overnight at 4°C. The extracted proteins were collected by centrifugation (35,000 rcf, 30 min) and the supernatant was mixed with 4 mL of Ni-NTA resin pre-equilibrated in extraction buffer and incubated at 4°C for 1 hour. The protein-bead mixture was loaded on a fritted column and the resin was washed with 10 CV of extraction buffer. The target proteins were eluted with 15 mL of elution buffer (6M guanidine HCl, 50 mM Tris, 250 mM imidazole, pH 7.5). Proteins were refolded using a stepwise dialysis method. First, the protein solution was dialyzed against refolding buffer 1 (4M urea, 50 mM Tris, 150 mM NaCl, 1 mM TCEP, pH 7.5) at 4°C for 3 hours, followed by refolding buffer 2 (2M urea, 50 mM Tris, 150 mM NaCl, 200 mM arginine, 1 mM TCEP, 10% v/v glycerol, pH 7.5) 4°C overnight. After one-hour dialysis with refolding buffer 2, 6xHis-Ulp1 protease was added into the solution to remove the 5xHis-SUMO tag. After the dialysis, the protein solution was again mixed with 4 mL of Ni-NTA resin and incubated at 4°C for 30 min to remove the cleaved 5xHis-SUMO tag and 6xHis-Ulp1 protease. The flow through was collected and concentrated to ~1 mL. The crude protein was then filtered and purified by size exclusion chromatography using an S200 increase 10/300 gel filtration column with degassed storage buffer (50 mM Tris, 150 mM NaCl, 200 mM arginine, 1 mM TCEP, 10% v/v glycerol, pH 7.5) as the mobile phase. Fractions were analyzed by SDS-PAGE and fractions containing pure protein were flash-frozen and stored at -80°C (Figs. S3 and S10). Note: $Vid^C$ is prone to aggregation at high concentration, so arginine and glycerol are necessary for the refolding steps.

**$Cat^C$-CEASA-GyrA**

Cell pellets containing 5xHis-SUMO-CEASA-GyrA were resuspended in 20 mL lysis buffer (100 mM phosphate, 150 mM NaCl, 20 mM imidazole, 1 mM PMSF, 1 mM TCEP, pH 7.2). The resuspended cells were then lysed by sonication on ice (25% amplitude, 10x 20 sec pulses on / 40 sec off). The soluble fraction was collected by centrifugation (35,000 rcf, 30 min). The supernatant was then mixed with 4 mL of pre-equilibrated (in lysis buffer) Ni-NTA resin and incubated at 4°C for 1 hr. The protein-bead mixture was loaded on a fritted column and the resin was washed with 10 CV of lysis buffer. The target proteins were eluted with 15 mL of elution buffer (100 mM phosphate, 150 mM NaCl, 250 mM imidazole, pH 7.2) and then dialyzed against dialysis buffer (100 mM phosphate, 150 mM NaCl, 1 mM TCEP) at 4°C overnight. Prior to dialysis, 6xHis-Ulp1 protease was added into the protein solution to remove the 5xHis-SUMO tag. Upon completion of overnight dialysis, this solution was then flash frozen and kept at –80 °C until further use (see EPL section, Fig. S16).

**$Cat^C$-CEDYREKQKIAFTEWIEPPKRERK-$Ava^N$**

Cell pellets containing 5xHis-SUMO-$Cat^C$- CEDYREKQKIAFTEWIEPPKRERK-$Ava^N$ were resuspended in 20 mL lysis buffer (100 mM phosphate, 150 mM NaCl, 20 mM imidazole, 1 mM PMSF, 1 mM TCEP, pH 7.2). The resuspended cells were then lysed by sonication on ice (25% amplitude, 10x 20 sec pulses on / 40 sec off). The soluble fraction was collected by centrifugation (35,000 rcf, 30 min). The supernatant was then mixed with 4 mL of pre-equilibrated (in lysis buffer) Ni-NTA resin and incubated at 4°C for 1 hr. The protein-bead mixture was loaded on a fritted column and the resin was washed with 10 CV of lysis buffer. The target proteins were eluted with 15 mL of elution buffer (100 mM phosphate, 150 mM NaCl, 250 mM imidazole, pH 7.2) and then dialyzed against dialysis buffer (100 mM phosphate, 150 mM NaCl, 1 mM TCEP) at 4°C for 3 h. Prior to dialysis, 6xHis-Ulp1 protease was added into protein solution to remove the 5xHis-

SUMO tag. Upon completion of dialysis, the solution was transferred to a 50 mL falcon tube and urea added to the solution to reach a concentration of 4 M. This solution was then flash frozen and kept at –80 °C until further use (see split intein EPL section, Figs. S19 and S20).

**Npu$^{C}$-AAFNSGGC-OMe**

Cell pellets containing Npu$^{C}$(N35A)-AAFNSGG-MxeGyrA-6xHis were resuspended in 20 mL lysis buffer (100 mM phosphate, 150 mM NaCl, 20 mM imidazole, 1 mM PMSF, 1 mM TCEP, pH 7.2). The resuspended cells were then lysed by sonication on ice (25% amplitude, 10x 20 sec pulses on / 40 sec off). The soluble fraction was collected by centrifugation (35,000 rcf, 30 min). The supernatant was then mixed with 4 mL of pre-equilibrated (in lysis buffer) Ni-NTA resin and incubated at 4°C for 1 hr. The protein-bead mixture was loaded on a fritted column and the resin was washed with 10 CV of lysis buffer. The target proteins were eluted with 15 mL of elution buffer (100 mM phosphate, 150 mM NaCl, 250 mM imidazole, pH 7.2) and then dialyzed against dialysis buffer (100 mM phosphate, 150 mM NaCl, 1 mM TCEP) at 4°C for 3 h. Upon completion of dialysis, the solution was transferred to a 50 mL falcon tube and *in situ* intein cleavage was performed by the addition of H-Cys-OMe (100 mM), TCEP-HCl (10 mM), and adjusting pH to 7.5. The reaction was carried out at r.t. for 24 h and at the end of the reaction, additional TCEP-HCl was added to quench nonspecific disulfides. This solution was filtered and purified by semi-preparative RP-HPLC (0% B for 2 minutes, then 0-73% B over 40 minutes). The fractions were analyzed by analytical RP-HPLC and ESI-MS, and pure fractions containing either the title product or the methyl ester hydrolysis product were pooled and lyophilized (Fig. S34). The lyophilized products were stored at -80°C until use for split intein EPL.

**Npu$^{C}$(N35A)-MBP-His$_6$**

Cell pellets containing Npu$^{C}$(N35A)-AAFNSGG-MBP-6xHis were resuspended in 20 mL lysis buffer (100 mM phosphate, 150 mM NaCl, 20 mM imidazole, 1 mM PMSF, 1 mM TCEP, pH 7.2). The resuspended cells were then lysed by sonication on ice (25% amplitude, 10x 20 sec pulses on / 40 sec off). The soluble fraction was collected by centrifugation (35,000 rcf, 30 min). The supernatant was then mixed with 4 mL of pre-equilibrated (in lysis buffer) Ni-NTA resin and incubated at 4°C for 1 hr. The protein-bead mixture was loaded on a fritted column and the resin was washed with 10 CV of lysis buffer. The target proteins were eluted with 15 mL of elution buffer (100 mM phosphate, 150 mM NaCl, 250 mM imidazole, pH 7.2) and then dialyzed against dialysis buffer (100 mM phosphate, 150 mM NaCl, 1 mM TCEP) at 4°C for 3 h. Upon completion of dialysis, the solution was transferred to a 50 mL falcon tube, flash frozen, and stored at -80°C until use for split intein EPL (Fig. S34).

**$Cfa^{N}_{m}$ ($Cfa^{C}$ M75L M81L)**

Cell pellets containing 5xHis-SUMO-$Cfa^{N}_{m}$ were resuspended in 20 mL cell lysis buffer (50 mM Tris, 150 mM NaCl, 20 mM imidazole, 1 mM PMSF, 5 mM BME, pH 7.5). The resuspended cells were then lysed by sonication on ice (25% amplitude, 10x 20 sec pulses on / 40 sec off). The soluble fraction was collected by centrifugation (35,000 rcf, 30 min). The supernatant was then mixed with 4 mL of pre-equilibrated Ni-NTA resin and incubated at 4°C for 1 hr. The mixture was loaded on a fritted column and the resin was washed with 10 CV of lysis buffer. The target proteins were eluted with 15 mL of elution buffer (50 mM Tris, 150 mM NaCl, 250 mM imidazole, pH 7.5) and then dialyzed against dialysis buffer (50 mM Tris, 150 mM NaCl, 1 mM TCEP) at 4°C overnight; during dialysis, 6xHis-Ulp1 protease was added into the solution to remove the SUMO tag and expose the N-terminal cysteine. The solution was then diluted with HPLC Solvent A and the pH adjusted to 2.0. This solution containing SUMO tag and $Cfa^{N}_{m}$ was filtered and purified by semi-preparative RP-HPLC (100% Solvent A for 5 min, then 30% Solvent B to 60% Solvent B for 40 min). The fractions were analyzed by analytical RP-HPLC and ESI-MS. Pure $Cfa^{N}_{m}$ was pooled and lyophilized (Fig. S34). The lyophilized products were stored at -80°C.

**dCas9 fusion proteins**

Cell pellets containing either $dCas9^{N}$-$Cat^{N}$-6xHis-$Cfa^{C}$-$dCas9^{C}$-NLS-VP64-HA or 6xHis-TEV-dCas9-NLS-VP64-HA were resuspended in 20 mL cell lysis buffer (50 mM Tris, 1M NaCl, 20 mM imidazole, 1 mM PMSF, 5 mM BME, pH 8.0) and then lysed by sonication on ice (25% amplitude, 10x 20 sec pulses on / 40 sec off). The soluble fraction was collected by centrifugation (35,000 rcf, 30 min). The supernatant was then mixed with 4 mL of pre-equilibrated Ni-NTA resin and incubated at 4°C for 1 hr. The mixture was loaded on a fritted column and the resin was washed with 10 CV of lysis buffer. The target proteins were eluted with 15 mL of elution buffer (50 mM Tris, 500 mM NaCl, 250 mM imidazole, pH 8.0) and then dialyzed against dialysis buffer (50 mM Tris, 250 mM NaCl, 1 mM TCEP, pH 7.5) at 4°C overnight; The dialyzed proteins were concentrated to ~1 mL and then filtered and purified by size exclusion chromatography using an S200 increase 10/300 gel filtration column with degassed storage buffer (50 mM Tris, 250 mM NaCl, 1 mM TCEP, 10% v/v glycerol, pH 7.5). Fractions were analyzed by SDS-PAGE. Pure protein was flash-frozen and stored at -80°C. Note: for the dCas9-fusion expressed from Plasmid 12, 500 nM TEV protease was added during dialysis to remove the 6xHis tag (Fig. S13).

**Purification of ACF1-FLAG**

ACF1-FLAG was expressed in Sf9 cells using a baculovirus expression system as described previously (*6, 11*). The Sf9 cells were infected with baculovirus for 48 hours and the cells were harvested via centrifugation (2000 rcf, 15 min). The cell pellet was first washed with PBS buffer and then pelleted via centrifugation (2000 rcf, 15 min). Then the cells were resuspended with 20 mL lysis buffer (25 mM HEPES, 400 mM NaCl, 2 mM EDTA, 2 mM EGTA, 5 mM BME, 1 mM PMSF, 10% v/v glycerol, pH 8.0) and then lysed by sonication on ice (25% amplitude, 10x 20 sec pulses on / 40 sec off) and the soluble fraction was collected by centrifugation (35,000 rcf, 1 hr). The supernatant was then mixed with 1 mL of anti-FLAG G1 resin pre-equilibrated in lysis buffer and incubated at 4°C for 1 hr. The resin was then washed with 10 mL of lysis buffer twice. The ACF1-FLAG was eluted by mixing the resin with 500 μL of 0.25 mg/mL FLAG peptide dissolved in lysis buffer and the flow through was separated from the resin by centrifugation (2000 rcf, 5 min). The elution step was repeated 4 times, and all flow through fractions were pulled together and then concentrated down to ~1 mL. The ACF1-FLAG was further purified by size exclusion chromatography using an S200 increase 10/300 gel filtration column with degassed ACF storage buffer (25 mM HEPES, 300 mM NaCl, 1 mM TCEP, 10% v/v glycerol, pH 7.5) as the mobile phase. Pure protein fractions were flash-frozen and stored at -80°C (Figs. S21 and S22).

**Purification of SMARCA5-containing proteins**

$SMARCA5^N$ (1-715)-$Cat^N$-HA-$Cfa^C$-C-$SMARCA5^C$ (747-1051) [SMARCA5-$HA_{ins}$] protein was expressed in *E. coli* BL21(DE3) following our previously published procedure (*11*). Cell pellets containing 6xHis-SUMO-$SMARCA5^N$ (1-715)-$Cat^N$-HA-$Cfa^C$-C-$SMARCA5^C$ (747-1051) were resuspended in 20 mL cell lysis buffer (25 mM HEPES, 300 mM NaCl, 20 mM imidazole, 5 mM BME, 1 mM PMSF, pH 8.0). The resuspended cells were then lysed by sonication on ice (25% amplitude, 10x 20 sec pulses on / 40 sec off) and the soluble fraction was collected by centrifugation (35,000 rcf, 30 min). The supernatant was then mixed with 4 mL of Ni-NTA resin pre-equilibrated in lysis buffer and incubated at 4°C for 1 hr. The protein-bead mixture was loaded on a fritted column and the resin was washed with 10 CV of lysis buffer. Additional washing steps were done with 10 CV of wash buffer 1 (25 mM HEPES, 500 mM NaCl, 20 mM imidazole, 5 mM BME, 1 mM PMSF, pH 8.0), 10 CV of wash buffer 2 (25 mM HEPES, 1 M NaCl, 20 mM imidazole, 5 mM BME, 1 mM PMSF, pH 8.0), 10 CV of wash buffer 1 again and then 10 CV of lysis buffer. Target proteins were eluted with 15 mL of elution buffer (25 mM HEPES, 300 mM NaCl, 400 mM imidazole, 5 mM BME, 1 mM PMSF, pH 7.6) and then dialyzed against dialysis buffer (25 mM HEPES, 300 mM NaCl, 1 mM TCEP, pH 7.6) at 4°C overnight; prior to dialysis, 6xHis-Ulp1 protease was added into the solution to remove the 6xHis-SUMO tag. After the dialysis, the protein solution was concentrated to ~1 mL. The crude protein was then filtered and purified by size exclusion chromatography on a S200 Increase 10/300 gel filtration column with degassed storage buffer (25 mM HEPES, 300 mM NaCl, 1 mM TCEP, 10% v/v glycerol, pH 7.2) as the mobile phase. Fractions containing pure protein were flash-frozen and stored at -80°C (Figs. S21 and S22).

#### Expressed Protein Ligation

**General Strategy for Protein Semisynthesis**

The semisynthesis of the split intein transposon constructs ($Cat^C$-CEASA-$PEG_{12}$-AEY-$Cfa^N{}_m$, $Cat^C$-SMARCA5(716-747, G716C, A740C, A743photoMet)-$Cfa^N{}_m$, and $Cat^C$-SMARCA5(716-747, G716C, A740C, pY742)-$Cfa^N{}_m$) was carried out by assembling $Cat^C$-X-thioester, synthetic peptide and $Cfa^{N}_{m}$ fragments through a two-step expressed protein ligation (EPL) procedure (Fig. S15), where X is a short linker region or truncated peptidic region of the full protein of interest. In brief, the $Cat^{C}$-X-thioester was formed either by thiolysis of the GyrA intein or by thiolysis of the $Ava^{N}$ intein by sEPL *(vide supra)*. This fragment is then ligated to a synthetic peptide containing a C-terminal hydrazide and subsequently purified by preparative RP-HPLC. The corresponding polypeptide is then subjected to hydrazide oxidation conditions (*12*), followed by trapping of the peptide acyl azide with excess MesNa. This intermediate can either be purified by preparative RP-HPLC or carried forward to the next EPL step after spin filtration to remove excess MesNa and to concentrate the reaction mixture. Chemical ligation of this Mes-thioester intermediate with recombinant $Cfa^{N}_{m}$ yielded the desired product, which was then purified by semi preparative RP-HPLC purification. The lyophilized product was then redissolved in reaction buffer containing 6 M guanidine hydrochloride and dialyzed in a stepwise fashion into reaction buffer (50 mM sodium phosphate, 150 mM NaCl, 1 mM TCEP, 10% v/v glycerol, pH 7.2). The refolded protein was then further purified by size exclusion chromatography [S200 increase 10/300 gel filtration column with degassed ACF storage buffer (50 mM sodium phosphate, 150 mM NaCl, 1 mM TCEP, 10% v/v glycerol, pH 7.2)] before use in further applications.

**GyrA Intein Expressed Protein Ligation with MesNa**

**$Cat^{C}$-CEASA-Mes ($Cat^{C}$-CEASA-$SCH_2CH_2SO_3^{-}Na^{+}$)**

Upon completion of overnight dialysis of $Cat^{C}$-CEASA-GyrA, solid MesNa (100 mM final concentration) and TCEP (10 mM final concentration) were directly added into the protein solution and the pH was adjusted to 7.0. The solution was incubated overnight at room temperature to generate the Mes-thioester protein via GyrA thiolysis. The formation of the Mes-thioester product was monitored by analytical RP-HPLC (100% Solvent A for 2 min, then 20% Solvent B to 75% Solvent B for 40 min). The crude product solution was then treated with an additional 1 mM TCEP (to reduce disulfides), diluted with HPLC Solvent A and equilibrated to pH 2.0. This solution containing cleaved His-SUMO tag, $Cat^{C}$-CEASA-Mes and GyrA was filtered and purified by semi-preparative RP-HPLC (100% Solvent A for 5 min, then 30% Solvent B to 60% Solvent B for 40 min). The purified fractions were checked by analytical RP-HPLC and ESI-MS, with pure

$Cat^{C}$-CEASA-Mes product pooled and lyophilized. The lyophilized products were stored at -80°C until further use (see EPL section).

**Split Intein Expressed Protein Ligation with MesNa**

Crude $Cat^{C}$-CEDYREKQKIAFTEWIEPPKRERK-$Ava^{N}$ was dissolved in phosphate buffer containing 4 M urea (100 mM phosphate, 150 mM NaCl, 1 mM EDTA, 20 mM imidazole, 1 mM TCEP. The streamlined EPL reaction was initiated by addition of 1.5 equiv of $Npu^{C}$-AAFNSGGC-OMe peptide, 200 mM MesNa (~200-300 mg) and 50 mM TCEP and adjusting the pH to 7.2. The reaction was incubated at room temperature overnight and conversion to $Cat^{C}$-linker-Mes thioester was determined by RP-HPLC. This solution was then diluted with HPLC Solvent A and pH adjusted to 2.0 before being filtered and purified by semi-preparative RP-HPLC (100% Solvent A for 5 min, then 30% Solvent B to 60% Solvent B for 40 min). The fractions were analyzed by analytical RP-HPLC and ESI-MS. Pure $Cat^{C}$-linker-Mes fractions were pooled and lyophilized. The lyophilized products were stored at -80°C. Split intein fusion $Npu^{C}$-AAFNSGGC-OMe can be substituted for $Npu^{C}$-AAFNSGG-MBP

**Peptide Synthesis**

Synthetic peptides were synthesized using standard Fmoc-based solid phase peptide synthesis as C-terminal hydrazides and with Boc-Cys(Trt)-OH for the N-terminal Cys residue:

Peptide 1: $H_2N$–C-$PEG_{12}$-AEY–$NHNH_2$

Peptide 2: $H_2N$–CNYM(photo)VDAY–$NHNH_2$

Peptide 3: $H_2N$–CNpYAVDAY–$NHNH_2$

Peptide 1 (0.1 mmol) was synthesized manually on ChemMatrix Trityl-OH resin, preloaded with a hydrazine linker. The solid phase synthesis cycle included: i) Fmoc deprotection with 20% piperidine in DMF (20 min) and ii) coupling of 5 equiv of each amino acid in DMF with 5 equiv of (7-Azabenzotriazol-1-yloxy)tripyrrolidinophosphonium hexafluorophosphate (PyAOP) and 10 equiv diisopropylethylamine (DIPEA) for 20 min. The coupling step was repeated twice for all amino acids. Finally, peptide cleavage and side-chain deprotection were carried out with a cleavage cocktail (92.5% TFA, 2.5% TIPS, 2.5% $H_2O$, 2.5% ethanedithiol) for 2 hours. Cold diethyl ether was added, and the peptides were precipitated and collected by centrifugation at 4000g for 10 minutes. The peptide was dissolved in HPLC solvent A, purified by preparative RP-HPLC (0%B for 2 minutes, then 0-73%B over 40 minutes) and lyophilized.

Peptides 2-3 (0.1 mmol) were synthesized manually on HO-TCP(Cl)-ProTide resin subjected to stepwise chlorination and Fmoc-hydrazine loading. In brief, HO-TCP(Cl)-ProTide resin was treated with thionyl chloride in DCM overnight at rt with nitrogen bubbling. After thorough washing with DCM and DMF, Fmoc-hydrazine (5 equiv) and DIEA (10 equiv) in DMF was added to the resin for 2 x 2 h treatments. The solid phase synthesis cycle included: i) Fmoc deprotection with 20% piperidine in DMF containing 0.1M 1-Hydroxybenzotriazole hydrate (HOBt) (20 min) and ii) coupling of 5 equiv of each amino acid in DMF with 5 equiv of hexafluorophosphate azabenzotriazole tetramethyl uronium (HATU) and 10 equiv DIPEA for 30 min. The coupling step was repeated twice for all amino acids. For non-proteogenic amino acids, the coupling step was performed with 3 equiv of each amino acid in DMF with 3 equiv of PyAOP and 10 equiv DIPEA for 3 h (if coupling step performed once) or 2 x 2 h (if double couplings were carried out). Finally, peptide cleavage and side-chain deprotection were carried out with a cleavage cocktail (92.5% TFA, 2.5% TIPS, 2.5% $H_2O$, 2.5% ethanedithiol) for 3 hours. Cold diethyl ether was added, and the peptides were precipitated and collected by centrifugation at 4000g for 10 minutes. The peptide was dissolved in HPLC solvent A, purified by preparative RP-HPLC (0%B for 2 minutes, then 0-73%B over 40 minutes) and lyophilized.

**Native Chemical Ligation**

*NCL Step 1*

The first NCL reaction step was performed using the following pairs:

1) $Cat^{C}$-CEASA Mes-thioester and H-C-$PEG_{12}$-AEY-$NHNH_2$
2) $Cat^{C}$-CEDYREKQKIAFTEWIEPPKRERK Mes-thioester and H–CNYM(photo)VDAY–$NHNH_2$
3) $Cat^{C}$-CEDYREKQKIAFTEWIEPPKRERK Mes thioester and H–CNpYAVDAY–$NHNH_2$

In all cases, the Mes-thioester fragment (0.5 mM) and synthetic peptide fragment (1 mM) were mixed at room temperature in ligation buffer (6M guanidine HCl, 100 mM sodium phosphate buffer, pH 7.0, 50 mM TCEP, 100 mM MPAA) and allowed to react overnight. The reaction mixtures were then diluted with HPLC Solvent A, acidified to pH 2 and purified by semi-preparative RP-HPLC (0% B for 5 minutes, then 30-55% B over 40 minutes). The pure products were identified by ESI-MS, collected and lyophilized.

*Acyl hydrazide oxidation and thiolysis*

The products from the first NCL reaction were then converted into the corresponding Mes-thioesters using a modified hydrazide oxidation procedure (*12*). Briefly, the fully lyophilized peptide hydrazides were first dissolved in 1 mL of conversion buffer (6M guanidine HCl, 100 mM sodium phosphate buffer, pH 3.0). 10 equiv of $NaNO_2$ was added into the pre-cooled solution at -20°C, which was then incubated at -20 °C for 20 minutes. Solid MesNa was then added into the solution to reach a final concentration of 100 mM, and the solution was brought to room temperature and the pH was slowly adjusted to 7.0. After incubation at pH 7.0 and room temperature for 45 minutes, 50 mM TCEP was added to remove the oxidative byproducts. Finally, the reaction mixtures were diluted with HPLC Solvent A, acidified to pH 2 and purified by semi-preparative RP-HPLC (0% B for 5 minutes, then 30-55% B over 40 minutes). The pure products were identified by ESI-MS, collected and lyophilized.

Alternatively, the reaction mixture can be carried to the next EPL step by spin filtration (3000 MWCO) with concurrent replenishment of ligation buffer (6M guanidine HCl, 100 mM sodium phosphate buffer, pH 7.0, 50 mM TCEP). The reaction mixture is finally concentrated to 0.25 mM before proceeding to the next ligation step.

*NCL Step 2*

The second NCL reaction was performed between $Cfa^{N}_{m}$ and the following semisynthetic constructs.

1) $Cat^{C}$-CEASAC-$PEG_{12}$-AEY-Mes
2) $Cat^{C}$-CEDYREKQKIAFTEWIEPPKRERKCNYM(photo)VDAY-Mes
3) $Cat^{C}$-CEDYREKQKIAFTEWIEPPKRERKCNpYAVDAY-Mes

In all cases, the Mes-thioester fragment (0.25 mM) and $Cfa^{N}_{m}$ (0.5 mM) were mixed at room temperature in ligation buffer (6M guanidine HCl, 100 mM sodium phosphate buffer, pH 7.0, 50 mM TCEP, 100 mM MPAA) and allowed to react overnight. The reaction mixtures were then diluted with HPLC Solvent A, acidified to pH 2 and purified by semi-preparative RP-HPLC (0% B for 5 minutes, then 30-55% B over 40 minutes). The products were identified by ESI-MS, collected and lyophilized. The lyophilized peptides were dissolved in denaturing buffer (6M guanidine HCl, 100 mM sodium phosphate buffer, pH 7.2, 1 mM TCEP) and then dialyzed in a stepwise fashion into reaction buffer (50 mM sodium phosphate, 150 mM NaCl, 1 mM TCEP, pH 7.2). The crude proteins were then further purified by size exclusion chromatography using an S200 increase 10/300 gel filtration column with degassed storage buffer (100 mM sodium phosphate, 150 mM NaCl, 1mM EDTA, 1 mM TCEP, 10% v/v glycerol, pH 7.2) as the mobile phase. Fractions were analyzed by SDS-PAGE. Pure protein was flash-frozen and stored at -80°C.

**Synthesis of LANA peptide (*6*)**

The LANA peptide corresponding to residues 2–22 with a norleucine at position 6 substituted for methionine was synthesized on a CEM Discover Microwave Peptide Synthesizer using Fmoc chemistry and a ChemMatrix Rink amide resin (0.47 mmol/g). Following chain assembly, peptides were cleaved from the resin using 95% TFA, 2.5% TIS, and 2.5% water and purified by C-18 RP-HPLC (0%B for 2 minutes, then 0-73%B over 40 minutes) and lyophilized (Fig. S27). Final products were characterized by analytical C-18 RP-HPLC and ESI-MS.

#### Methods for Protein Transposition

##### General Strategy for Protein Transposition

All protein transposition reactions were performed at room temperature unless otherwise indicated. Briefly, the recipient proteins and the transposon proteins were individually pre-incubated in reaction buffer (100 mM phosphate, 150 mM NaCl, 1 mM EDTA, 1 mM TCEP, pH 7.2 unless otherwise indicated,) at equal concentration for 15 minutes on ice. The reaction was initiated by mixing equal volumes of recipient and transposon proteins. For analysis using SDS-PAGE, the reactions were quenched at indicated times by adding SDS loading buffer. When the recipient protein was smaller than 75 kDa, the SDS-PAGE was run on a 10% bis-tris gel at 160 V for 45 min, using MES-SDS as the running buffer. For immuno-blotting, the gel containing the reaction mixture were transferred to a nitrocellulose membrane (0.7 A, 100 min), which was then blocked with 3% w/v BSA in TBST (20 mM Tris-Cl, pH 7.6, 150 mM NaCl, 0.1% v/v Tween-20) for 30 mins at room temperature. Next, membranes were incubated with the indicated primary antibodies (diluted in TBST containing 1% BSA) at 4°C overnight. After washing three times with TBST, the appropriate Li-Cor IRDye secondary antibodies (1:10000 dilution in TBST) were applied for 1 h at room temperature. Following incubation with the dye-labeled secondary antibodies, membranes were washed three times with TBST and imaged with Li-Cor Odyssey Infrared Imaging System. For RP-HPLC and ESI-MS analysis, the reaction mixtures were diluted with an equal volume of HPLC Solvent A prior to analysis.

##### Mixed Nitrogen Isotope Protein Transposition

$^{14}$N labeled and $^{15}$N labeled recipient proteins were mixed and diluted to reach 2 μM each with reaction buffer (100 mM phosphate, 150 mM NaCl, 1 mM EDTA, 1 mM TCEP, pH 7.2). The reaction was initiated by mixing equal volumes of mixed recipient proteins and transposon protein to reach the final concentration of 2 μM. After 2 hours, the reaction mixture was diluted with equal volumes of HPLC Solvent A and the transposition product isolated by analytical RP-HPLC for ESI-MS analysis. For experiments where cleavage of the TEV site is required, a final concentration of 500 nM TEV protease was added into the recipient mixture and incubated at 4°C overnight prior to the transposition reaction.

**Protein Transposition on eGFP and Relevant Fluorescence Measurements**

The eGFP recipient protein and $Cat^{C}$-HA-$Cfa^{N}_{m}$ were each diluted to 4 μM in reaction buffer (100 mM phosphate, 150 mM NaCl, 1 mM EDTA, 1 mM TCEP, pH 7.2). The reaction was initiated by mixing 50 μl of eGFP recipient and 50 μl of $Cat^{C}$-HA-$Cfa^{N}_{m}$ together. The fluorescence emission spectrum before and after the transposition reaction (2 hours at room temperature) was measured by using 488 nm as the excitation wavelength and scanning from 490 nm to 550 nm. The change of fluorescence over time during transposition was measured by monitoring the fluorescence intensity at 509 nm while excited at 488 nm; the fluorescence intensity was recorded every 10 seconds for 1 hour.

**Assembly of the Recipient ACF complex**

The ACF complex containing ACF1-FLAG and SMARCA5-$HA_{ins}$ was assembled by equimolar addition of the two separate components into ACF transposition buffer (25 mM HEPES, 60 mM KCl, 10 mM $MgCl_2$, 1 mM TCEP, 10% v/v glycerol, 0.02% v/v IPEGAL CA-630) for 30 minutes at 4 °C. The final concentration of the recipient ACF complex is 1 μM. To confirm the successful formation of ACF complex by immunoprecipitation, 50 μL of the mixture was diluted with ACF transposition buffer to a final volume of 500 μL. 20 μL of anti-FLAG G1 resin (preequilibrated with ACF transposition buffer) was added and the mixture was incubated at 4 °C overnight. The resin was washed with 500 μL of ACF transposition buffer twice and the proteins were eluted with 50 μL ACF transposition buffer containing 0.25 mg/mL of FLAG peptide. The eluted proteins were characterized by SDS-PAGE and western blotting.

**Protein Transposition on the Recipient ACF Complex.**

Prior to protein transposition, the protein transposons were first diluted with ACF transposition buffer to final concentration of 1 μM and incubated at room temperature for 30 minutes. To initiate the transposition, the ACF complex containing SMARCA5-$HA_{ins}$ was mixed with transposons corresponding to SMARCA5(716-746 (i.e. $Cat^{C}$-SMARCA5(716-746)-$Cfa^{N}{}_{m}$) at equal volume and incubated at 4 °C overnight to yield the final ACF complex containing various functional sequences. The products were validated by running them on SDS-PAGE with 7% bis-tris gel at 160 V for 65 min using MOPS-SDS as the running buffer and visualized by Coomassie staining and/or western blotting with indicated antibodies. To confirm the successful formation of ACF complex by immunoprecipitation, 50 μL of the reaction mixture was diluted with ACF transposition buffer to a final volume of 500 μL. 20 μL of anti-FLAG G1 resin (preequilibrated with ACF transposition buffer) was added and the mixture was incubated at 4 °C overnight. The resin was washed with 500 μL of ACF transposition buffer twice and the proteins were eluted with 50 μL ACF transposition buffer containing 0.25 mg/mL of FLAG peptide. The eluted proteins were characterized by SDS-PAGE and western blotting. The complexes were flash-frozen and stored at -80 °C.

The SMARCA5 transposons containing the APB domain used in this study are listed below. APB1 and APB2 controls contain the judicious cysteine mutations to enable protein transposition with minimal effect on SMARCA5 activity. SMARCA5 APB1 transposon was used when the activity of transposed ACF complex was compared with the recipient (Fig. 4C); SMARCA5 APB1 transposon was also used as a positive control when the activity of photoMet-containing ACF complex was validated; the activity of ACF complex containing pY was compared to ACF complex reacted with SMARCA5 APB2.

**SMARCA5 APB1: $Cat^{C}$-CEDYREKQKIAFTEWIEPPKRERKANYAVDAY-$Cfa^{N}{}_{m}$**

**SMARCA5 APB2: $Cat^{C}$-CEDYREKQKIAFTEWIEPPKRERKCNYAVDAY-$Cfa^{N}{}_{m}$**

**SMARCA5 APB+pY:**
**$Cat^{C}$-CEDYREKQKIAFTEWIEPPKRERKCN(phosphoY)AVDAY-$Cfa^{N}{}_{m}$**

**SMARCA5 APB+photoMet:**
**$Cat^{C}$-CEDYREKQKIAFTEWIEPPKRERKCNY(photoMet)VDAY-$Cfa^{N}{}_{m}$**

**Lambda Phosphatase Experiments for an ACF Complex with SMARCA5 (pY742)**

40 μL of ACF complex bearing SMARCA5(pY742) or SMARCA5(APB2) were mixed with 5 μL of 10x PMP buffer (NEB), 5 μL of 10x $MnCl_2$ (NEB) and 0.5 μL of 1 X PMP buffer. For the lambda phosphatase treated samples, 40 μL of ACF complex bearing SMARCA5(pY742) or SMARCA5(APB2) were mixed with 5 μL of 10x PMP buffer (NEB), 5 μL of 10x $MnCl_2$ (NEB) and 0.5 μL of lambda phosphatase (NEB). The mixtures were incubated at 30°C for 1 hour. The treated complexes were flash-frozen and stored at -80°C, and they were used in nucleosome remodeling experiments without further purification.

**Synthesis of SMARCA5 (photoMet743) by Protein Transposition**

Prior to the transposition reaction, the protein transposon containing photo-methionine and recipient SMARCA5 were first diluted with ACF transposition buffer to final concentration of 2 μM and incubated at room temperature for 30 minutes. To initiate the transposition, the recipient SMARCA5 and transposon proteins were mixed at equal volume (final concentration is 1 μM). The mixture was incubated at 4°C overnight to yield the final SMARCA5 containing either the photo-crosslinking group or the control sequence. Successful protein transposition was validated by SDS-PAGE, and the sample was flash-frozen and stored at -80°C.

**Transposition Reactions on Recipient dCas9**

For transposition with HA-tag transposon, purified dCas9 recipient protein was combined with HA-tag transposon in dCas9 storage buffer (50 mM Tris, 250 mM NaCl, 1 mM TCEP, 10% v/v glycerol, pH 7.5) at a final concentration of 1.5 μM and the reaction progress was monitored by SDS-PAGE with Coomassie staining. The SDS-PAGE was run on a 7% bis-tris gel and used MOPS-SDS as the running buffer. The gel band containing the transposition product was excised for further LC-MS/MS validation. For transposition with $PEG_{12}$ transposon, Purified dCas9 recipient protein was combined with $Cat^C$-$PEG_{12}$-$Cfa^N$m in dCas9 storage buffer (50 mM Tris, 250 mM NaCl, 1 mM TCEP, 10% v/v glycerol, pH 7.5) to a final concentration of 1.5 μM (total volume of 1 mL). The reaction mixture was incubated at 4 °C overnight. Unreacted recipient protein, which contains a 6xHis tag, was then removed by incubating the reaction mixture with 100 μL Ni-NTA resin. The flowthrough was collected by centrifugation at 1000 rcf for 5 minutes. The resin was then washed twice with 1 mL of dCas9 storage buffer and the combined flowthroughs concentrated to 250 μL. The purified dCas9 transposition product was aliquoted and stored at -80°C.

**LC-MS/MS Analysis of Transposition Products**

After Coomassie staining of the bis-tris gel used for characterization of transposition products, the bands corresponding to the molecular weight of the product were excised and subjected to in-gel thiol reduction/alkylation and trypsin Gold (Promega) digestion overnight (*13*). Samples were dried completely in a SpeedVac and resuspended with 21 μL of 0.1% formic acid pH 3. 2 μL was injected per run using an Easy-nLC 1200 UPLC system. Samples were loaded directly onto a 45 cm-long 75 μm-inner diameter nano-capillary column packed with 1.9 μm C18-AQ (Dr. Maisch, Germany) mated to metal emitter in-line with an Orbitrap Fusion Lumos (Thermo Scientific, USA). The column temperature was set at 50°C and one-hour gradient method with a flow rate of 300 nL/min flow was used. The mass spectrometer was operated in data dependent mode with the 120,000 resolution MS1 scan (positive mode, profile data type, AGC 4e5, Max IT 54 ms, 300-1500 m/z) in the Orbitrap Fusion Lumos followed by up HCD fragmentation in the ion trap with 35% collision energy. A dynamic exclusion list was invoked to exclude previously fragmented peptides for 60 s and maximum cycle time of 3 s was used. Peptides were isolated for fragmentation using the quadrupole (1.2 Da window). Ion-trap was operated in Rapid mode with AGC $1e^4$, maximum IT of 54 ms and minimum of 5000 ions. Raw files were searched using Sequest HT algorithms (*14*) within the Proteome Discoverer 2.5 suite (Thermo Scientific, USA). 10 ppm MS1 and 0.4 Da MS2 mass tolerances were specified. Carbamidomethylation of cysteine was used as fixed modification, oxidation of methionine, acetylation of protein N-termini as dynamic modifications. Trypsin digestion with maximum of 2 missed cleavages were allowed. Result files were searched against sequences of predicted transposition product and human database downloaded from UniProt.org. Scaffold (version Scaffold_5.3.1, Proteome Software Inc., Portland, OR) was used to validate MS/MS based peptide and protein identifications. Peptide identifications were accepted if they could be established at greater than 95.0% probability by the Scaffold Local FDR algorithm. Protein identifications were accepted if they could be established at greater than 99% probability and contained at least 2 identified peptides. Protein probabilities were assigned by the Protein Prophet algorithm (*15*). Proteins that contained similar peptides and could not be differentiated based on MS/MS analysis alone were grouped to satisfy the principles of parsimony.

#### Methods for dCas9 experiments

##### *In Vitro* Transcription of gRNA

*In-vitro* gRNAs were synthesized using HiScribe T7 High Yield RNA Synthesis Kit (New England Biolabs). The DNA templates for gRNA synthesis were synthesized by Twist Bioscience followed by standard PCR amplification conditions.

The DNA template sequence is listed below:

IL1RN (T7 Consensus Promoter-Complementary gRNA-TracrRNA):

GAAATTAATACGACTCACTATAGGGacgcagataagaaccagttgtttaagagctatgctggaaacagcatagcaa gtttaaataaggctagtccgttatcaacttgaaaaagtggcaccgagtcggtgc

Gal4-1 (T7 Consensus Promoter-Non-complementary gRNA-TracrRNA):

GAAATTAATACGACTCACTATAGGGGGAGTACTGTCCTCCGAGgtttaagagctatgctggaaaca gcatagcaagtttaaataaggctagtccgttatcaacttgaaaaagtggcaccgagtcggtgc

Following gRNA amplification, the reaction mixture was treated with RNase-free DNase I (Qiagen 79254; 2ul/100uL of reaction mixture at 25 °C for 15 mins) and extracted with phenol:chloroform. Finally, ethanol precipitation was performed, and the gRNAs were resuspended in RNase-free water. gRNAs were denatured at 68 °C for 10 min and immediately put on ice to remove the secondary structure. gRNAs were flash frozen and stored in single-use aliquots at -80 °C. See Fig. S33 for details.

##### *In Vitro* dCas9:gRNA:DNA Binding Gel Shift Assay

The IL1RN promoter sequence were amplified from a reporter plasmid (*16*) by standard PCR conditions. Products were purified by PCR purification and gel extract columns (Thomas Scientific 1158P49).

Sequence (fragment from IL1RN gene promoter, Blue: target sequence) as follows:

ggtaagcacgaaggcccagctcagttctctgcatgtgacctcccatcttacgcagataagaaccagtttggtttctgctagcctgagtcaccct cctggaaactgggcctgcttggcatcaagtcagccatcagccggcccatctcctcatgctggccaaccctctgtgagtgtgtgggaggggagactgggctcctccttgtactctctgaggtgctctggaaggagaagcttggcaatccggtactgttgg

Primer (5' to 3'):

IL1RN_DNA_F: ggtaagcacgaaggcccagctcag

IL1RN_DNA_R: ccaacagtaccggattgccaagcttctcc

The dCas9 proteins (dCas9 WT and *in vitro* transposition products), gRNA and DNA were mixed together to reach a final concentration of 100 nM dCas9: 500 nM gRNA: 10 nM DNA. The mixtures were then incubated at 37 °C for 2 hours. For gel shift assays, 50% of sucrose was added to the samples to reach a final concentration of 10% (v/v). Samples were then loaded onto a 5% native PAGE TBE gel and run with 0.5x TBE Buffer. Finally, the native TBE gels were stained with SYBR™ Gold Nucleic Acid Gel Stain (Invitrogen™ S11494) at 1:10,000 in 0.5x TBE Buffer for imaging.

#### Methods for Nucleosome Remodeling by Designer ACF Complexes

##### Nucleosome preparation

Nucleosomes used in crosslinking and remodeling assays were prepared following standard procedures (*6, 17*). Briefly, recombinant human histones (H2A, H2B, H3.1 and H4) were expressed in *E. coli* and purified under denaturing conditions. The purified histones were then mixed at approximately equimolar ratios and the mixture was dialyzed against 2 M NaCl to generate histone octamers. Octamers were purified by size-exclusion chromatography. 207 bp DNA consisting of the Widom 601 sequence (modified to contain a PstI site) with 45 and 15 bp overhangs was prepared by polymerase chain reaction using a 6FAM (6-carboxyfluorescein)-labeled (IDT) primer at one end.

The sequence of the DNA is:

**6FAM-**

*GTGGCGGCCGCTCTAGAACTAGTGGATCCGATATCGCTGCCATGG*ACAGGATGTATATA

TCTGACACGTGCCTGGAGACTAGGGAGTAATCCCCTTGGCGGTTAAAACGCGGGGG ACAGCGCGTACGTGCGTTTAAGCGGTGCTAGAGCTGTCTACGACCAATTGAGCGGCTGCAGCACCGGGATTCTCCAG*CATCAGAGACCTAGG*

where the underlined portion is the Widom 601 sequence and overhangs are italicized. The PstI site is denoted in red. Nucleosomes were assembled by mixing this DNA with histone octamers and gradually reducing the ionic strength by salt gradient dialysis. The final buffer was 10 mM KCl, 10 mM Tris, 0.1 mM EDTA, 1 mM DTT. The molar ratio of DNA to octamer was chosen empirically to optimize the purity of the final nucleosomes, as assessed by native gel electrophoresis (7% acrylamide, 190 V for 35 min). Purity of the assembled nucleosomes was verified by fluorescence in the FAM channel or by staining with SYBR™ Gold (2 uL in 30 mL of 0.5x TBE) for ~10 mins and imaging in the Cy3 channel:

**Remodeling assays**

Remodeling by ACF was observed using the restriction enzyme accessibility assay (REAA) and electrophoretic mobility shift assay (EMSA) for nucleosome sliding, as previously described, with some modifications (*6, 11, 18*).

For REAA experiments, 10 nM mononucleosomes and 10 nM ACF were combined in REA buffer (12 mM HEPES, pH 7.6, 4 mM Tris, 60 mM KCl, 10 mM $MgCl_2$, 10% glycerol, 0.02% (v/v) IGEPAL CA-630) in the presence of 2 U/uL PstI-HF restriction enzyme. To allow for cutting of any free DNA, all reaction components except ATP were incubated for 20 min at 30 °C. ATP was then added to a final concentration of 2 mM to initiate remodeling. Reactions were carried out for a total of 64 min at 30 °C, with 6 uL aliquots removed at 0, 2, 4, 8, 16, 32, and 64 min and immediately quenched with 9 uL of quench buffer (20 mM Tris, 70 mM EDTA, pH 8.0, 10% (v/v) glycerol, 2% SDS, 20 U/mL Proteinase K). One additional sample was prepared without ATP and was incubated for 64 min at 30 C to serve as a negative control. Following quenching, all samples were incubated for 1 h at 37 °C to allow for protein digestion. Samples (DNA remaining) were analyzed on a 7% TBE gel stained with SYBR Gold Nucleic Acid Gel Stain and remodeling rates calculated as described previously.

For EMSA experiments, 10 nM mononucleosome substrate and 8.33 nM ACF were combined in REA buffer (12 mM HEPES, pH 7.6, 4 mM Tris, 60 mM KCl, 10 mM $MgCl_2$, 10% glycerol, 0.02% (v/v) IGEPAL CA-630). ATP was added to a final concentration of 2 mM to initiate remodeling, and the reactions (total volume 6 uL) were incubated at 30 °C. After 4 minutes, reactions were quenched using 1 uL of 6x purple loading dye, no SDS (from NEB; contains 60 mM EDTA, 20 mM Tris-HCl, 15% Ficoll-400) supplemented with sheared salmon sperm DNA (1 mg/mL) to titrate ACF from the nucleosomes. Samples were analyzed by native gel electrophoresis (7% acrylamide) and imaged in the FAM channel.

**Photo-crosslinking Protocol**

Nucleosomes (5 pmol, 1 equiv.) and semisynthetic or recombinant SMARCA5 (10 pmol, 2 equiv.) were combined in a 30-µl reaction of REAA buffer (20 mM HEPES, 4 mM Tris-HCl pH 7.75, 60 mM KCl, 10 mM $MgCl_2$, 0.02% (v/v) IGEPAL CA-630/NP-40, 10% glycerol). For samples containing LANA peptide (10 µM), the peptide was preincubated with the nucleosomes at 4 °C for 30 min before addition of SMARCA5. The reaction mixture was incubated at 4 °C 10 min, followed by an increase in temperature to 30 °C for 30 min, then switched to 4 °C for 20 min. The photocrosslinking reaction was then carried out using UV irradiation at 4 °C for 20 min with 365 nm UV light (MAXIMA ML-3500S). The samples were then immediately flash frozen and lyophilized for 1 h away from light. The lyophilized samples were then dissolved in 1X loading buffer (10 uL per sample) and the protein mixtures were boiled at 95 °C for 10 min and resolved by SDS–PAGE (4%–12% Bis-Tris, 1 h 25 min at 150 V). The crosslinked products were transferred to a nitrocellulose membrane (0.7 A, 2 h), which was then blocked with 5% w/v nonfat dry milk in TBST (20 mM Tris-Cl, pH 7.6, 150 mM NaCl, 0.1% v/v Tween-20) for 30 mins at room temperature. Next, membranes were incubated with the indicated primary antibodies (diluted in TBST containing 1% BSA) at 4 °C overnight. After washing three times with TBST, the appropriate Li-Cor IRDye secondary antibodies (1:10000 dilution in TBST containing 1% BSA) were applied for 1 h at room temperature. Following incubation with the dye-labeled secondary antibodies, membranes were washed 3 times with TBST and imaged with Li-Cor Odyssey Infrared Imaging System.

***In Nucleo* Protein Trans-Splicing Protocol**

One plate of HEK 293T cells reaching 60% confluency was transfected with 5ug of pCDNA-SMARCA5$^{N}$(1-715)-CatN-HA-CfaC-SMARCA5$^{C}$(747-1051) plasmid using Invitrogen™ Lipofectamine™ 3000 (L3000001) in line with the manufacturer's instruction. After 24 hours, the cells were lysed under hypotonic conditions using 1mL of RSB buffer (10 mM Tris pH 7.4, 15 mM NaCl, 1.5 mM $MgCl_2$, Roche cOmplete EDTA-free protease inhibitors) for 10 min on ice. The crude nuclei were isolated by centrifugation (Note: all centrifugations were conducted at 500 g for 5 min at 4 °C unless noted otherwise), resuspended in 1 ml RSB buffer, and homogenized for using ten strokes of a loose pestle Dounce homogenizer. The isolated nuclei were then pelleted by centrifugation and resuspended in 500 μL of transposition buffer (20 mM HEPES, 1.5 mM $MgCl_2$, 150 mM KCl, Roche cOmplete EDTA-free protease inhibitors, pH 7.4). The nuclei were centrifuged and washed again with 500 μL of transposition buffer. Finally, the pelleted nuclei were resuspended in 300 μL of transposition buffer per 10 cm dish of transfected HEK 293T cells. A indicated amount of Cat$^{C}$-FLAG-Cfa$^{N}_{m}$ in transposition buffer was added to each nuclei aliquot. The reactions were incubated at 37 °C for 30 min. The samples were then washed twice with 200 μL of transposition buffer. Next, the treated nuclei were dissolved in 1xSDS sample loading buffer (100 mM Tris-Cl, pH 6.8, 3% SDS, 15% Glycerol, 2.25% β-mercaptoethanol, 0.015% bromophenol blue). Samples were boiled at 98 °C for 30 mins and were subsequently loaded on a 7% bis-tris polyacrylamide gel. Proteins were transferred to a nitrocellulose membrane which was then blocked with 5% w/v nonfat dry milk in TBST (20 mM Tris-Cl, pH 7.6, 150 mM NaCl, 0.1% v/v Tween-20) for 30 mins at room temperature. Next, membranes were incubated with the indicated primary antibodies (diluted in TBST containing 1% BSA) at 4 °C overnight. After washing three times with TBST, the appropriate Li-Cor IRDye secondary antibodies (1:10000 dilution in TBST) were applied for 1 h at room temperature. Following incubation with the dye-labeled secondary antibodies, membranes were washed 3 times with TBST and imaged with Li-Cor Odyssey Infrared Imaging System.

Antibodies and Concentrations Used

| **Epitope** | **Antibody** | **Vendor** | **Western Blot Dilution** |
|---|---|---|---|
| H2A | Rabbit anti-H2A | Abcam (ab18255) | 1:2000 |
| FLAG | Mouse anti-FLAG | Sigma Aldrich (F1804) | 1:3000 |
| FLAG | Rabbit anti-FLAG | Sigma Aldrich (F7425) | 1:1000 |
| HA | Rabbit anti-HA | Abcam ab9110 | 1:3000 |
| HA | Mouse anti-HA | Proteintech 66006-2-1g | 1:3000 |
| Phospho-tyrosine | Rabbit anti-pY | EMD Millipore 4G10 | 1:2000 |
| SMARCA5 | Rabbit anti-SMARCA5 | Abcam (ab72499) | 1:5000 |
| Rabbit IgG | Goat IRDye 680RD anti-Rabbit IgG (H + L) | LI-COR (926-68071) | 1:10,000 |
| Rabbit IgG | Goat IgG IRDye 800CW anti-Rabbit IgG (H + L) | LI-COR (926-32211) | 1:10,000 |
| Mouse IgG | Goat IgG IRDye 680RD anti-Mouse IgG (H + L) | LI-COR (926-68070) | 1:10,000 |
| Mouse IgG | Goat IgG IRDye 800CW anti-Mouse IgG | LI-COR (926-32210) | 1:10,000 |

#### Supplemental Figures

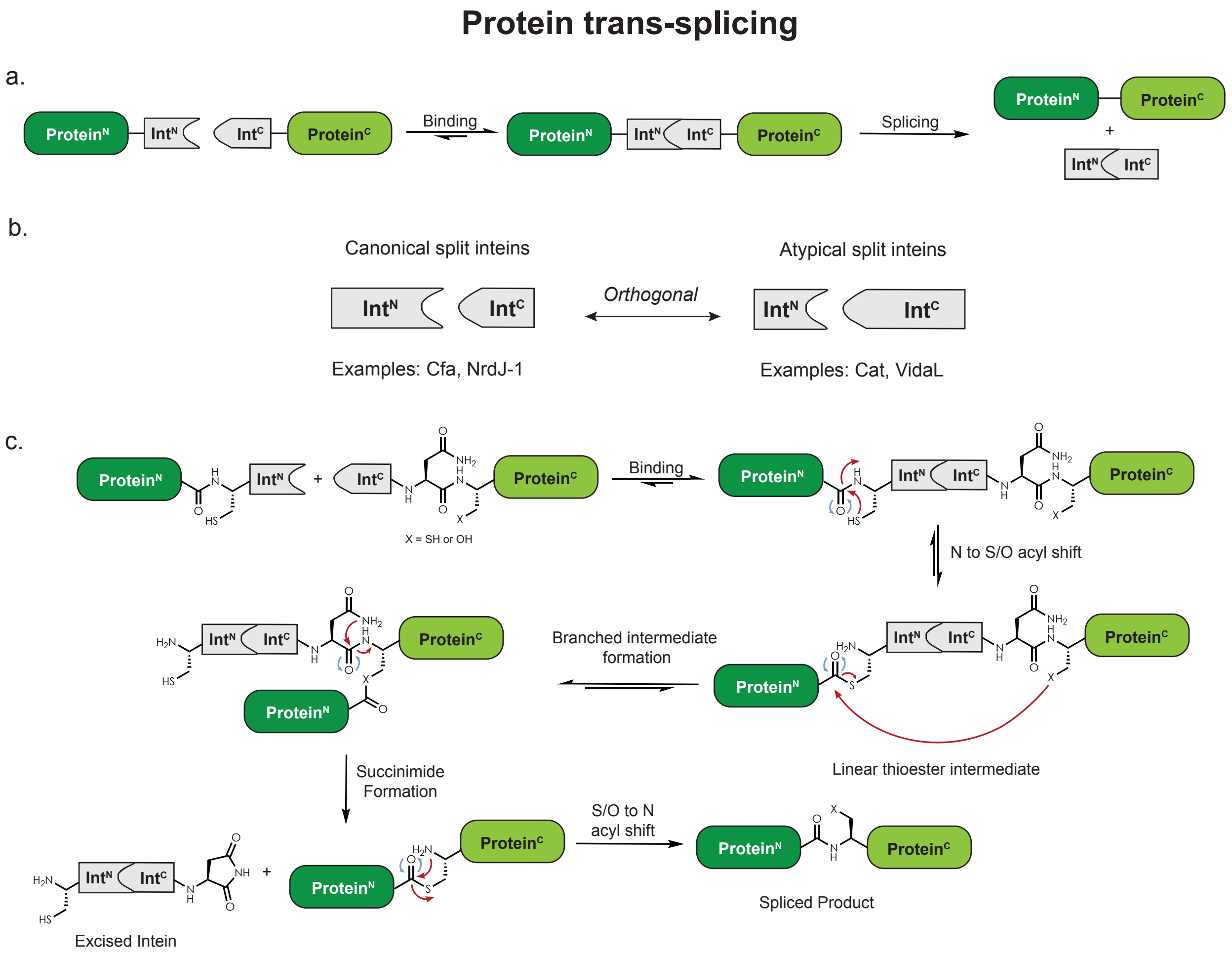


**Fig. S1.** Schematic of Protein Trans-Splicing with Split-Inteins. (a). General mechanism for split-intein trans-splicing. (b). Split inteins are characterized as either canonical or atypical based on the lengths of their N- and C-intein fragments. (c) Detailed mechanism of protein trans-splicing.

**HA-MBP-Vid$^N$-TEV-Cfa$^C$-EGFP-FLAG**
(HA-MBP-NESG-Vid$^N$-GSESNKRKSNFSENLYFQGSKKKREQSNDIARG-Cfa$^C$-CFN-EGFP-FLAG)

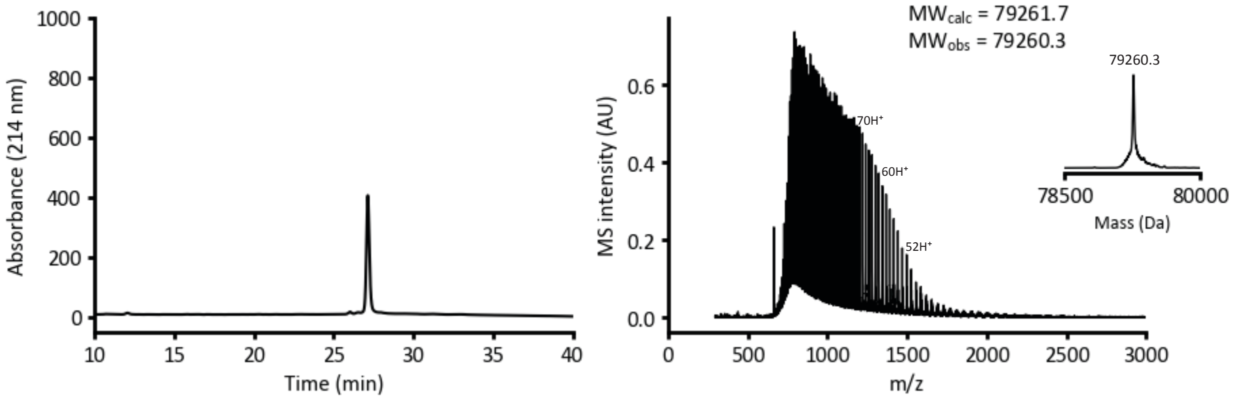


**HA-MBP-Cat$^N$-TEV-Cfa$^C$-EGFP-FLAG**
(HA-MBP-NESG-Cat$^N$-GSESNKRKSNFSENLYFQGSKKKREQSNDIARG-Cfa$^C$-CFN-EGFP-FLAG)

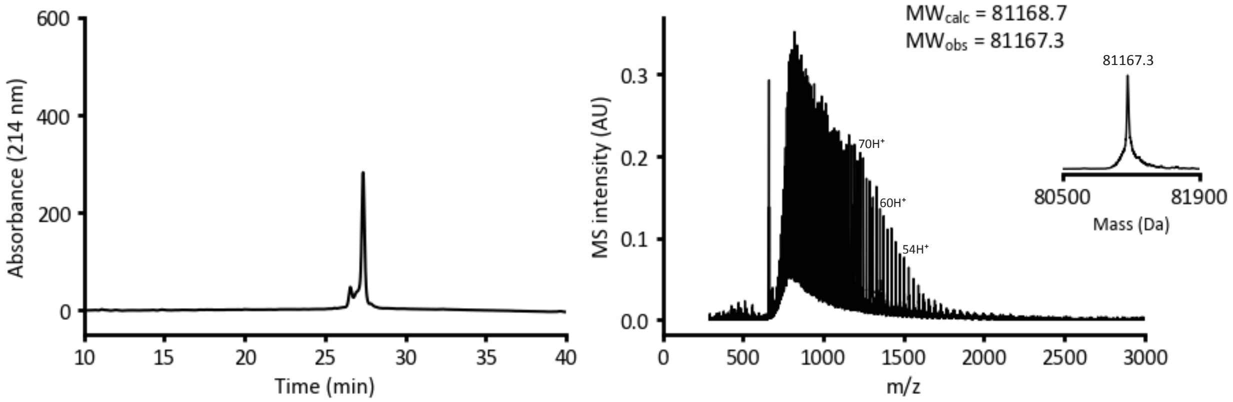


**Fig. S2. Characterization of Model MBP-eGFP Fusion Proteins for Transposition by RP-HPLC and ESI-MS.** Top: Analytical C-18 RP-HPLC chromatogram (gradient used: 20-75% B, 10-40 min) (left) for the model protein containing VidaL$^N$ and Cfa$^C$ split inteins embedded between MBP and eGFP, and the corresponding ESI-MS spectra with deconvoluted spectra shown in insert (right). Bottom: Analytical C-18 RP-HPLC chromatogram (gradient used: 20-75% B, 10-40 min) (left) for the model protein containing Cat$^N$ and Cfa$^C$ split inteins, and the corresponding ESI-MS spectra with deconvoluted spectra shown in insert (right).

**Vid$^{C}$-HA-Cfa$^{N}$** (Vid$^{C}$-CEASGKYPYDVPDYAGAEY-Cfa$^{N}$)

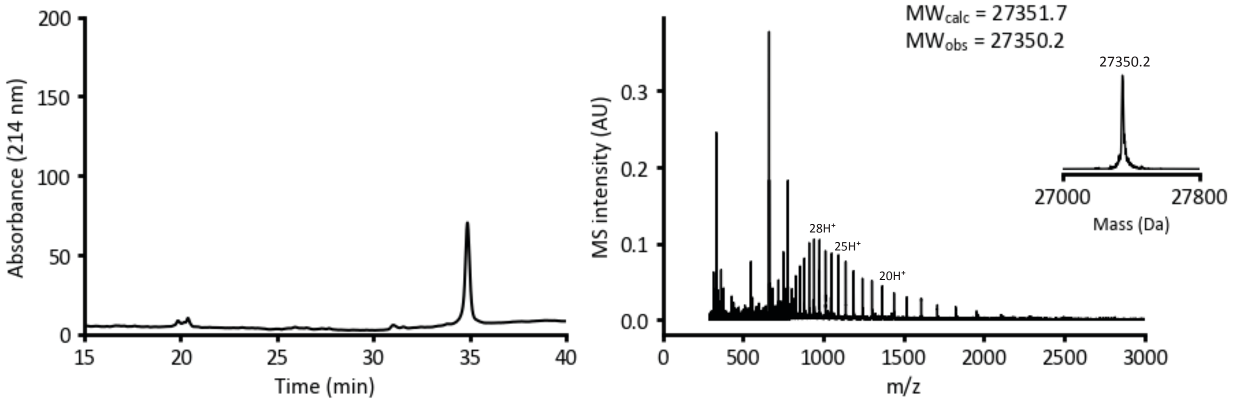


**Cat$^{C}$-HA-Cfa$^{N}$** (Cat$^{C}$-CEASGKYPYDVPDYAGAEY-Cfa$^{N}$)

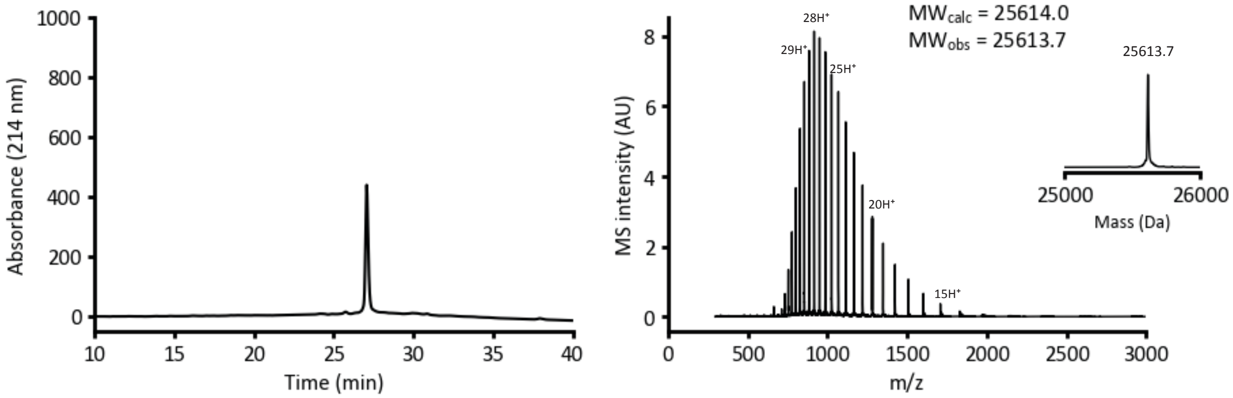


**Fig. S3. Characterization of HA-Containing Protein Transposons for Transposition by RP-HPLC and ESI-MS.** Top: Analytical C-4 RP-HPLC chromatogram (gradient used: 20-75% B, 15-40 min) (left) for the transposon containing VidaL$^{C}$ and Cfa$^{N}$ split inteins flanking the HA tag (in red), and the corresponding ESI-MS spectra with deconvoluted spectra shown in insert (right). Bottom: Analytical C-18 RP-HPLC chromatogram (gradient used: 20-75% B, 10-40 min) (left) for the transposon containing Cat$^{C}$ and Cfa$^{N}$ split inteins flanking the HA tag (in red), and the corresponding ESI-MS spectra with deconvoluted spectra shown in insert (right).

a.

Protein Receeipient: HA-MBP-NESG-**Vid$^{N}$**-SESNKRKSNFSENLYFQGSKKKREQSNDIARG-**Cfa$^{C}$**-CFN-eGFP-FLAG
Protein Transposon: **Vid$^{C}$**-SGKYPYDVPDYAGAEY-**Cfa$^{N}$/Cfa$^{N}_{m}$**
Protein Product: HA-MBP-NESGSGKYPYDVPDYAGAEYCFN-eGFP-FLAG

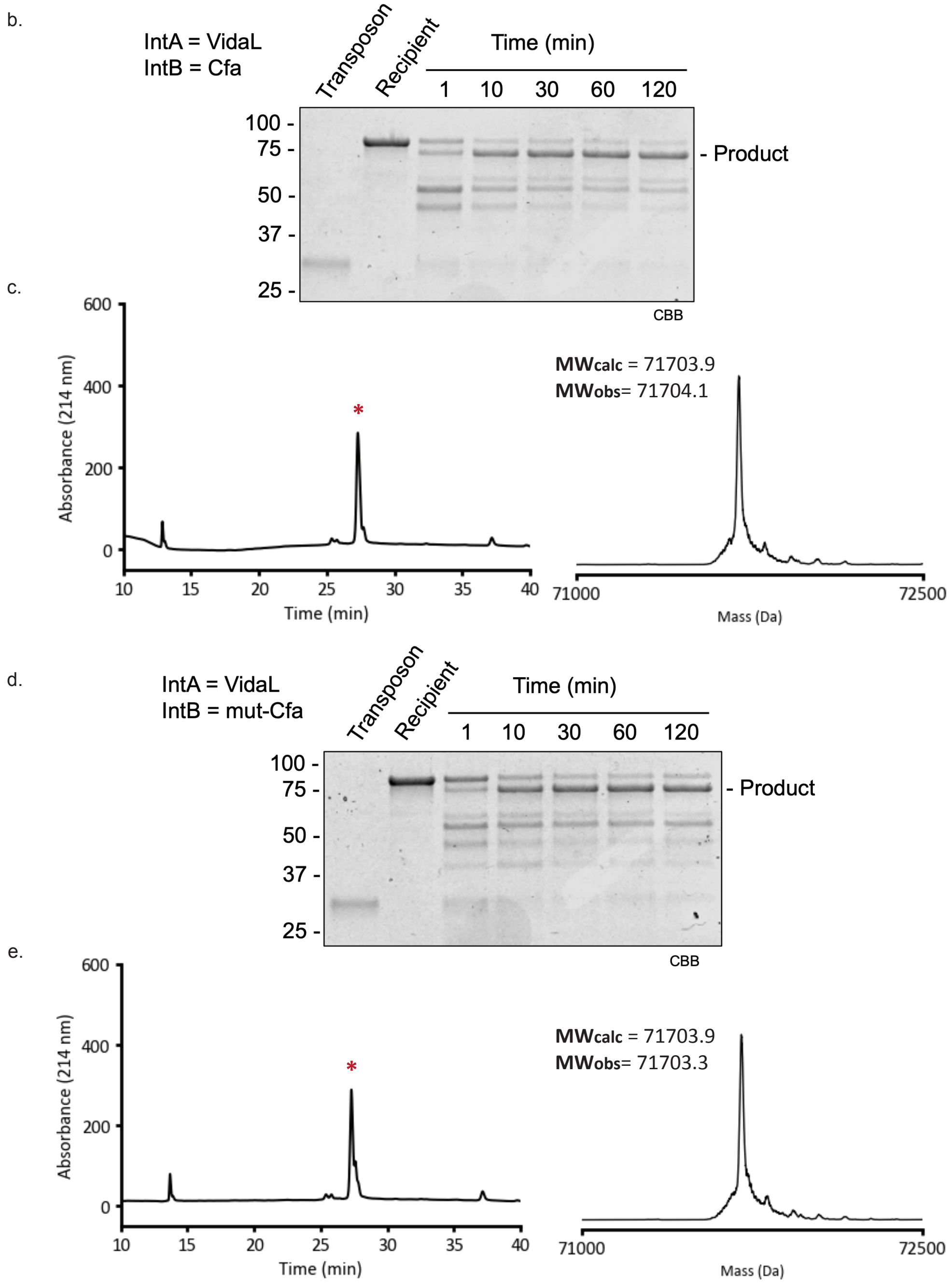


**Fig. S4.** Protein transposition using (a) orthogonal split intein pairs VidaL and Cfa/mut-Cfa fused to a MBP-eGFP protein recipient and a HA tag protein transposon. The transposition reaction was initiated by combining recipient and transposon (2 µM) in 100 mM phosphate, 150 mM NaCl, 1 mM EDTA, 1 mM TECP, pH 7.2). The mixture was incubated at room temperature for 2 hours, with reaction progress monitored at different time points and characterized by SDS-PAGE with Coomassie brilliant blue (CBB) staining (b, d). The final reaction products (red asterisk in the

HPLC trace denotes the transposition product) were separated by C-18 RP-HPLC (gradient used: 20-75% B, 10-40 min) and the deconvoluted ESI-MS spectrum of the product is shown (c, e).

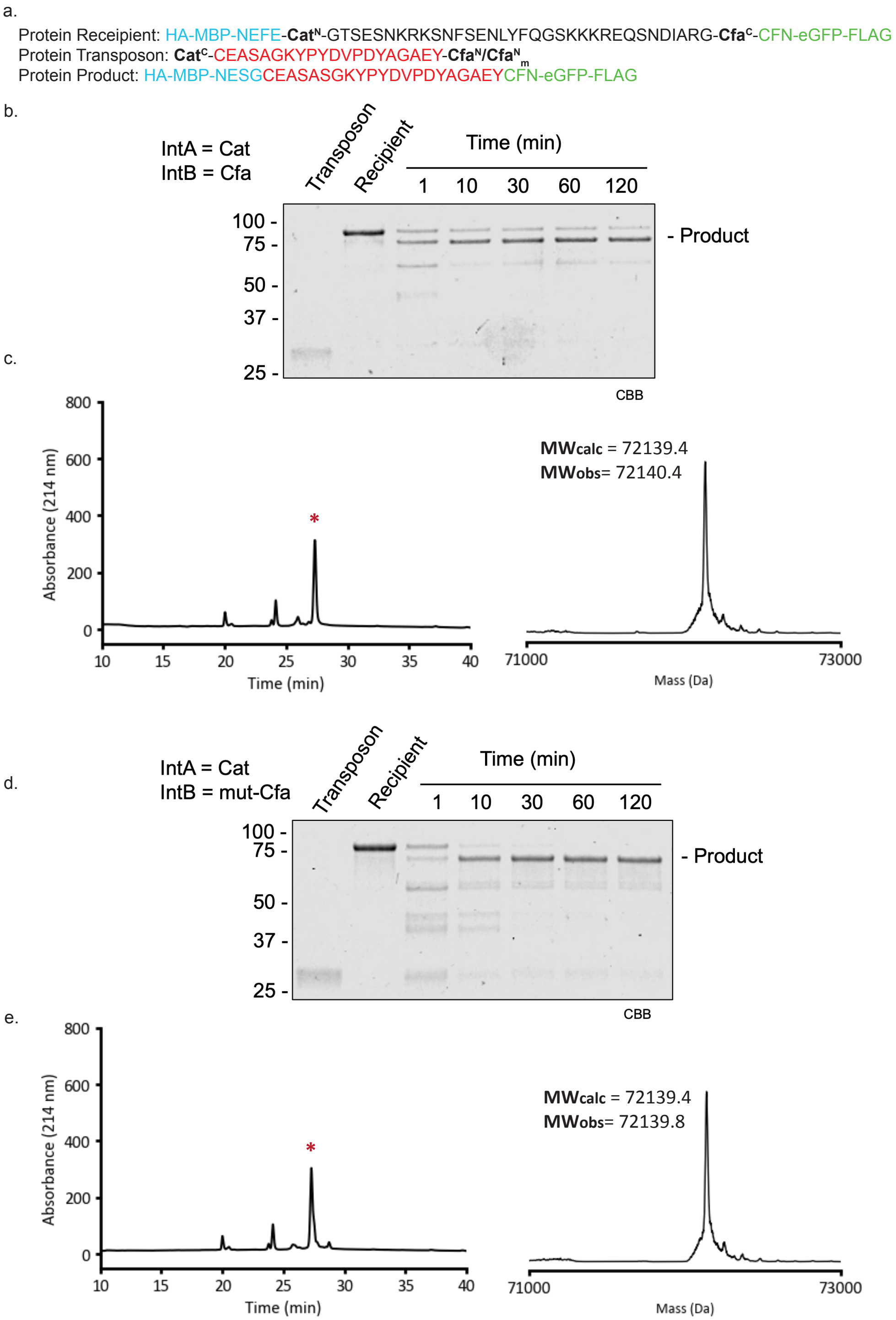


**Fig. S5.** Protein transposition using (a) orthogonal split intein pairs Cat and Cfa/mut-Cfa fused to a MBP-eGFP protein recipient and a HA tag protein transposon. The transposition reaction was initiated by combining recipient and transposon (2 μM) in 100 mM phosphate, 150 mM NaCl, 1 mM EDTA, 1 mM TECP, pH 7.2). The mixture was incubated at room temperature for 2 hours, with reaction progress monitored at different time points and characterized by SDS-PAGE with Coomassie brilliant blue (CBB) staining (b, d). The final reaction products (red asterisk in the HPLC trace denotes the transposition product) were separated by C-18 RP-HPLC (gradient used: 20-75% B, 10-40 min) and the deconvoluted ESI-MS spectrum of the product is shown (c, e).

##### HA-MBP-Vid$^{N}$-TEV-Cfa$^{C}$-EGFP-FLAG ($^{15}$N)

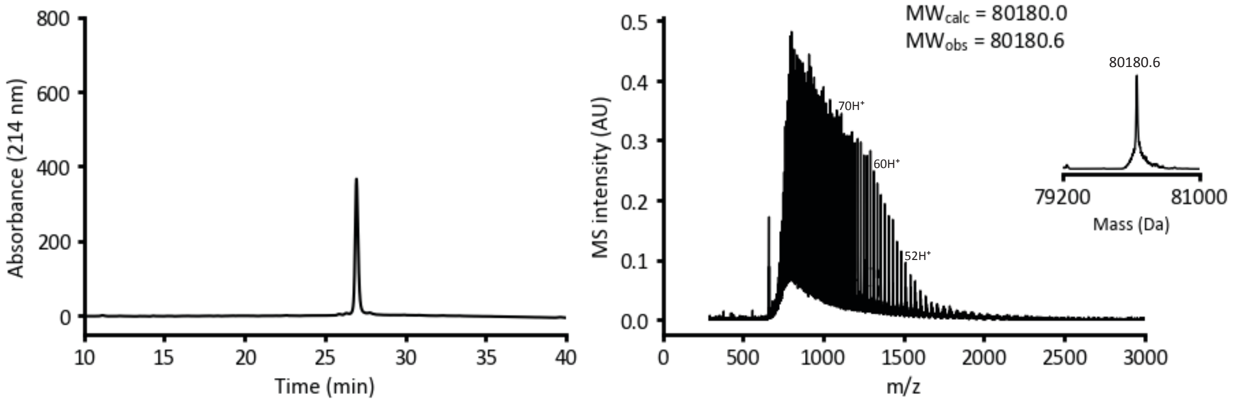


##### HA-MBP-Cat$^{N}$-TEV-Cfa$^{C}$-EGFP-FLAG ($^{15}$N)

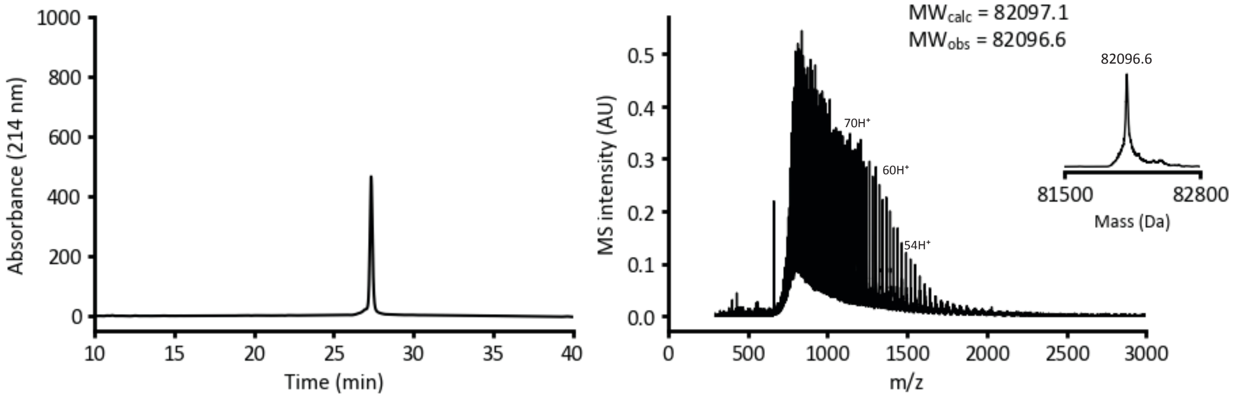


**Fig. S6. Characterization of $^{15}$N Isotopically-Labeled Model MBP-eGFP Fusion Proteins for Transposition by RP-HPLC and ESI-MS.** Top: Analytical C-18 RP-HPLC chromatogram (gradient used: 20-75% B, 10-40 min) (left) for the $^{15}$N isotopically-labeled model protein containing VidaL$^{N}$ and Cfa$^{C}$ split inteins embedded between MBP and eGFP, and the corresponding ESI-MS spectra with deconvoluted spectra shown in insert (right). Bottom: Analytical C-18 RP-HPLC chromatogram (gradient used: 20-75% B, 10-40 min) (left) for the $^{15}$N isotopically-labeled model protein containing Cat$^{N}$ and Cfa$^{C}$ split inteins, and the corresponding ESI-MS spectra with deconvoluted spectra shown in insert (right).

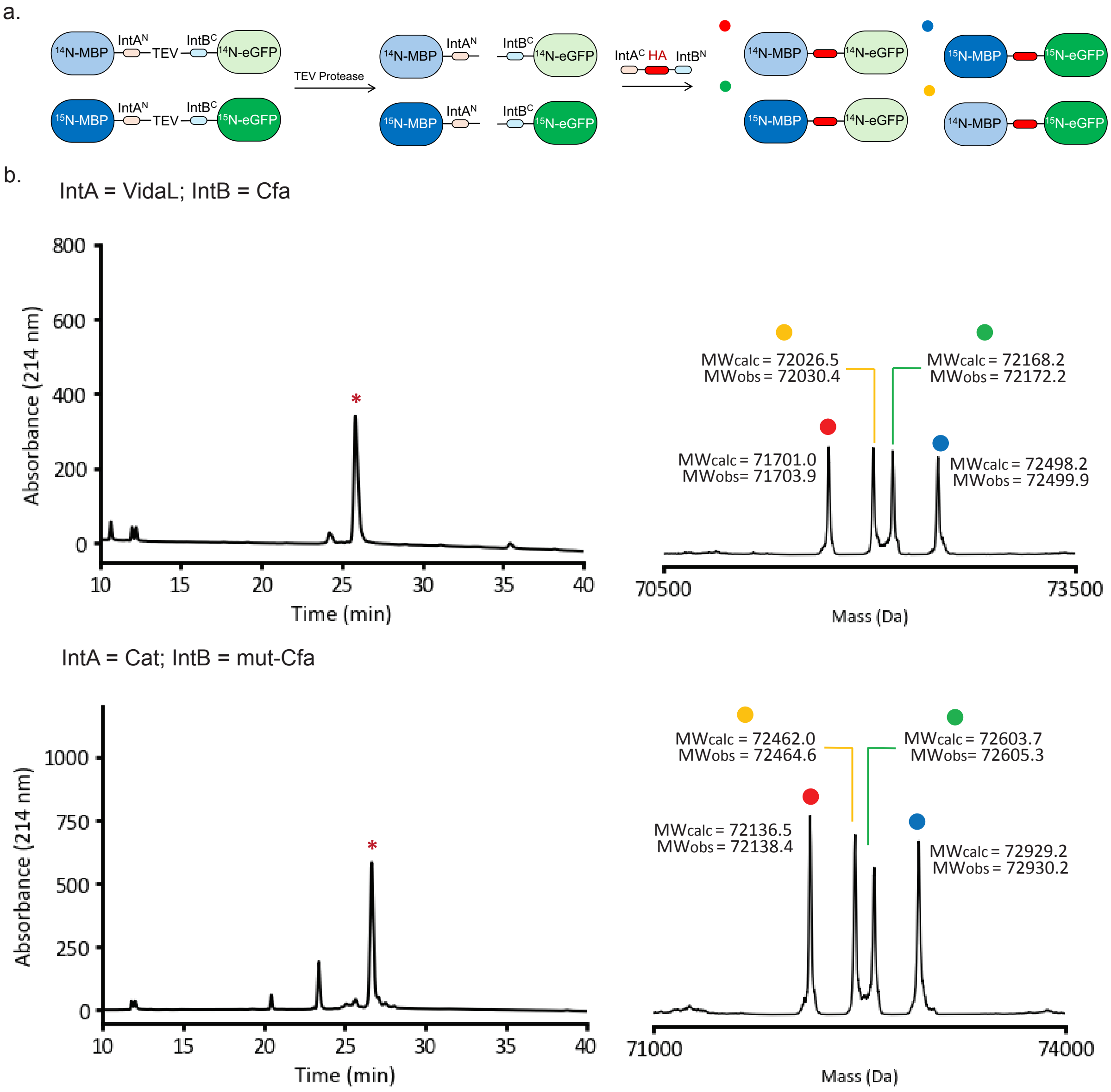


**Fig. S7. TEV Protease-Mediated Cleavage of MBP-TEV-eGFP Proteins Serves as a Positive Control for Stepwise Protein Transposition.** When light ($^{14}$N) and heavy ($^{15}$N) MBP-TEV-eGFP protein recipients were mixed (both at 1 µM) and treated with TEV protease (500 nM) at 4°C overnight, the MBP and eGFP components of the fusion are no longer associated, which leads to stepwise protein transposition when treated with a HA tag protein transposon (2 µM). The transposition reaction was carried out in 100 mM phosphate, 150 mM NaCl, 1 mM EDTA, 1 mM TCEP, pH 7.2 at room temperature for 2 hours. (b) This mechanistic outcome is observed by equal isotopic distribution of the four transposition outcomes (denoted by each respective colored ball) regardless of intein pair. The protein mixture (red asterisk in the HPLC trace denotes the transposition product) was purified by analytical C-18 RP-HPLC (gradient used: 20-75% B, 10-40 min) and the identity of each isotopic product was determined by the deconvoluted ESI-MS spectra. Each different isotopically-labeled product is matched to its corresponding colored ball.

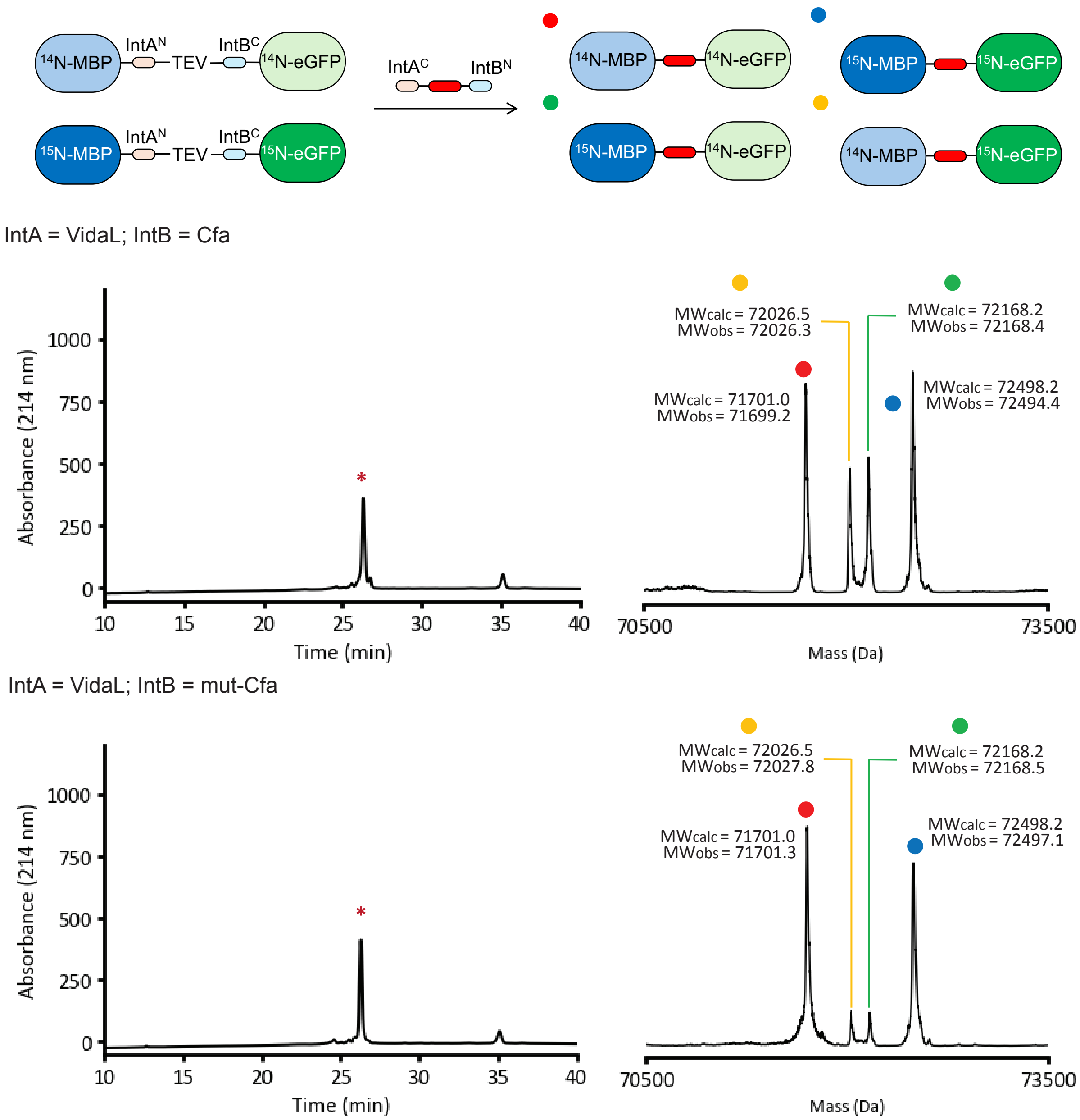


**Fig. S8. Mixed-Isotope Transposition Experiment Comparing VidaL and Cfa/mut-Cfa kinetics.** The product distribution from the mixed-isotope transposition experiment using orthogonal split intein pairs VidaL and Cfa/mut-Cfa inserted into a MBP-eGFP protein recipient (1 µM of each isotopically labeled recipient) and a HA tag protein transposon (2 µM). The reaction was carried out in 100 mM phosphate, 150 mM NaCl, 1 mM EDTA, 1 mM TCEP, pH 7.2 at room temperature for 2 hours. The protein mixture (red asterisk in the HPLC trace denotes the transposition product ) was purified by analytical C-18 RP-HPLC (gradient used: 20-75% B, 10-40 min) and the identity of each isotopic product was determined by the deconvoluted ESI-MS spectra. The mechanistic outcome of the transposition, as identified by the ESI-MS results, is then correlated to the respective colored ball denoting its identity. When using the VidaL/Cfa pair, the stepwise pathway is dominant as an increase in the scrambled products ($^{14}$N-MBP-$^{15}$N-eGFP and

$^{15}$N-MBP-$^{14}$N-eGFP) was observed. The isotope scrambled products are significantly reduced when Cfa is replaced by mut-Cfa, which was predicted to have a slower splicing rate, which is hypothesized to match the binding and kinetics of the VidaL better than wild-type Cfa.

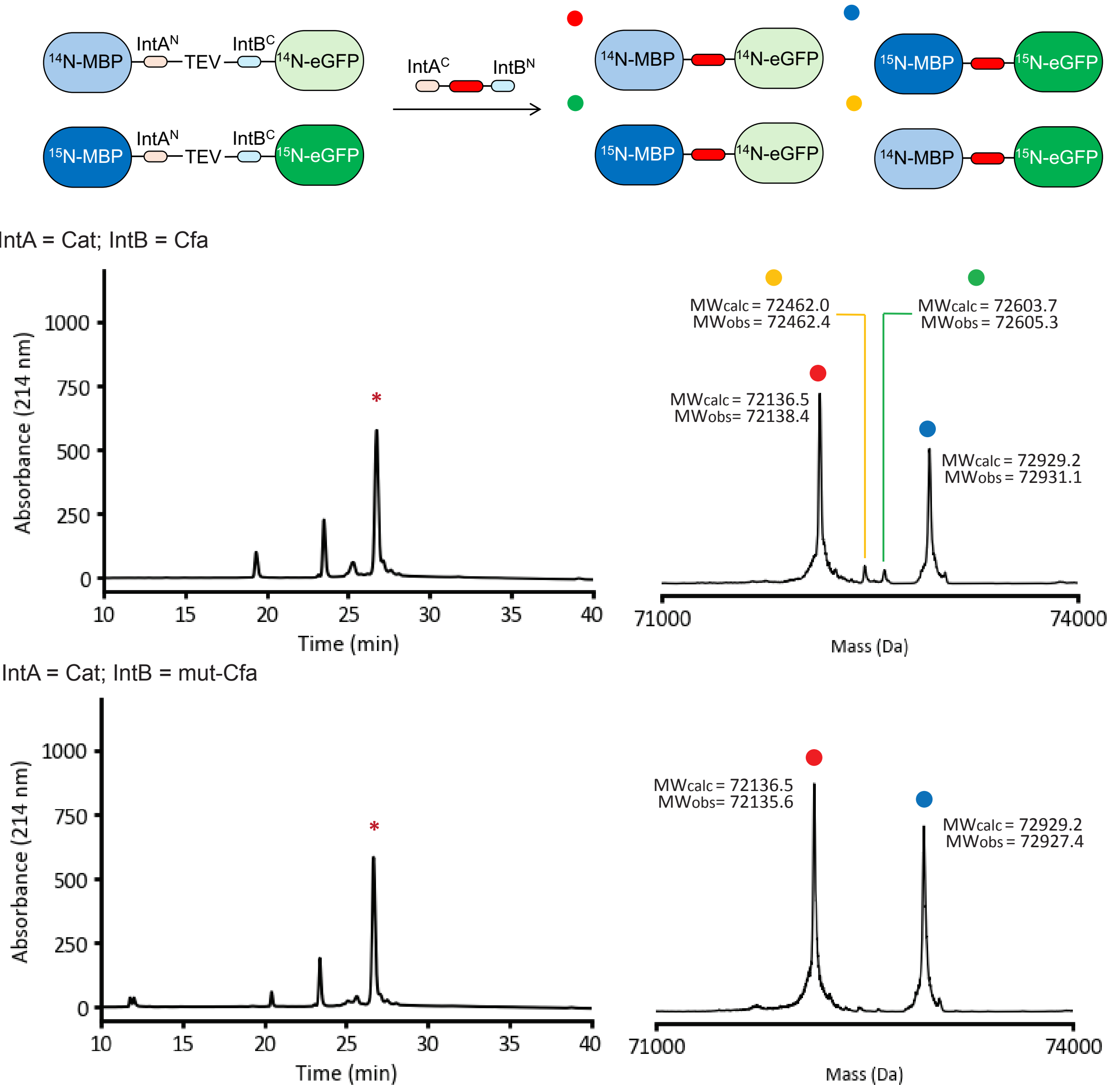


**Fig. S9. Mixed-Isotope Transposition Experiment Comparing Cat and Cfa/mut-Cfa kinetics.** The product distribution from the mixed-isotope transposition experiment using orthogonal split intein pairs Cat and Cfa/mut-Cfa inserted into a MBP-eGFP protein recipient (1 µM of each isotopically labeled recipient) and a HA tag protein transposon (2 µM). The reaction was carried out in 100 mM phosphate, 150 mM NaCl, 1 mM EDTA, 1 mM TCEP, pH 7.2 at room temperature for 2 hours. The protein mixture (red asterisk in the HPLC trace denotes the transposition product) was purified by analytical C-18 RP-HPLC (gradient used: 20-75% B, 10-40 min) and the identity of each isotopic product was determined by the deconvoluted ESI-MS spectra. The mechanistic outcome of the transposition, as identified by the ESI-MS results, is then correlated to the respective colored ball denoting its identity. Using the Cat/Cfa pair, minimal amounts of scrambled products were observed, which can be further ameliorated by using the Cat/mut-Cfa pair. This observation suggests that protein transposition using the Cat/mut-Cfa pair proceeds in a nearly fully concerted manner.

**Vid$^{C}$-HA-Cfa$^{N}_{m}$** (Vid$^{C}$-CEASGKYPYDVPDYAGAEY-Cfa$^{N}_{m}$)

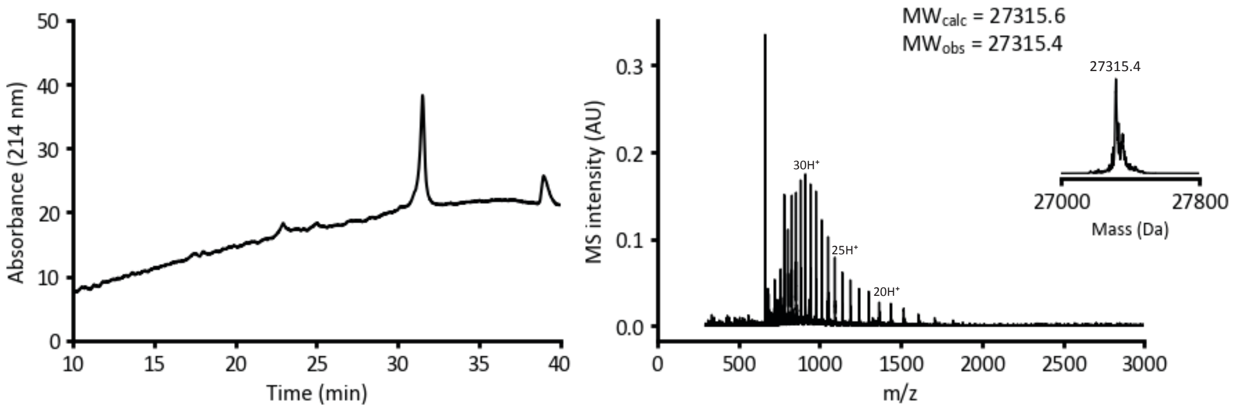


**Cat$^{C}$-HA-Cfa$^{N}_{m}$** (Cat$^{C}$-CEASGKYPYDVPDYAGAEY-Cfa$^{N}_{m}$)

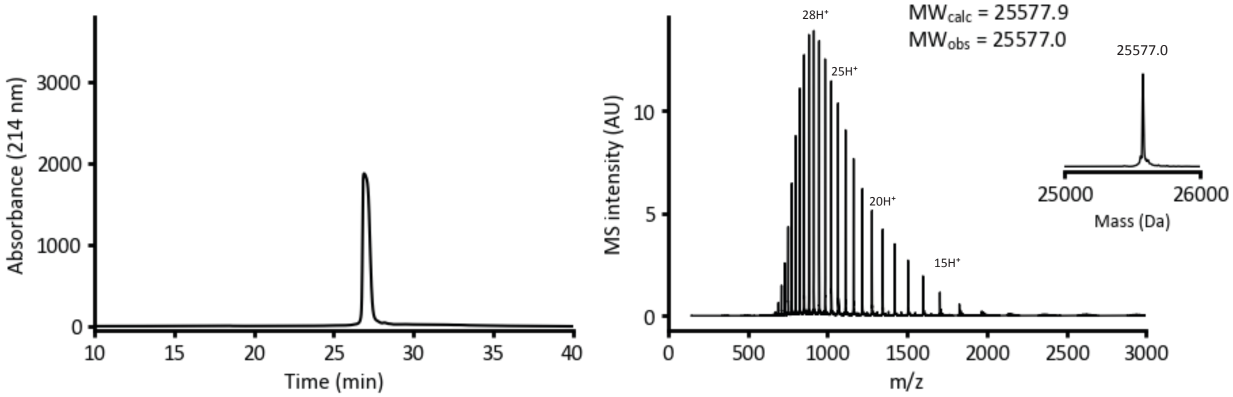


**Fig. S10. Characterization of mut-Cfa$^{N}$ and HA-Containing Protein Transposons for Transposition by RP-HPLC and ESI-MS.** Top: Analytical C-4 RP-HPLC chromatogram (gradient used: 20-75% B, 15-40 min) (left) for the transposon containing VidaL$^{C}$ and Cfa$^{N}_{m}$ split inteins flanking the HA tag (in red), and the corresponding ESI-MS spectra with deconvoluted spectra shown in insert (right). Bottom: Analytical C-18 RP-HPLC chromatogram (gradient used: 20-75% B, 10-40 min) (left) for the transposon containing Cat$^{C}$ and Cfa$^{N}_{m}$ split inteins flanking the HA tag (in red), and the corresponding ESI-MS spectra with deconvoluted spectra shown in insert (right).

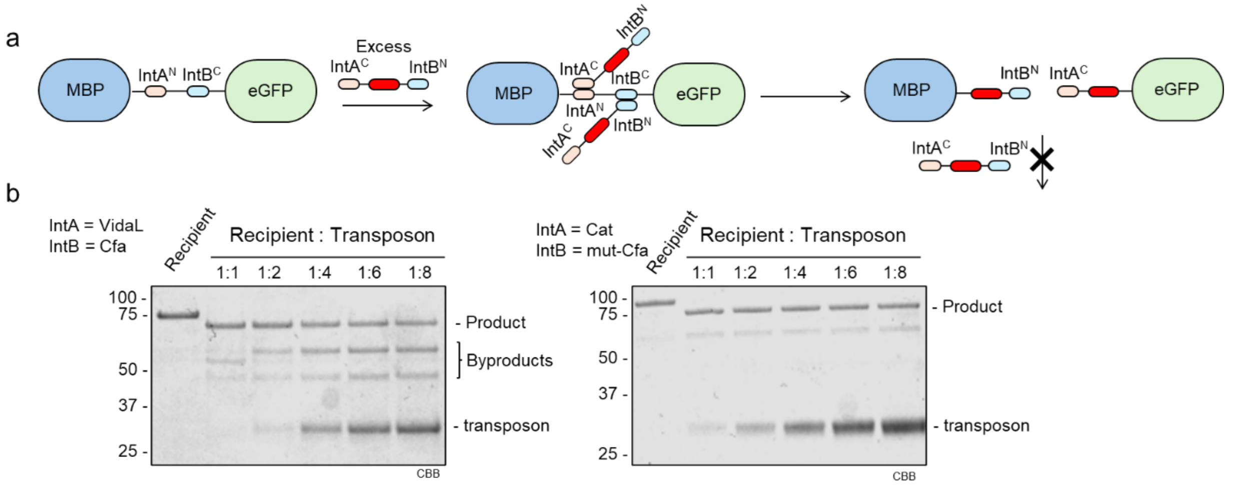


**Fig. S11. Effect of concentration mismatch between protein recipient and transposon.** (a). When an excess amount of protein transposon is present, two transposon proteins could bind to the same recipient and undergo two independent protein trans-splicing events, leading to the formation of two 'catalytically-dead' byproducts. If the recipient proteins are rapidly depleted by an excess of the protein transposons, then a buildup of these 'catalytically-dead' byproducts will be observed. (b) A comparison of product distribution for the split intein pairs VidaL/Cfa or Cat/mut-Cfa inserted into a MBP-eGFP protein recipient and a HA tag protein transposon. The concentration of recipients were fixed at 2 μM and the concentration of the transposon were 2, 4, 8, 12 and 16 μM, respectively. The transposition reaction was carried out in 100 mM phosphate, 150 mM NaCl, 1 mM EDTA, 1 mM TCEP, pH 7.2 for 2 hours at room temperature and the mixture was anlayzed by SDS-PAGE with Coomassie brilliant blue (CBB) staining. With the VidaL/Cfa pair, byproduct accumulation (MBP-HA-Cfa$^N$ and Vid$^C$-HA-eGFP) was observed when the transposon: recipient ratio is greater than 2. On the contrary, when using Cat/mut-Cfa pair, significant accumulation of the corresponding byproducts (MBP-HA-Cfa$^N$ and Cat$^C$-HA-eGFP) was not detected. These results show that Cat/mut-Cfa pair is less prone to concentration mismatch effects than the VidaL/Cfa pair, which can be helpful when the concentration of one of the two protein components cannot be measured accurately. For *in vitro* experiments, we highly recommended using a recipient to transposon ratio of 1:1 to minimize potential accumulation of byproducts.

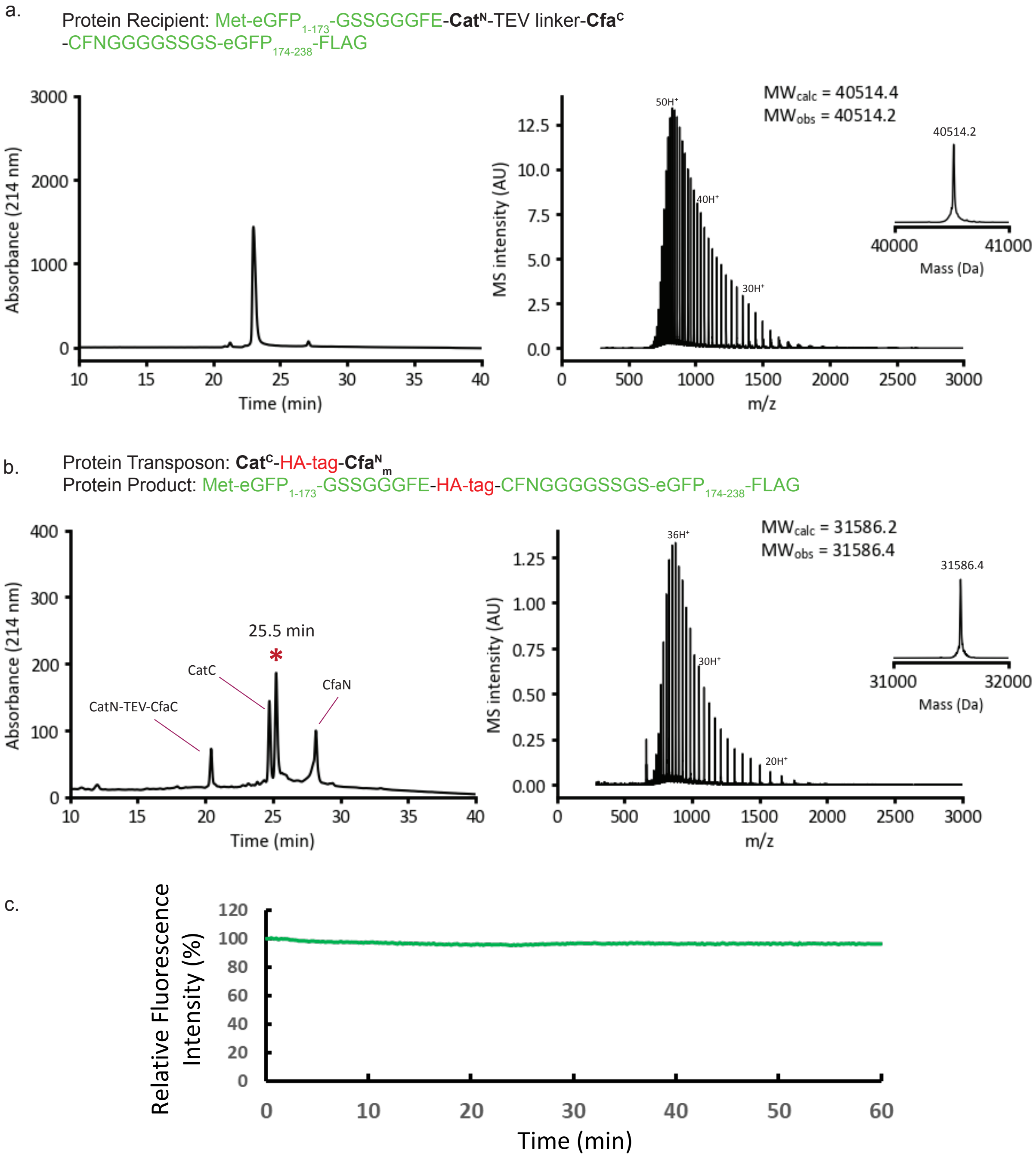


**Fig. S12. SDS-PAGE, ESI-MS and Fluorescence Spectroscopy Characterization of Protein Transposition on an eGFP Loop Using a HA Transposon Flanked by Cat$^{N}$ and Cfa$^{C}$ Split Inteins**. (a). The analytical C-18 RP-HPLC (gradient used: 20-75% B, 10-40 min) and ESI-MS analysis (deconvoluted spectrum in insert) of purified recipient eGFP. (b). Protein transposition was initiated by combining a HA tag transposon containing Cat$^{C}$ and Cfa$^{N}_{m}$ (2 μM) with the recipient eGFP protein (2 μM) in 100 mM phosphate, 150 mM NaCl, 1 mM EDTA, 1 mM TCEP, pH 7.2. The reaction was allowed to proceed at room temperature for 2 hours, The transposition product (red asterisk in the HPLC trace) was isolated by analytical C-18 RP-HPLC (gradient used: 20-75% B, 10-40 min) and characterized by ESI-MS analysis (deconvoluted spectrum in insert)

(c). The relative fluorescence intensity ($\lambda_{ex}$ = 488 nm, $\lambda_{em}$ = 509 nm) was monitored during the transposition reaction. Intensity is normalized to t =0.

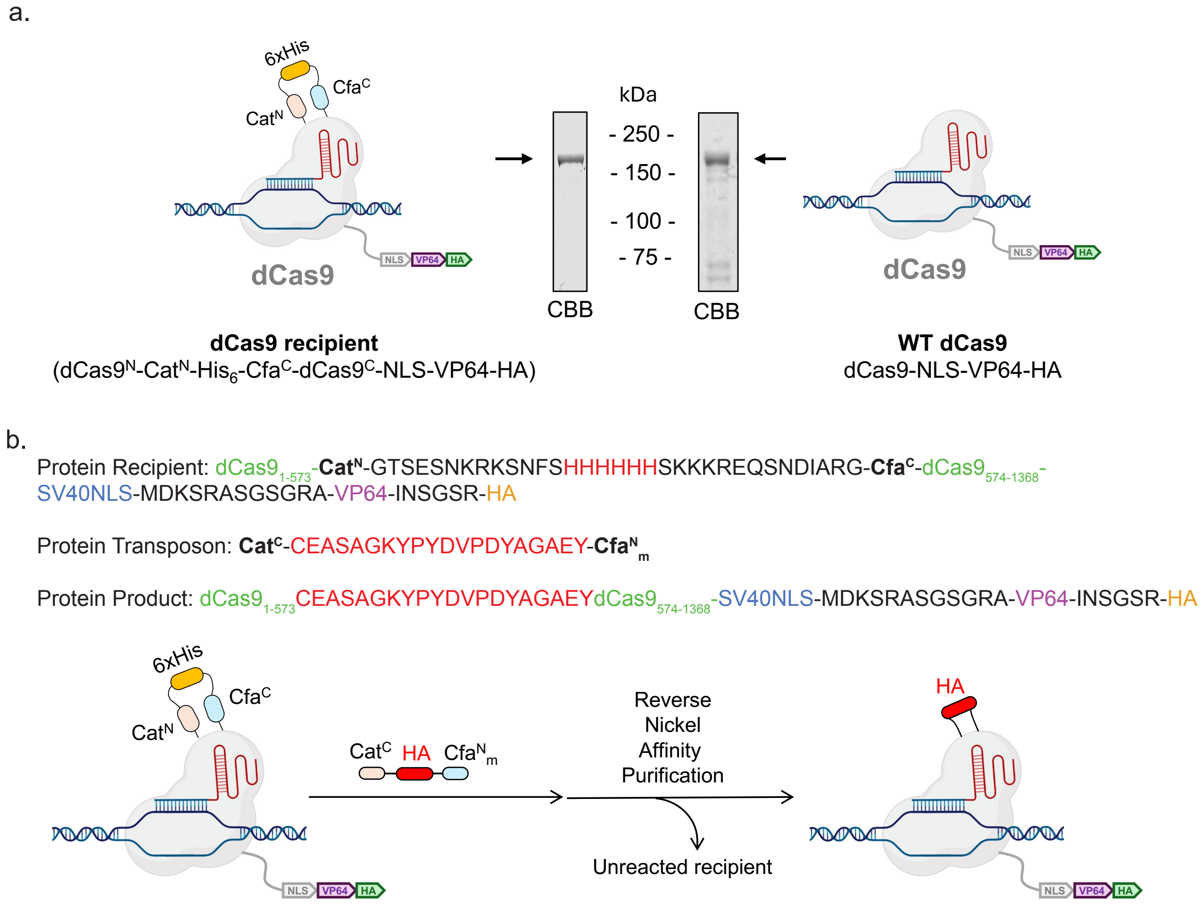


**Fig. S13. Additional details for dCas9 purification and expression** (a) SDS PAGE analysis (stained with Coomassie Brilliant Blue) of recombinant dcas9 recipient ($dCas9^N$-$Cat^N$-$His_6$-$Cfa^C$-$dCas9^C$-NLS-VP64-HA) and wild type dCas9 (dCas9-NLS-VP64-HA). (b) A representative example of the purification details for protein transposition using a dCas9 recipient protein and HA-protein transposon. The 6x His tag incorporated between the two split intein fragments on the recipient dCas9 enables nickel affinity purification as the 6x His tag is removed post-transposition, and a reverse nickel affinity purification step allows for removal of unreacted recipient, demonstrating a robust strategy for further purification of transposition products. This approach can be extended to other transposons.

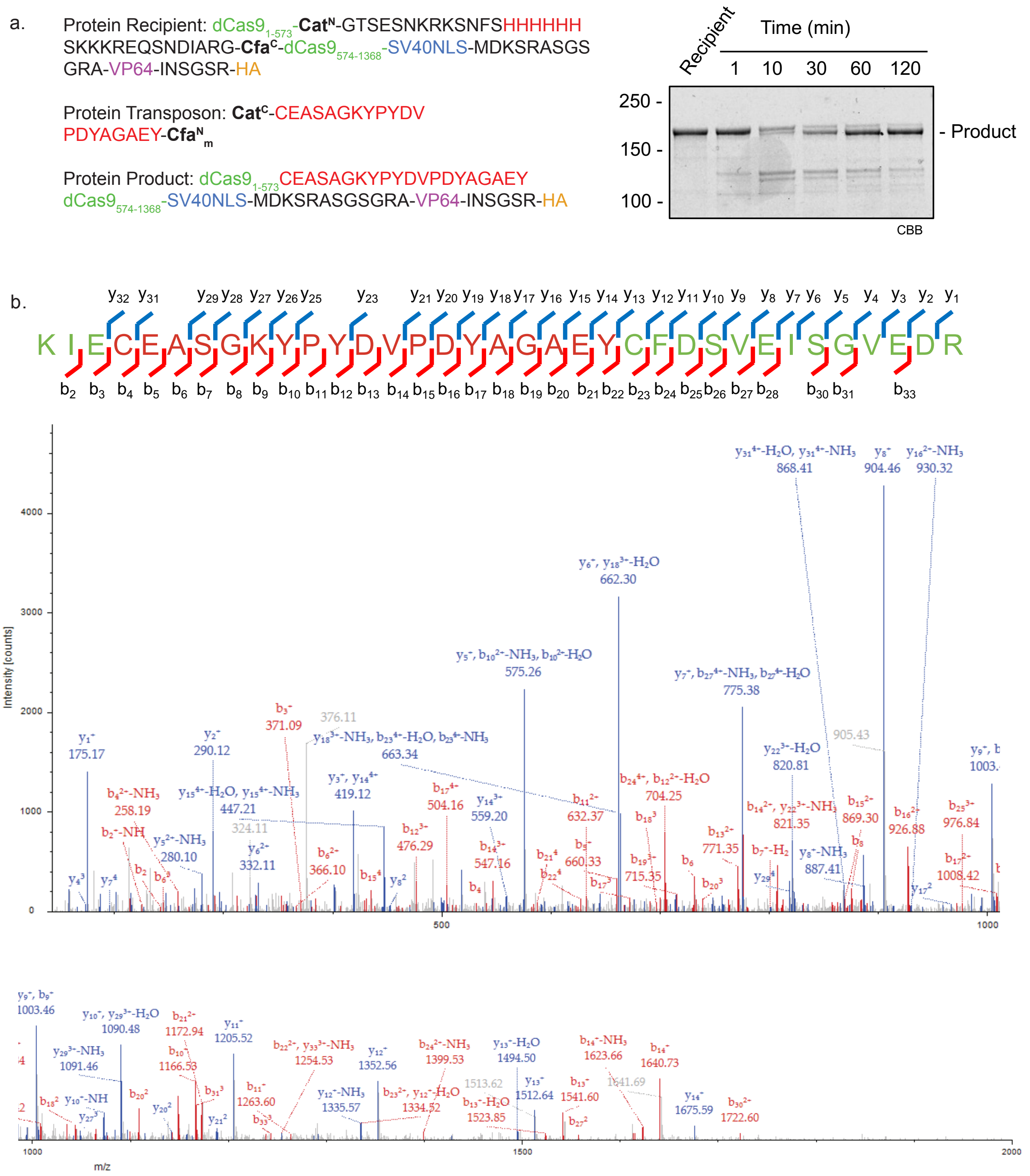


**Fig. S14. LC-MS/MS Characterization of Protein Transposition on dCas9 using a HA Transposon flanked by Cat$^{C}$ and mut-Cfa$^{N}$ Split Inteins.** (a) The transposition reaction (1.5 µM) was carried out at room temperature for 2 hours in 50 mM Tris, 250 mM NaCl, 1 mM TCEP, 10% v/v glycerol, pH 7.5. The reaction progress was monitored and characterized by SDS-PAGE with Coomassie brilliant blue (CBB) staining. (b). The gel band containing the product was excised and digested with trypsin, followed by LC-MS/MS analysis. See methods for details. The peptide containing both splicing junctions was identified and characterized.

#### General strategy for protein semi-synthesis via sequential EPL

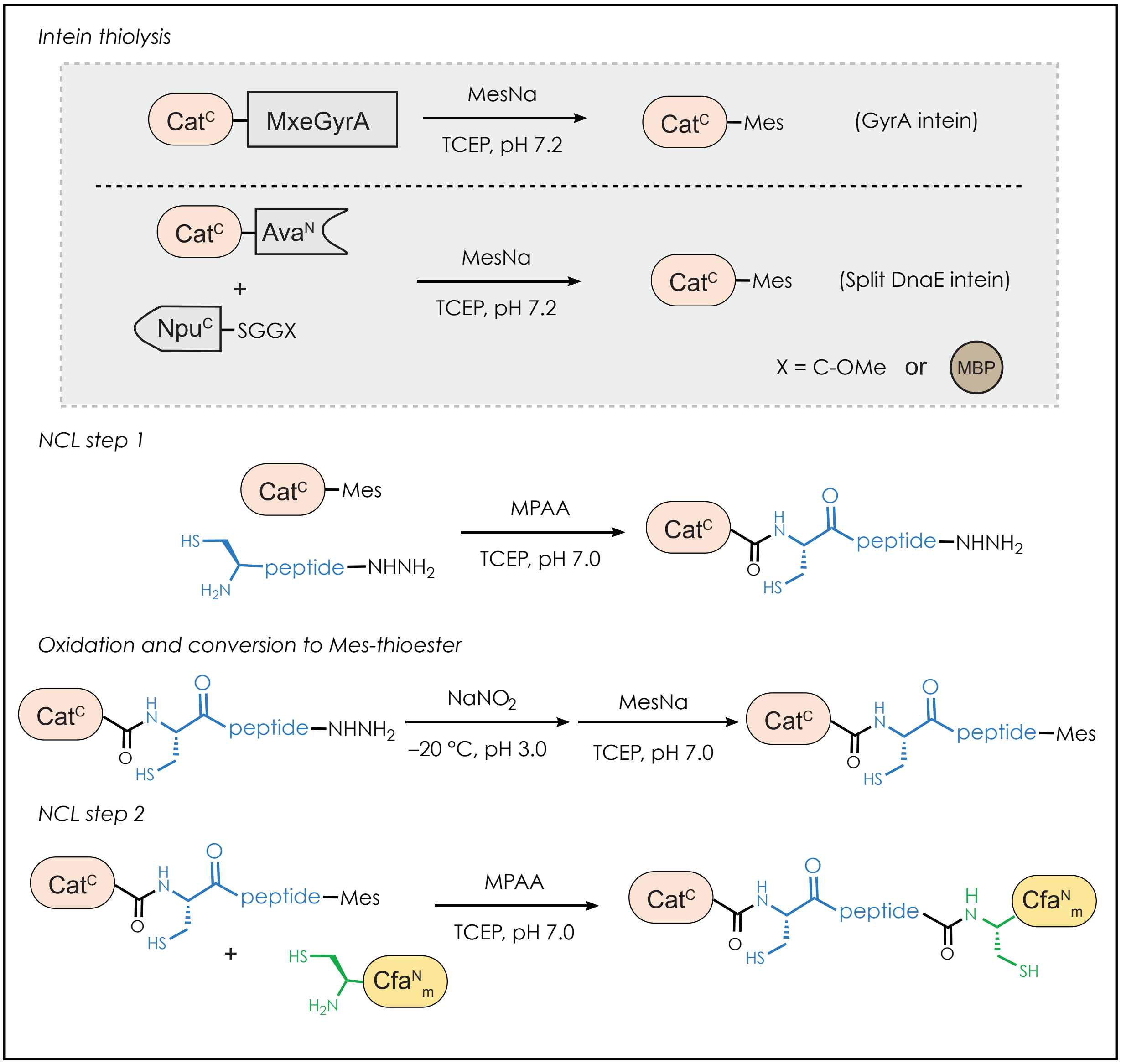


**Fig. S15. General strategy for the semisynthesis of non-native peptide transposons used in this study.** Transposons contain a functional peptide motif flanked by orthogonal split inteins. The Cat$^{C}$-Mes thioester is obtained either by MesNa thiolysis of a recombinant C-terminal MxeGyrA intein or a recombinant C-terminal Ava$^{N}$ split intein in the presence of catalytically inactive Npu$^{C}$. The Cat$^{C}$-Mes thioester is then condensed with the functional peptide containing a C-terminal acyl hydrazide through Native Chemical Ligation (NCL). The purified NCL product is then oxidized with $NaNO_2$ to the acyl azide, which was then trapped with MesNa to yield the CatC-peptide-Mes thioester. This intermediate was then ligated with recombinant Cfa$^{N}_{m}$ to yield the final protein product.

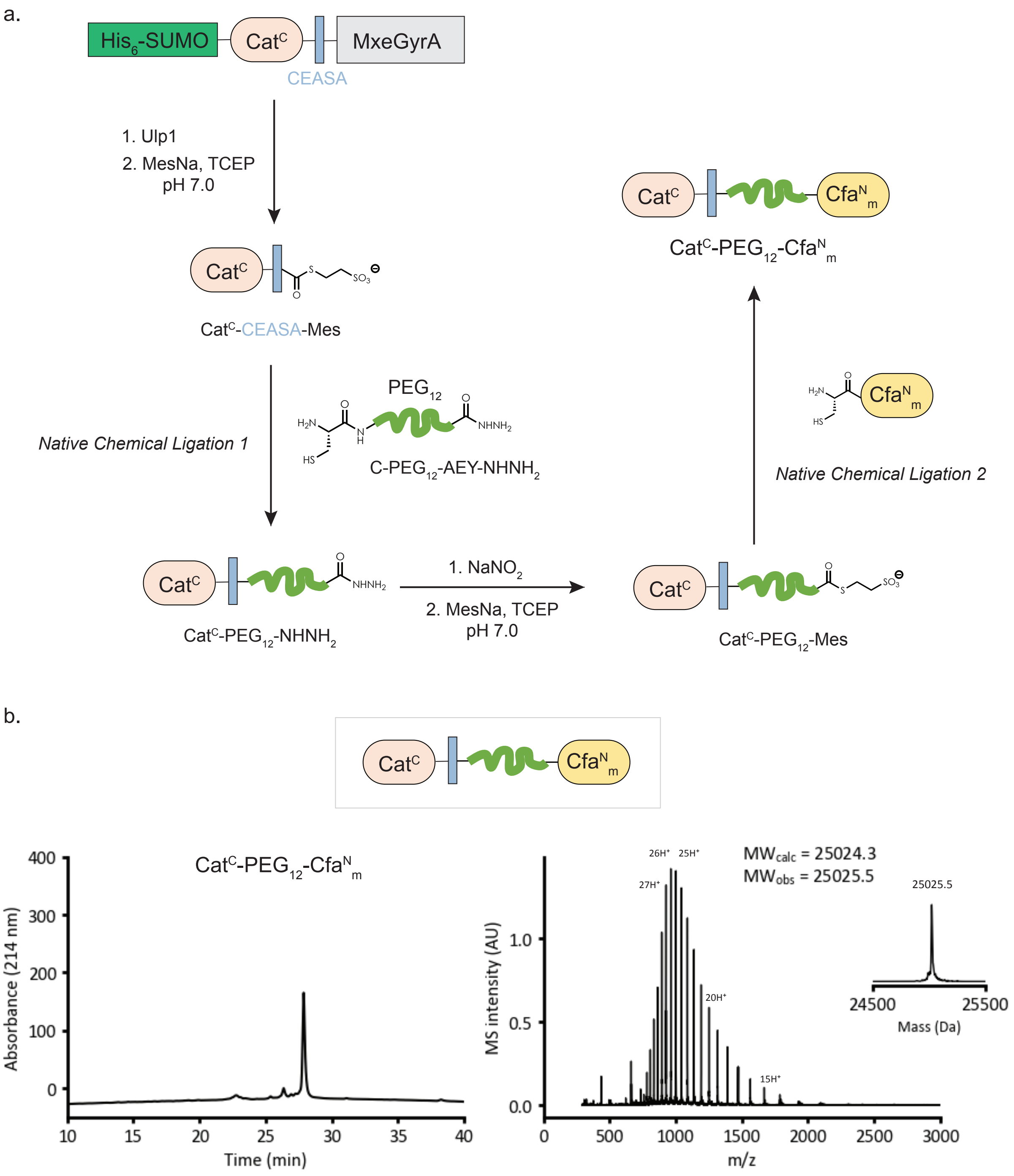


**Fig. S16. Scheme for the Semisynthesis of Cat$^{C}$-CEASAC-PEG$_{12}$-AEY-Cfa$^{N}_{m}$ from Three Separate Peptides.** (a) The Cat$^{C}$-CEASA-Mes thioester was obtained from Ulp1 cleavage and MesNa thiolysis of recombinant His$_6$-SUMO-Cat$^{C}$-CEASA-MxeGyrA and subsequently condensed with C-PEG$_{12}$-AEY-NHNH$_2$ peptide through Native Chemical Ligation (NCL). The purified NCL product (Cat$^{C}$-PEG$_{12}$-AEY-NHNH$_2$) was then oxidized with $NaNO_2$ to the acyl azide, which was then trapped with MesNa to yield the Cat$^{C}$-PEG$_{12}$-AEY-Mes thioester. This intermediate was then ligated with recombinant Cfa$^{N}_{m}$ to yield the final protein product (Cat$^{C}$-

$PEG_{12}$-$Cfa^{N}_{m}$). See methods for reaction conditions. (b) Analytical C-18 RP-HPLC chromatogram (gradient used: 20-75% B, 10-40 min) (left) and ESI-MS (deconvoluted spectra shown in insert)

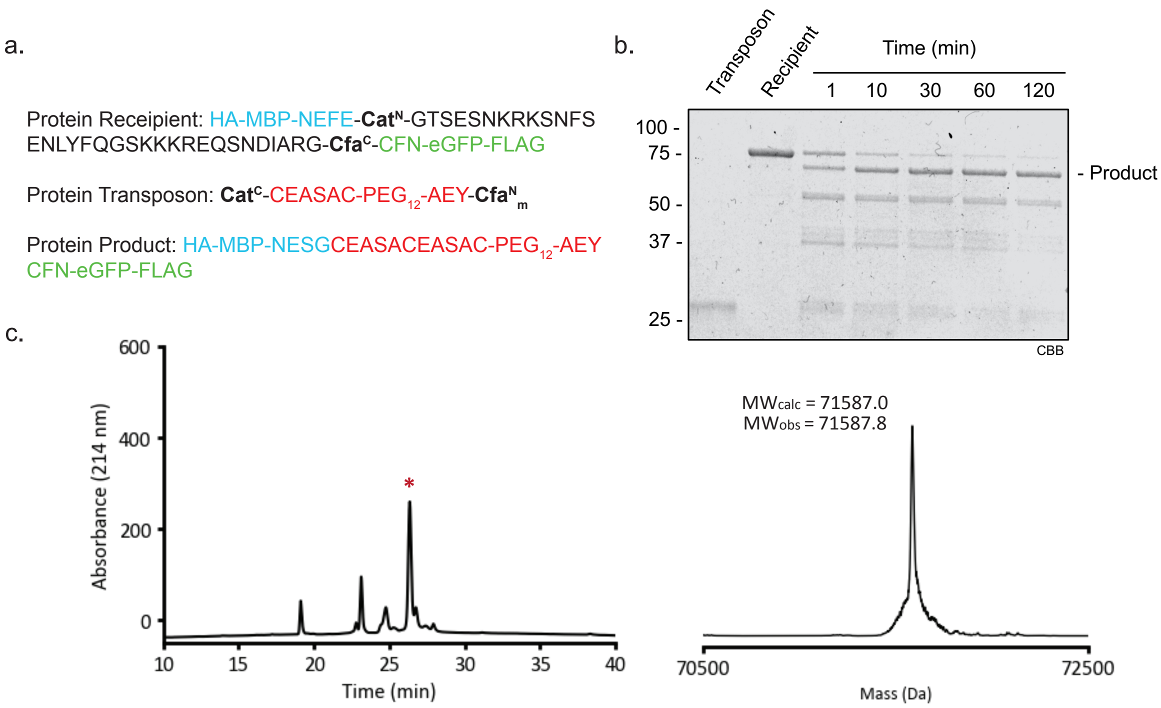


**Fig. S17. Characterization of Transposition on a Model MBP-eGFP Fusion Protein with a $PEG_{12}$-containing transposon by RP-HPLC and ESI-MS** (a). The transposition using $PEG_{12}$-protein chimera transposon was first tested on MBP-eGFP model recipient using a HA transposon flanked by $Cat^{C}$ and mut-$Cfa^{N}$ split inteins. (b). The transposition reaction (2 µM) was carried out at room temperature for 2 hours and the reaction progress was monitored and characterized by SDS-PAGE. (c). The transposition product (red asterisk) was also isolated by analytical C-18 RP-HPLC (gradient used: 20-75% B, 10-40 min) (left) and confirmed by ESI-MS (right).

#### SMARCA5 APB1 Transposon

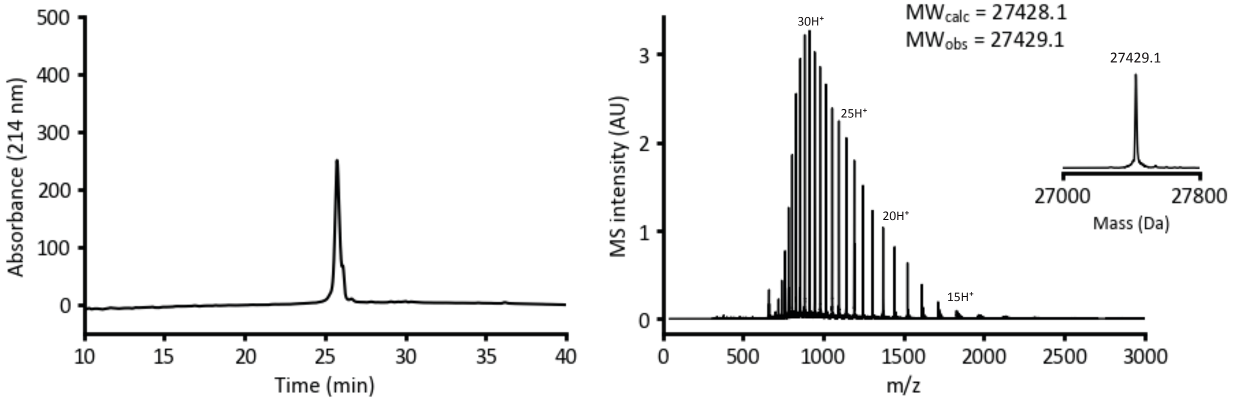


#### SMARCA5 APB2 Transposon

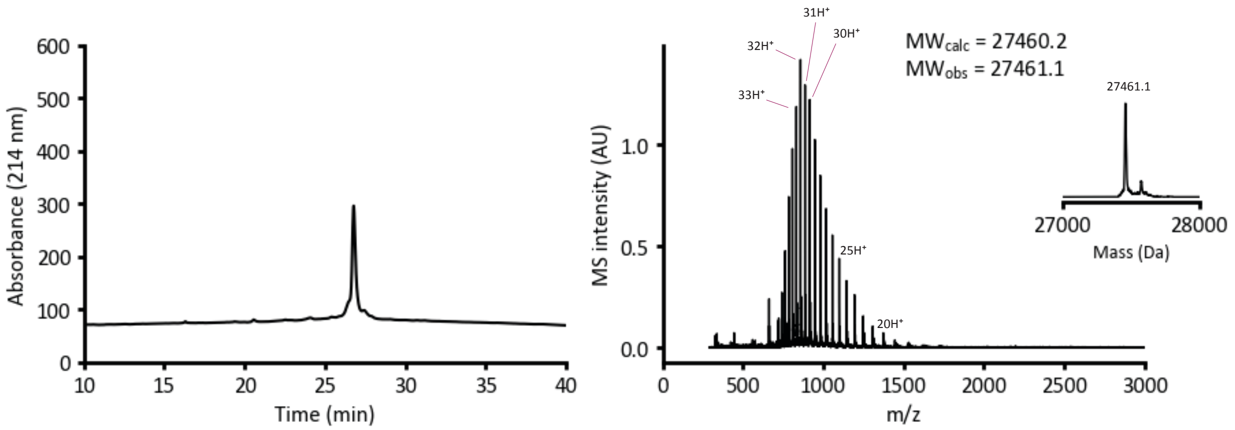


**Fig. S18. Characterization of SMARCA5 APB1 and APB2 Transposons by RP-HPLC and ESI-MS.** Top: Analytical C-18 RP-HPLC chromatogram (gradient used: 20-75% B, 10-40 min) (left) for SMARCA5 APB1 transposon and the corresponding ESI-MS spectra with deconvoluted spectra shown in insert (right). Bottom: Analytical C-18 RP-HPLC chromatogram (gradient used: 20-75% B, 10-40 min) (left) for SMARCA5 APB2 transposon and the corresponding ESI-MS spectra with deconvoluted spectra shown in insert (right).

a.

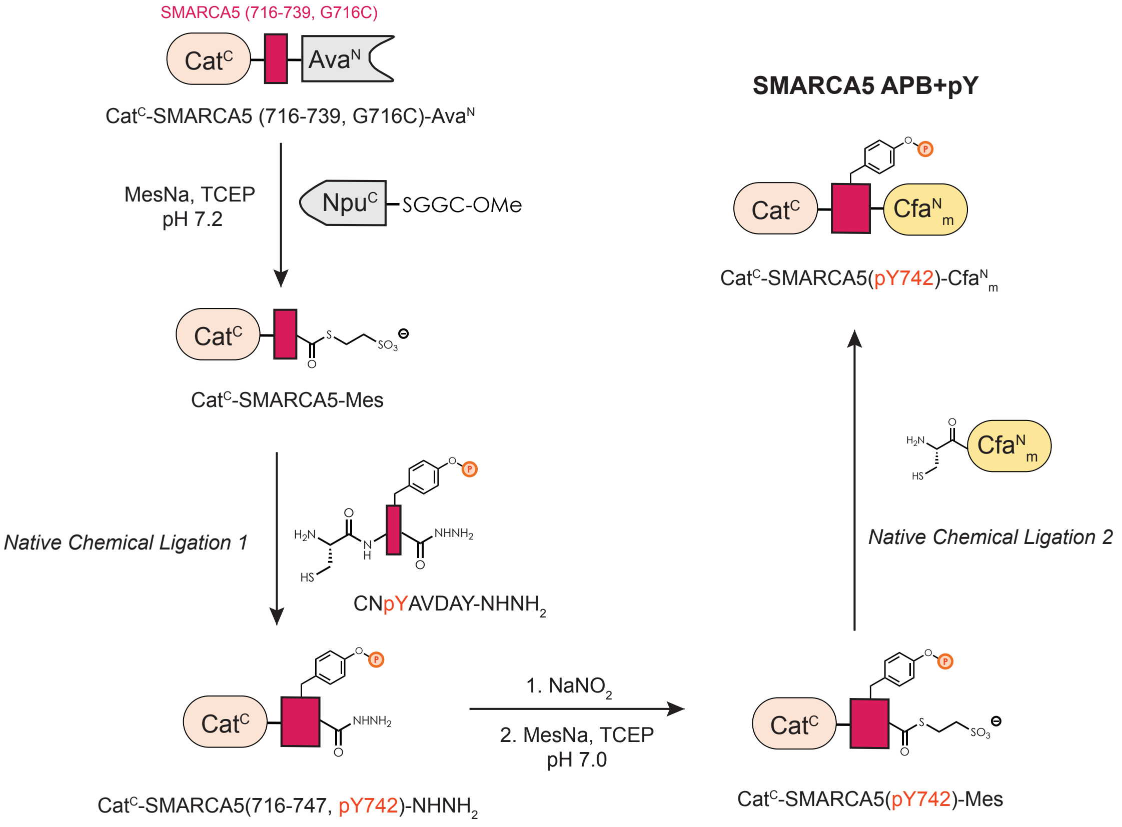


b.

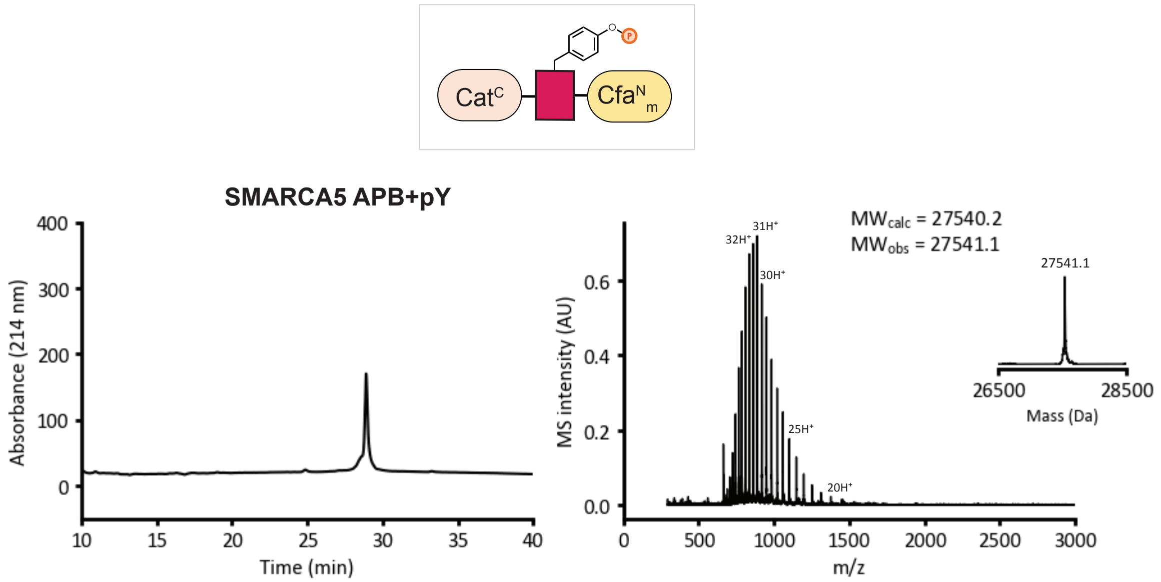


**Fig. S19. Scheme for the Semisynthesis of SMARCA5 APB+pY Transposon**. (a) The Cat$^{C}$-SMARCA5-Mes thioester was obtained via MesNa thiolysis of recombinant Cat$^{C}$-SMARCA5(716-747, G716C)-Ava$^{N}$ in the presence of Npu$^{C}$-SGGCOMe and subsequently condensed with CNpYAVDAY-$NHNH_2$ peptide through Native Chemical Ligation (NCL). The purified NCL product (Cat$^{C}$-SMARCA5(716-747, pY742)-$NHNH_2$) was then oxidized with $NaNO_2$ to the acyl azide, which was then trapped with MesNa to yield the Cat$^{C}$-

SMARCA5(pY742)-Mes thioester. This intermediate was then ligated with recombinant $Cfa^{N}_{m}$ to yield the final protein product (SMARCA5 APB+pY transposon). See methods for reaction conditions. (b) Analytical C-18 RP-HPLC chromatogram (gradient used: 20-75% B, 10-40 min) (left) and ESI-MS (deconvoluted spectra shown in insert) of the product.

a.

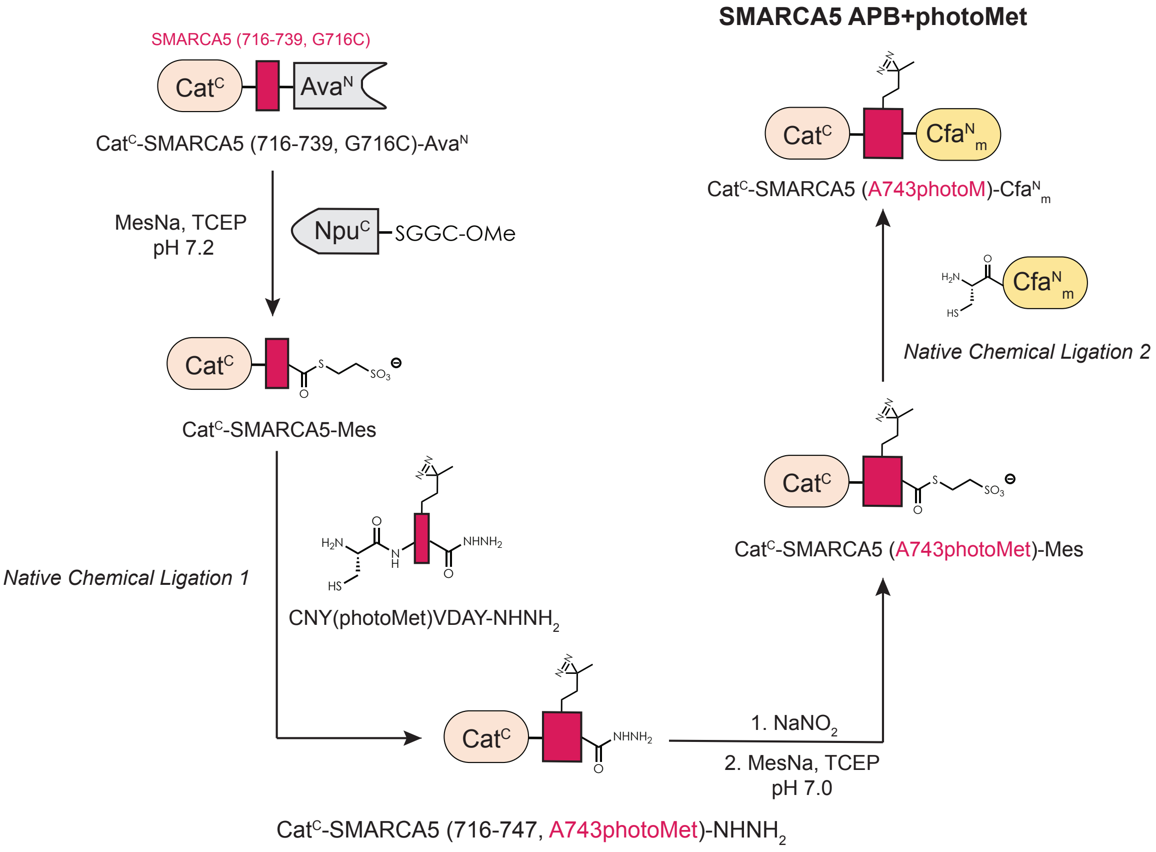


b.

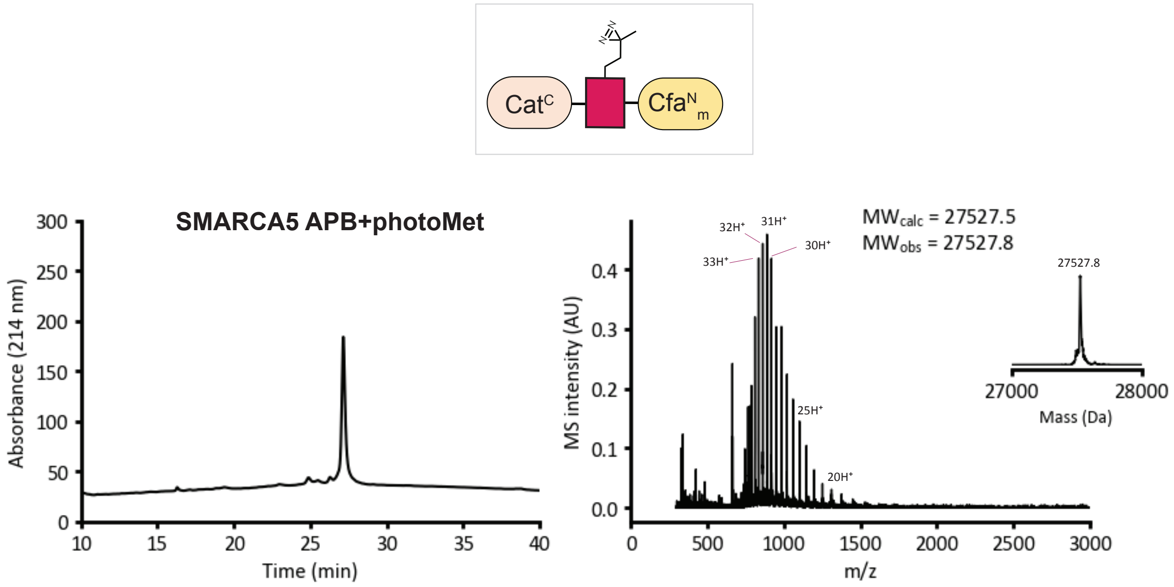

**Fig. S20. Scheme for the Semisynthesis of SMARCA5 APB+photoMet Transposon**. (a) The $Cat^{C}$-SMARCA5-Mes thioester was obtained via MesNa thiolysis of recombinant $Cat^{C}$-SMARCA5(716-747, G716C)-$Ava^{N}$ in the presence of $Npu^{C}$-SGGCOMe and subsequently condensed with CNY(photoMet)VDAY-$NHNH_2$ peptide through Native Chemical Ligation (NCL). The purified NCL product ($Cat^{C}$-SMARCA5(716-747, A743photoMet)-$NHNH_2$) was then oxidized with $NaNO_2$ to the acyl azide, which was then trapped with MesNa to yield the $Cat^{C}$-SMARCA5(A743photoMet)-Mes thioester. This intermediate was then ligated with recombinant $Cfa^{N}_{m}$ to yield the final protein product (SMARCA5 APB+photoMet transposon). See methods for reaction conditions. (b) Analytical C-18 RP-HPLC chromatogram (gradient used: 20-75% B, 10-40 min) (left) and ESI-MS (deconvoluted spectra shown in insert) of the product.

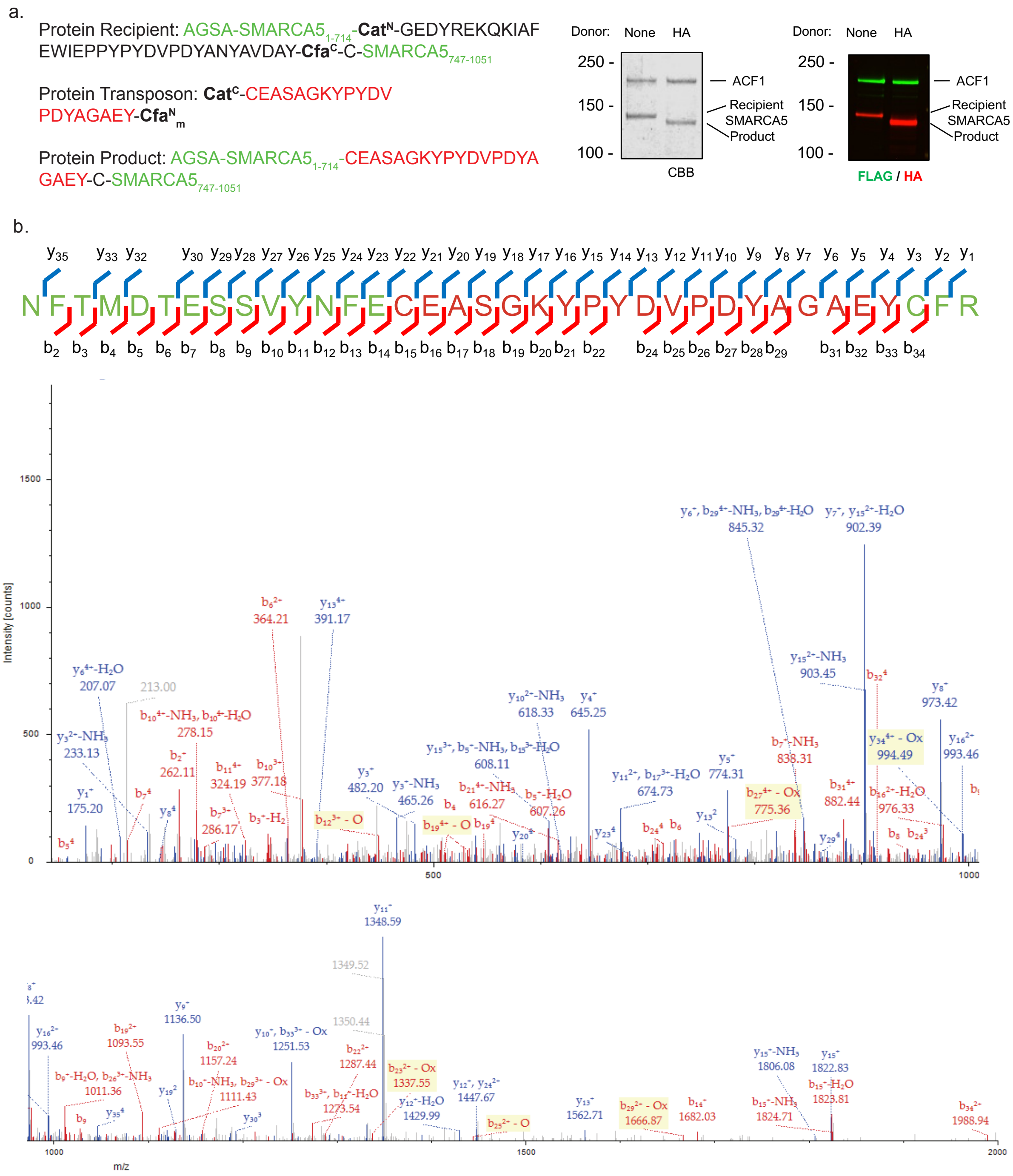


**Fig. S21. LC-MS/MS Characterization of Protein Transposition on SMARCA5 as a Subunit of the ACF Complex Using a HA Transposon Flanked by Cat$^C$ and mut-Cfa$^N$ Split Inteins.** The transposition reaction (500 nM) was carried out at room temperature for 2 hours in ACF transposition buffer (25 mM HEPES, 60 mM KCl, 10 mM $MgCl_2$, 1 mM TCEP, 10% v/v glycerol, 0.02% v/v IPEGAL CA-630). The reaction product was characterized by SDS-PAGE. (b). The gel band containing the product was excised and digested by trypsin, followed by LC-MS/MS analysis. See methods for details. The peptide containing both splicing junctions was identified and characterized.

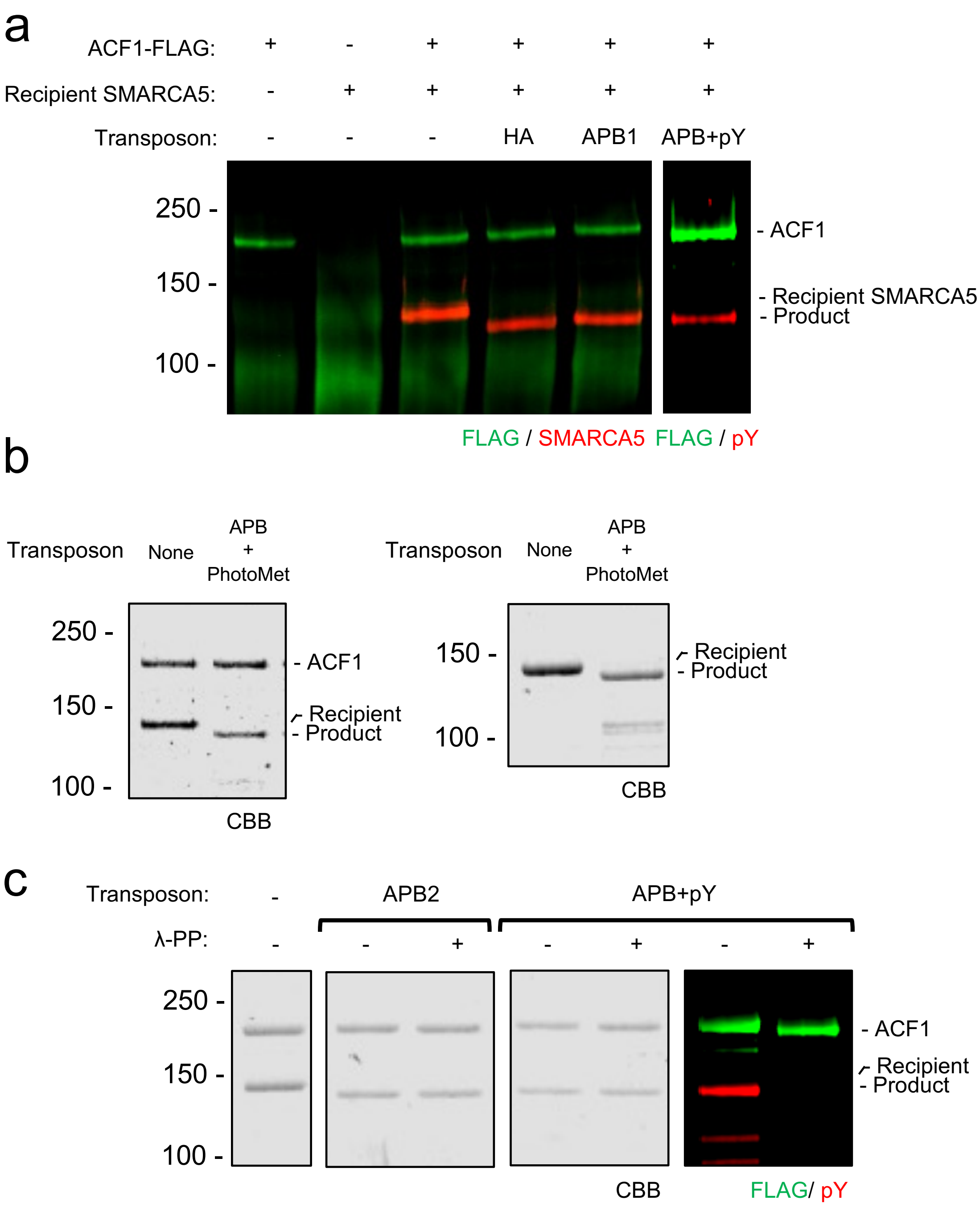


**Fig. S22. Analysis of protein transposition on ACF complex and SMARCA5.** (a). Result of immunoprecipitation of ACF1-FLAG using anti-FLAG G1 resin. 50 µL of each protein or complex (all at 500 nM) was diluted with ACF transposition buffer and then incubated with 20 µL of anti-FLAG G1 resin; after washing, the bound proteins or complexes were eluted by 50 µL of FLAG peptide. The different eluents were characterized by SDS-PAGE and western blotting (visualized by the indicted antibodies). When ACF1-FLAG is absent, SMARCA5 signal was not detectable; whereas when ACF1-FLAG is present, both ACF1-FLAG and SMARCA5 were observed. These results have shown that the recipient SMARCA5 and all of the transposed SMARCA proteins are capable of forming the ACF complex with ACF1-FLAG. The APB transposons used in this experiment are as follows: $Cat^{C}$-HA-$Cfa^{N}_{m}$ (HA), SMARCA5 APB1 (APB), and SMARCA5 APB+pY (APB + pY) (b). Transposition products using recipient SMARCA5 and different transposons used for the photo-crosslinking experiments. For validation of ACF activity, the recipient ACF complex and the photoMet transposon (both at 500 nM) were reacted in ACF transposition buffer at 4 °C overnight. For the designer SMARCA5 used for the photo-crosslinking experiment, the recipient SMARCA5 and the respective transposon (both at 1 μM) were reacted in ACF transposition buffer at 4C° overnight. The products were characterized by SDS-PAGE with Coomassie staining. The APB transposon used in this experiment SMARCA5 APB+photoMet (APB+photoMet). (c). SDS-PAGE and western blotting analysis of protein transposition on ACF complex with various transposons as well as λ-phosphatase (λ-PP) treated ACF complexes used in nucleosome remodeling assays. the recipient ACF complex and the respective transposon (both at 500 nM) were reacted in ACF transposition buffer at 4 °C overnight. The products were characterized by SDS-PAGE with Coomassie staining and western blotting with respective antibodies. The APB transposons used in this experiment are as follows: SMARCA5 APB2 (APB2) and SMARCA5 APB+pY (APB + pY). APB2 and APB+pY differ only by the pTyr742 modification.

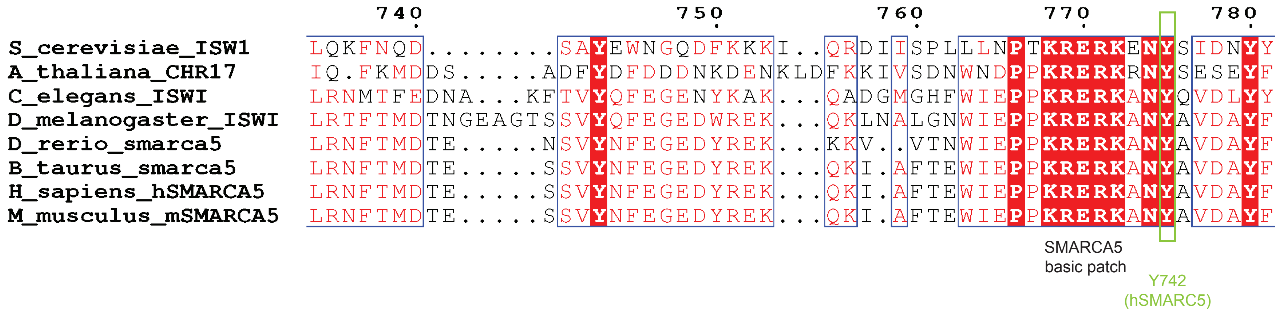


**Fig. S23.** Alignment of ISWI/SMARCA5 orthologs reveals a conserved basic motif (KRERK) with a proximal conserved tyrosine (green box, Y742 in hSMARCA5)

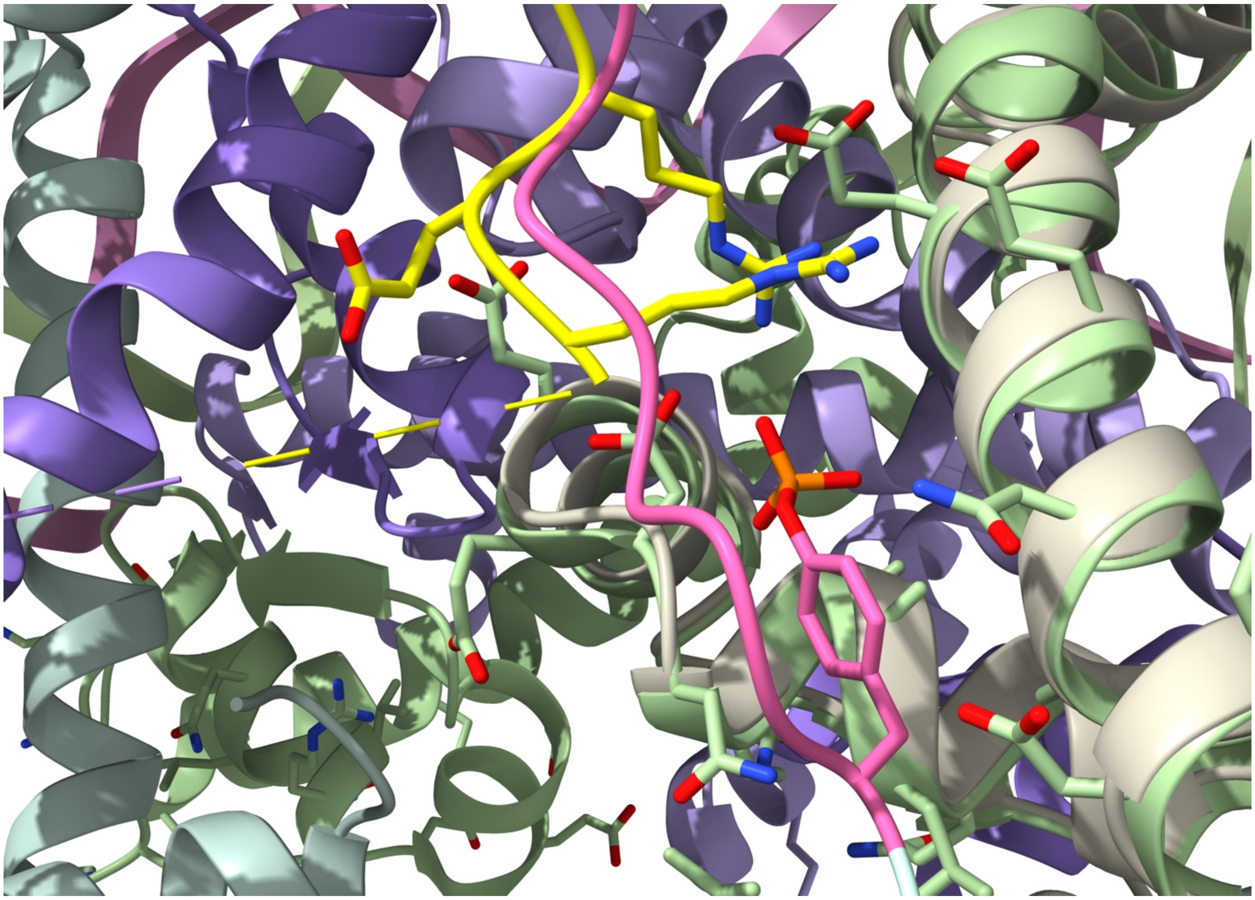

**Fig. S24. Computational Modeling of a APB-Proximal Phosphotyrosine PTM Within the IWS1-Nucleosome Complex.** An overlay of an AlphaFold3-derived ISWIa(pTyr775)-H2A interaction (pink and white respectively) with the cryo-EM structure of the IWSIa-nucleosome complex (PDB 6JYL) (yellow ISW1a, green H2A) shows that the APB-proximal phosphotyrosine (pink-orange) is positioned to interact with the key arginines of the basic APB motif (yellow). The phosphotyrosine residue was modeled using Vienna-PTM 2.0. pTyr775 in ISW1a maps onto pTyr742 in hSMARCA5.

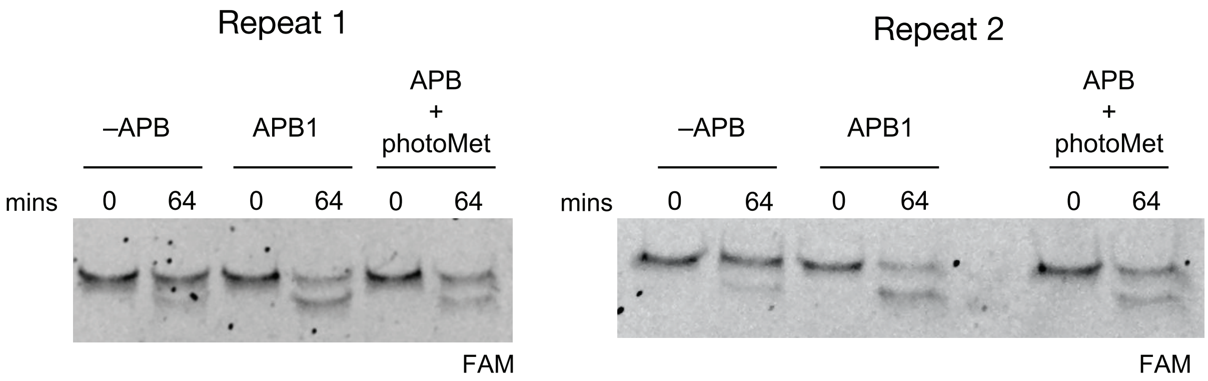


**Fig. S25. REAA Remodeling Assays Show that Photomethionine Incorporation into SMARCA5 Does not Ablate ACF Remodeling.** Transposed ACF containing the photocrosslinker (photoM743) is competent for remodeling, as assayed by restriction enzyme accessibility assay (REAA). Activity is compared to -APB and APB1, which are also shown in Fig. 4C. In all cases REAA is performed with 10 nM ACF and 10 nM MNs (FAM-labeled, 45 and 15 bp DNA overhangs).

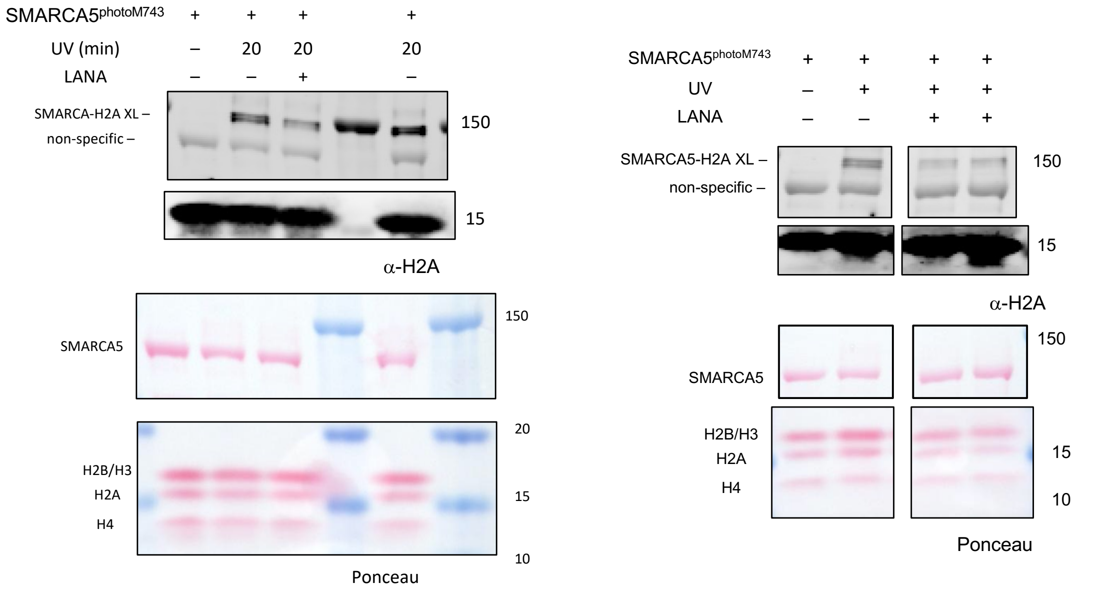


Crosslinking conditions: Nucleosomes (with 45/15 bp overhang) (5 pmol), SMARCA5 (10 pmol), LANA peptide 10 uM. UV 20 min at 4 °C

**Fig. S26. UV-Crosslinking of a SMARCA5 Photo-Methionine to a Mononucleosome Substrate shows Covalent Attachment to Histone H2A.** Nucleosomes were incubated with SMARCA5 containing the photoMet probe at position 743 either with or without LANA peptide and UV irradiation. Mixtures were analyzed by western blot using Ponceau staining (loading) and anti-H2A antibody (ab18255). A UV-dependent (+/-) crosslink was observed for the sample in the absence of LANA peptide, and the addition of LANA peptide significantly decreases the SMARCA5-H2A crosslink. Repeats are shown here.

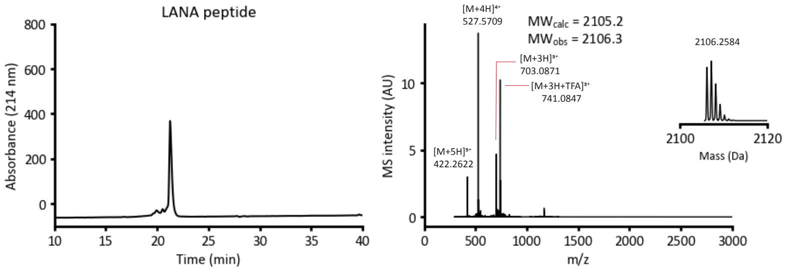


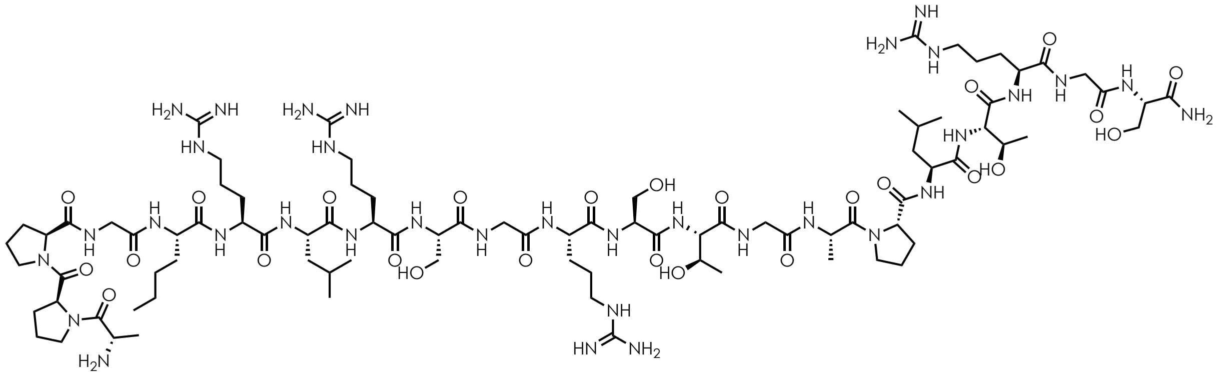

$NH_2$-APPG(Nle)RLRSGRSTGAPLTRGS-$NH_2$

**Fig. S27.** Analytical HPLC trace and ESI-TOF MS analyses (deconvoluted spectra in insert) for the LANA peptide used in the photocrosslinking experiment (Fig. 4E and Fig. S26).

**Cat$^{C}$-FLAG-Cfa$^{N}_{m}$** (Cat$^{C}$-CEASGKDYKDDDDKGAEY-Cfa$^{N}_{m}$

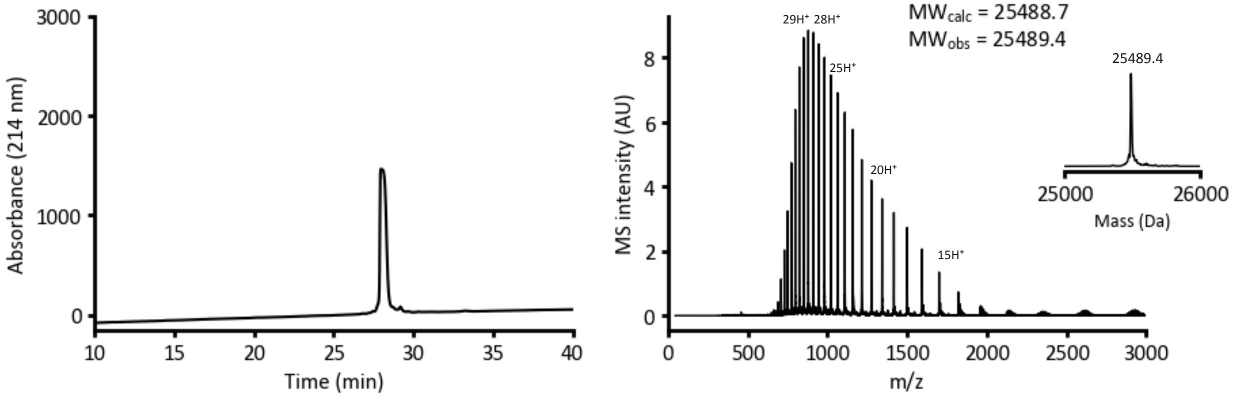


**Fig. S28. Characterization of SMARCA5 Transposon for *In Nucleo* Splicing by RP-HPLC and ESI-MS.** Top: Analytical C-18 RP-HPLC chromatogram (gradient used: 20-75% B, 10-40 min) (left) for the transposon containing Cat$^{C}$ and Cfa$^{N}_{m}$ split inteins flanking the peptide corresponding to SMARCA5 with an embedded FLAG tag (red) and the corresponding ESI-MS spectra with deconvoluted spectra shown in insert (right).

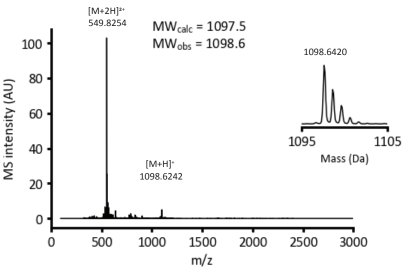


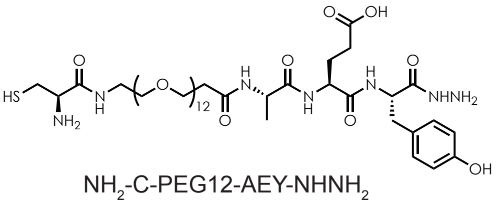

NH$_2$-C-PEG12-AEY-NHNH$_2$

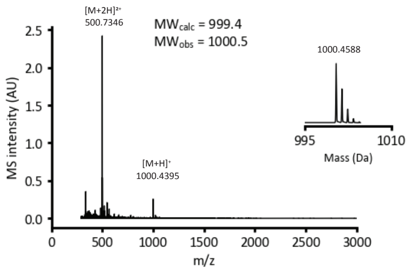


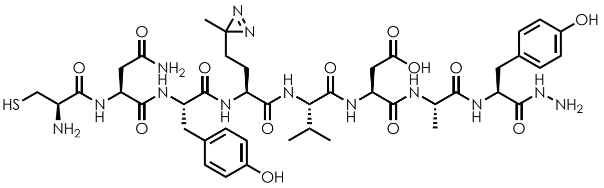

NH$_2$-CNYM(photo)VDAY-NHNH$_2$

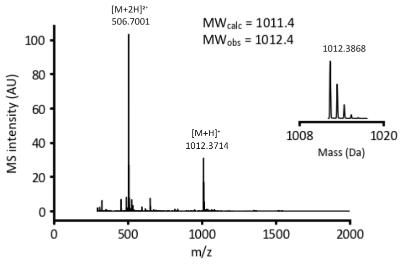


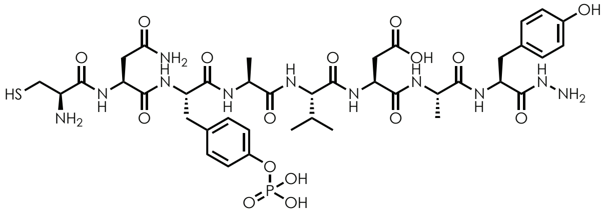

NH$_2$-CNpYAVDAY-NHNH$_2$

**Fig. S29** ESI-TOF MS analyses (deconvoluted spectra in insert) for the peptides used in Figs. S16, S19, and S20.

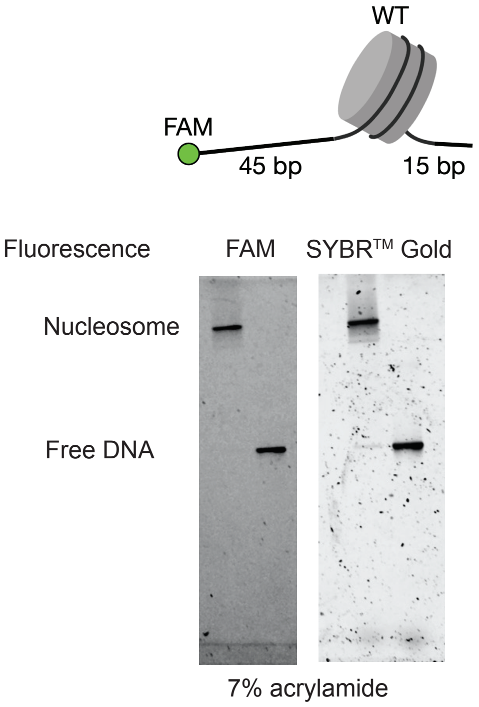


**Fig. S30.** SDS-PAGE of nucleosomes containing histones H2A, H2B, H3.1, H4 with 207 bp DNA base pair with 45/15 overhangs used in remodeling and photo crosslinking studies.

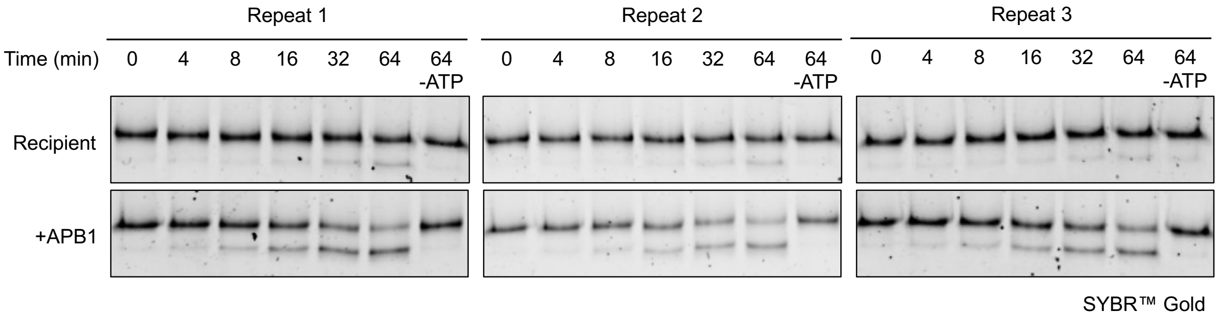


**Fig. S31 Reintroducing the APB domain into the recipient SMARCA5 in an ACF complex rescues its remodeling activity.** Transposed ACF containing the APB domain regained its remodeling activity comparing to the recipient ACF, which lacks the APB domain. The remodeling activity of the two ACF complexes were assayed by restriction enzyme accessibility assay (REAA) and the results at indicated times were further subjected to kinetic analysis as shown in Fig. 4C. In all cases REAA is performed with 10 nM ACF and 10 nM MNs (45 and 15 bp DNA overhangs). The gels were visualized by SYBR™ Gold staining.

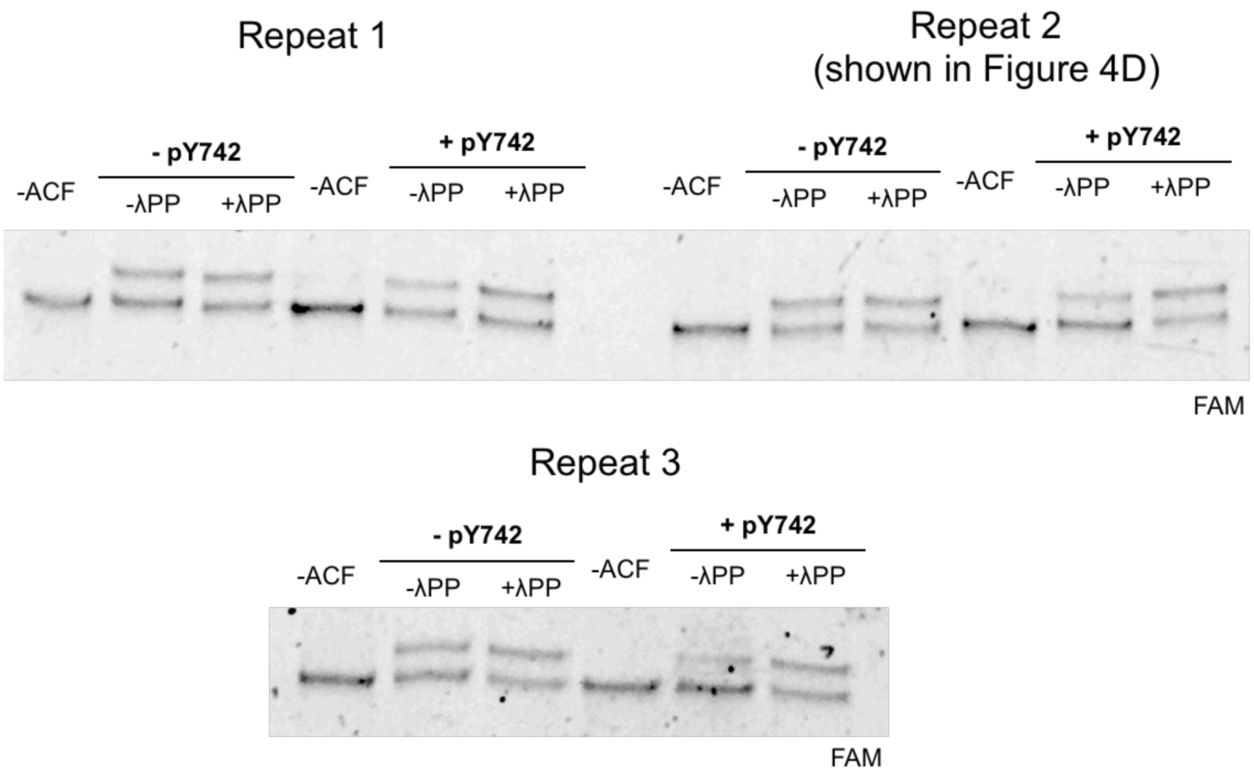


**Fig. S32.** Three repeats of the EMSA remodeling assay (8.33 nM ACF, 10 nM MNs, 4 minutes). used to determine the effects of pY742 on ACF activity. Repeat 2 is shown in Fig. 4D. All three repeats quantified by densitometry for the bar chart shown in Fig. 4D.

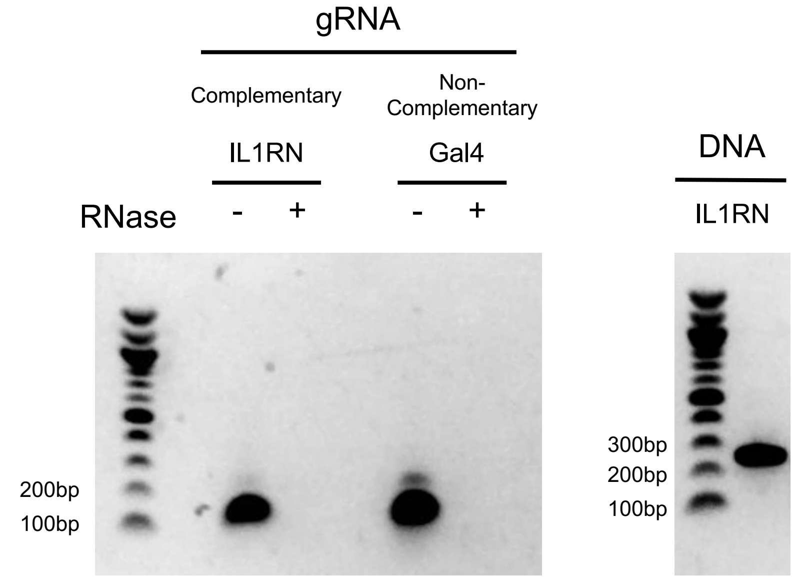


**Fig. S33. Validation of RNA and DNA products used in the dCas9 binding assay. Left**: A representative agarose gel (1.2%) showing analysis of in vitro transcribed complementary (IL1RN) and non-complementary (Gal4) gRNA constructs (left). **Right:** A representative agarose gel (1.2%) showing the DNA product (a fragment from IL1RN promoter) used in the in-vitro dCas9:gRNA:DNA binding gel shift assay (Fig. 3F).

**$Cfa^{N}{}_{m}$ ($Cfa^{C}$ M75L M81L)**

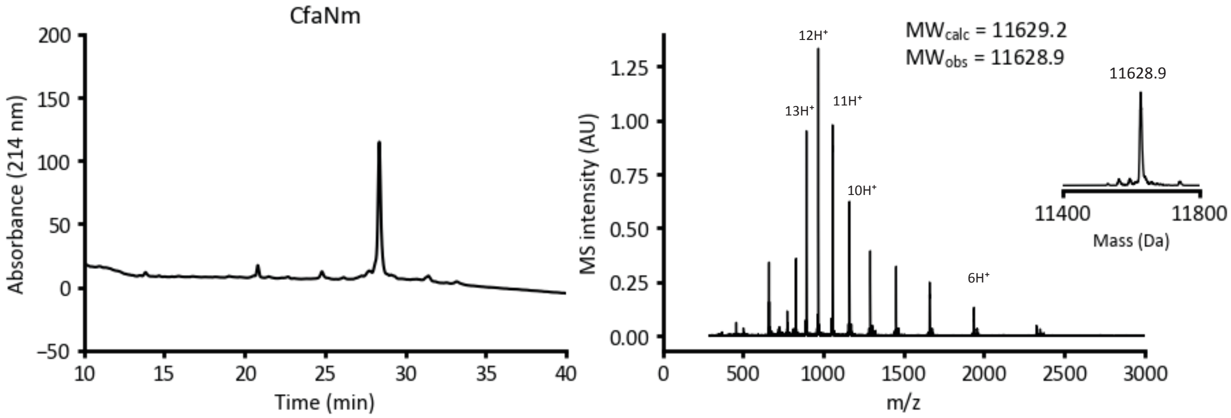


**$Npu^{C}$(N35A)-AFNSGG-MBP**

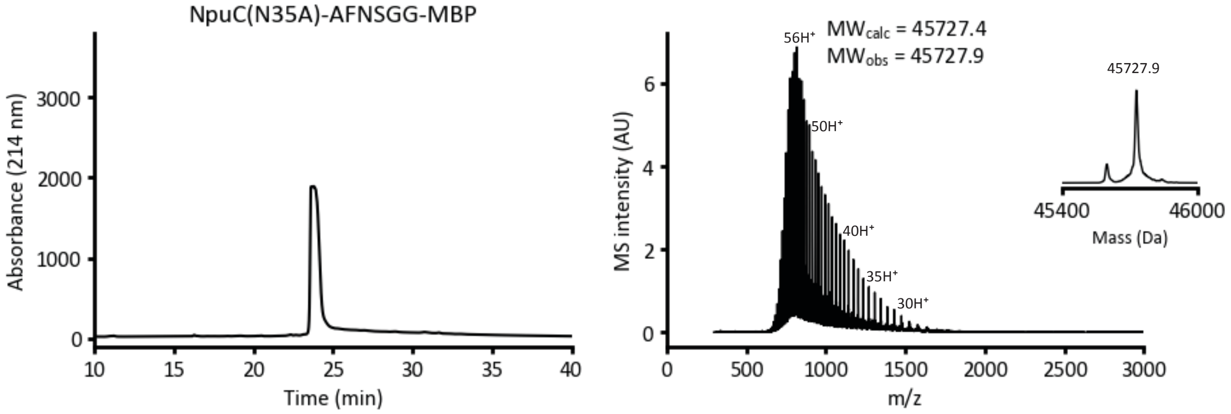


**$Npu^{C}$(N35A)-AFNSGG-COMe**

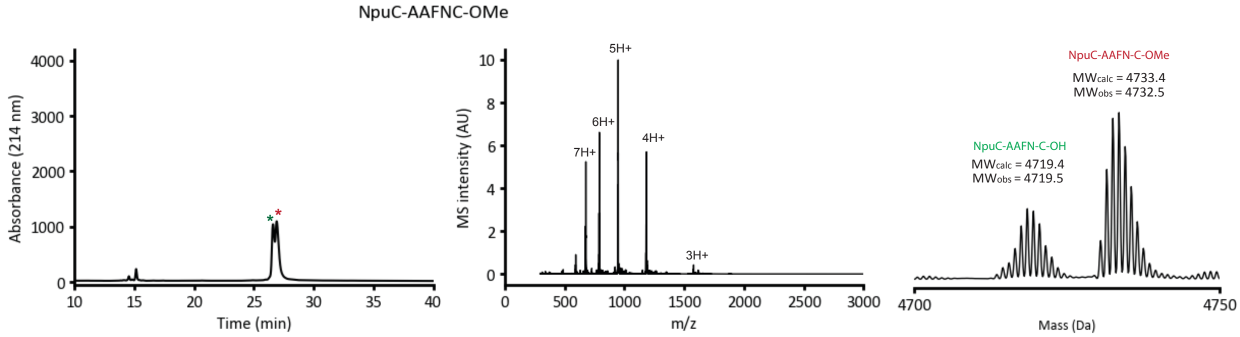


**Fig. S34. Characterization of Miscellaneous Proteins Used in the Semisynthesis of Protein Transposons by RP-HPLC and ESI-MS. Top:** Analytical C-18 RP-HPLC chromatograms (gradient used: 20-75% B, 10-40 min) (left) for recombinant $Cfa^{N}{}_{m}$ split intein and the corresponding ESI-MS spectra with deconvoluted spectra shown in insert (right). **Middle:** Analytical C-18 RP-HPLC chromatograms (gradient used: 20-75% B, 10-40 min) (left) for mutant $Npu^{C}$-MBP used in expressed protein ligation with split inteins and the corresponding ESI-MS spectra with deconvoluted spectra shown in insert (right). **Bottom:** Analytical C-18 RP-HPLC chromatograms (gradient used: 20-75% B, 10-40 min) (left) for mutant $Npu^{C}$-AAFNC-OMe (red) and its methyl ester hydrolysis product (green) used in expressed protein ligation with split inteins and the corresponding ESI-MS spectra with deconvoluted spectra shown in insert (right).

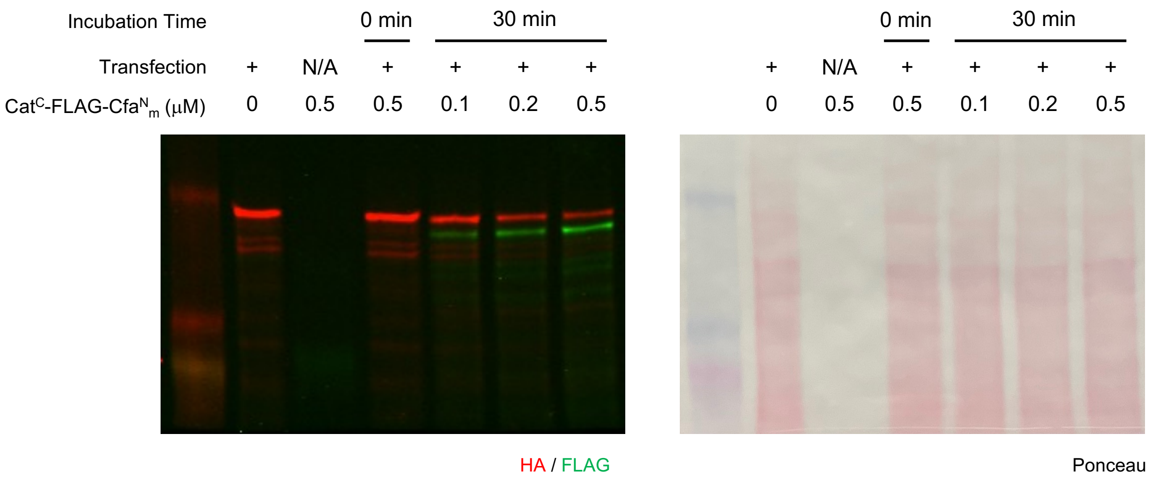


**Fig. S35. Immunoblot Analysis of *In Nucleo* Protein Transposition and Associated Loading Control** *In nucleo* protein transposition reaction between endogenously expressed recipient SMARCA5 in HEK 293T cells containing an embedded HA tag (red signal) and exogenously added transposon construct (0.1–0.5 uM) containing a FLAG tag (green signal) over 30 min. Successful *in nucleo* transposition is denoted by the decrease in the HA signal with a concomitant increase in the FLAG signal. Protein loading is shown with ponceau staining.

#### Uncropped Gels and Blots Used in this Study

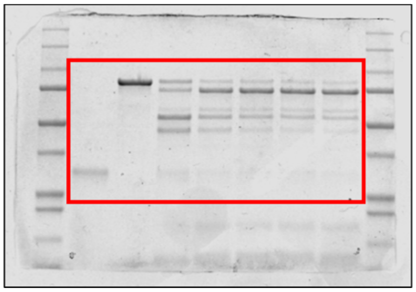

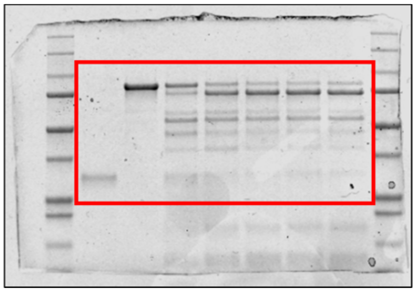

Left: Fig. 2c left and Fig. S4b; Right: Fig. S4d

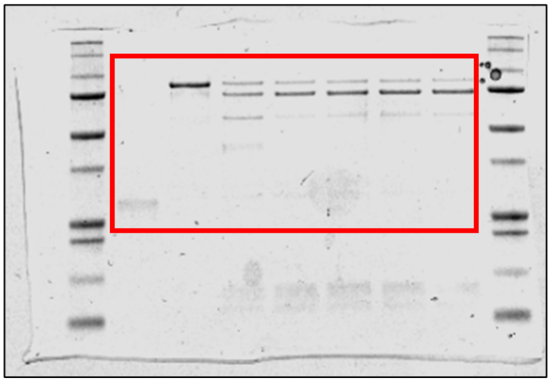

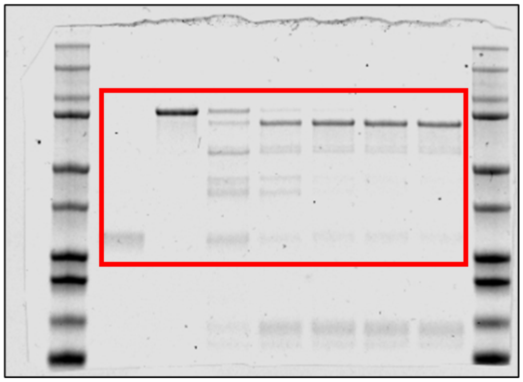

Left: Fig. 2c left and Fig. S5b; Right: Fig. S5d

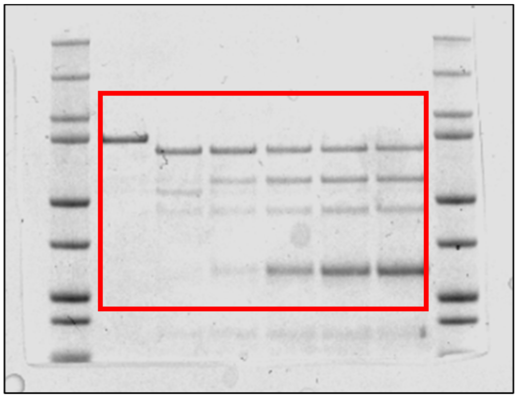

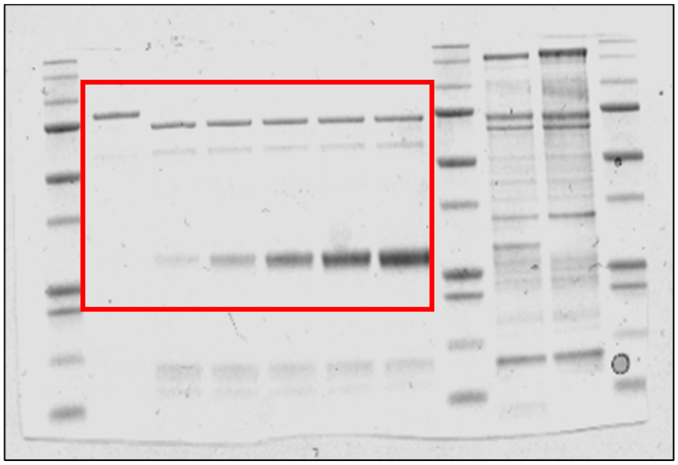

Fig. S11b

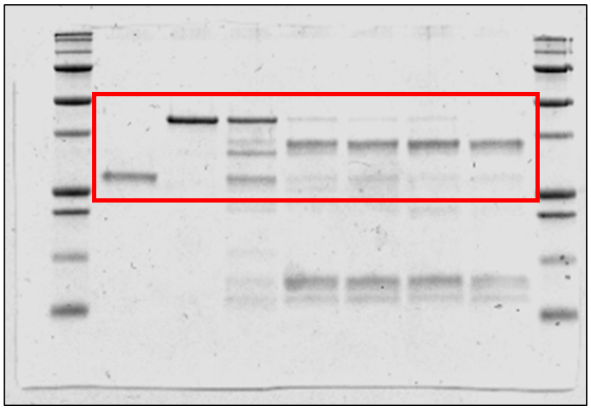

Fig. 3b

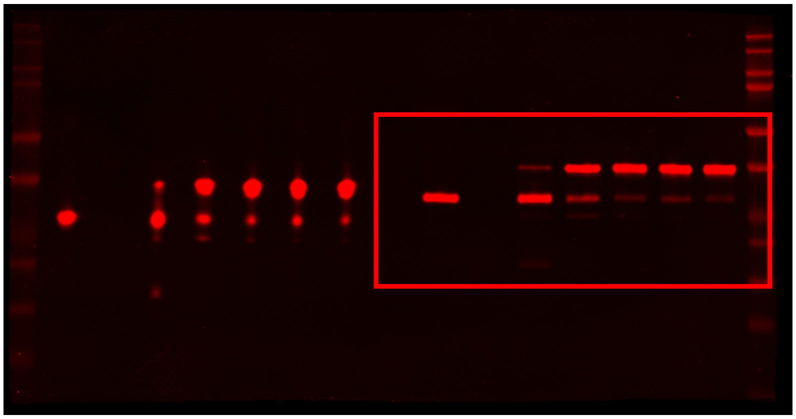

Fig. 3b

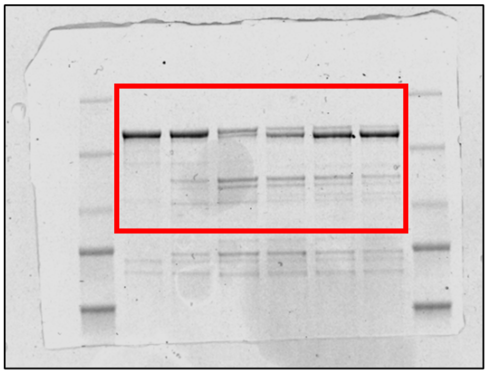

Fig. S14a

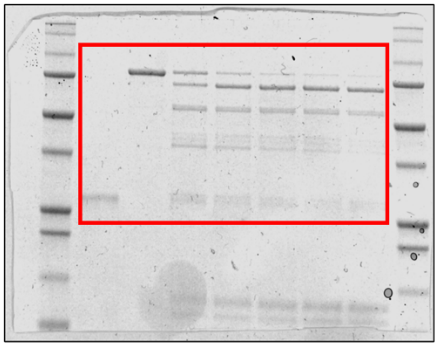

Fig. S17b

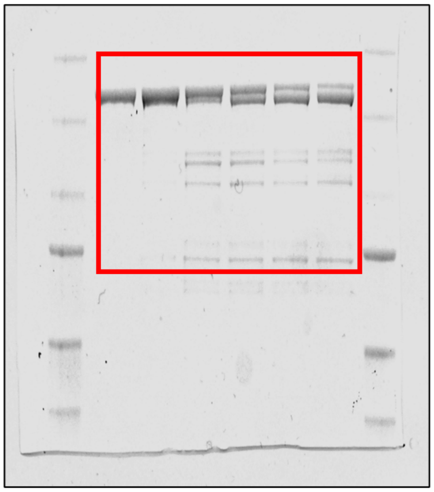

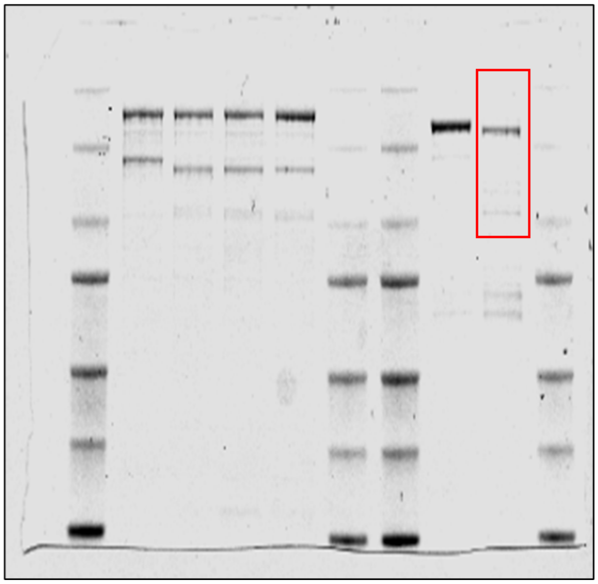

Fig. 3e and Fig. S13a

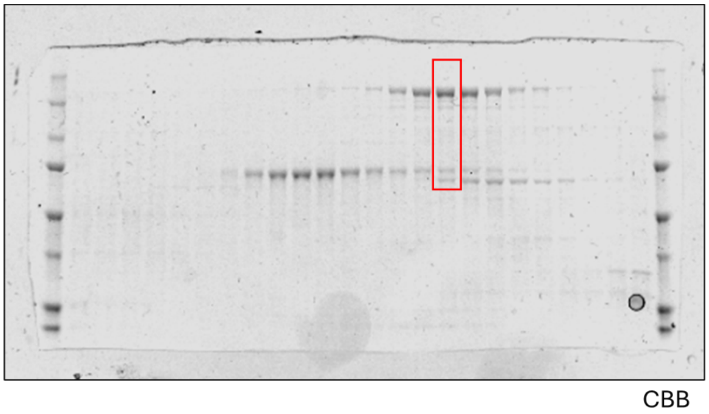


Fig. S13a

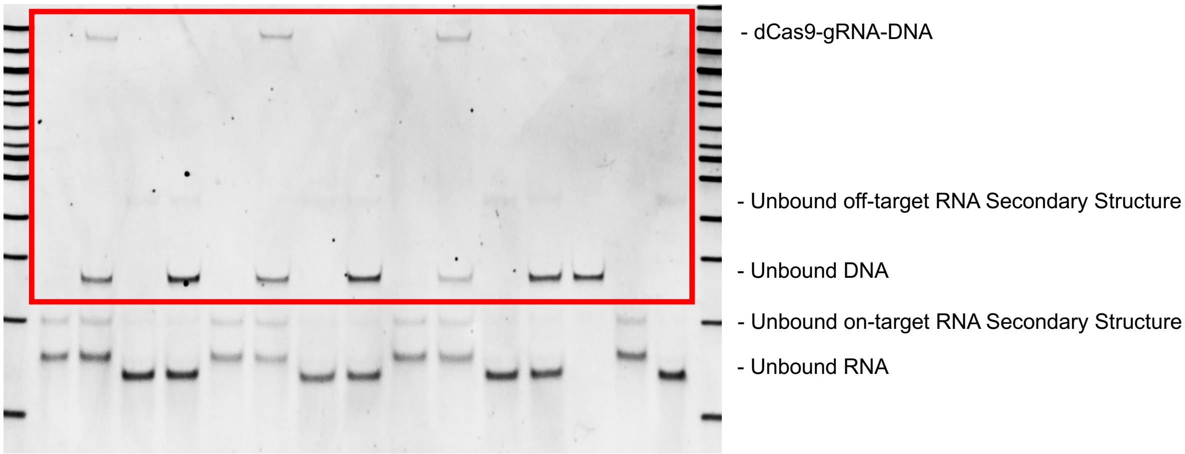


Fig. 3f

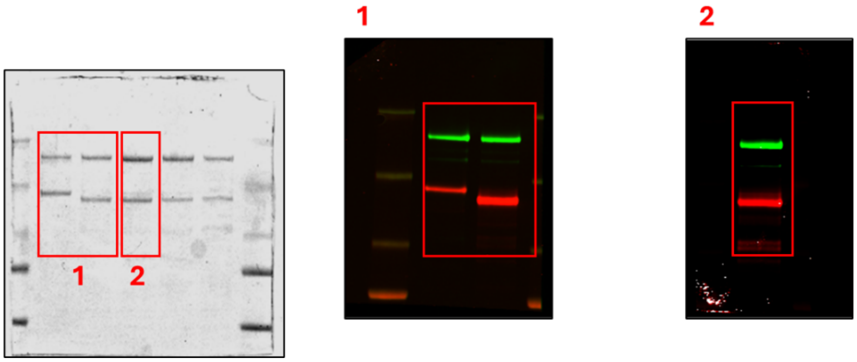


1. Fig. 4b, Fig. 21a
2. Fig. 4b

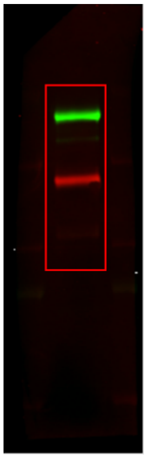

Fig. 4b (Transposition between recipient ACF complex and SMARCA5-transposon B which contains the photoMet743 probe)

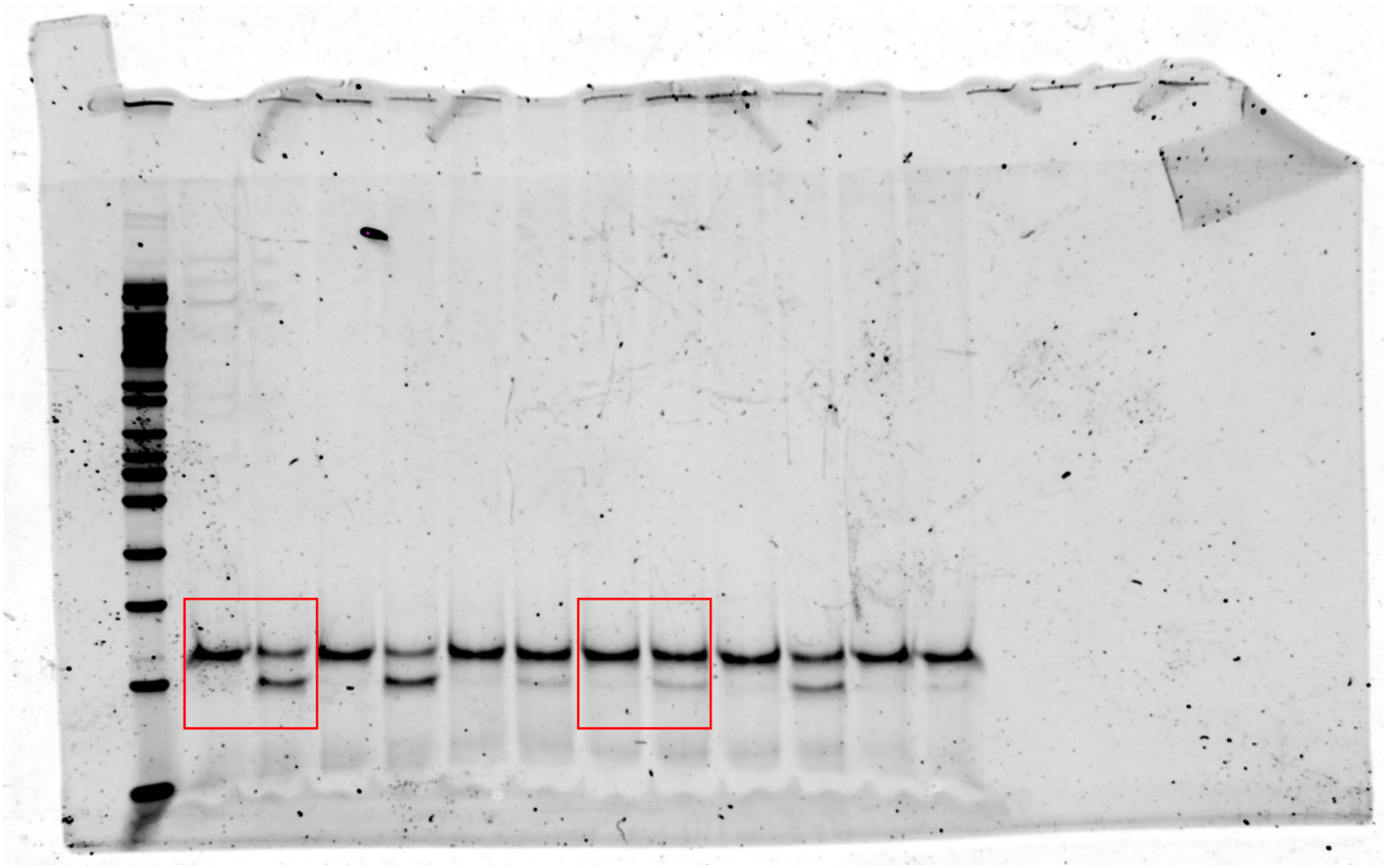

Fig. 4c

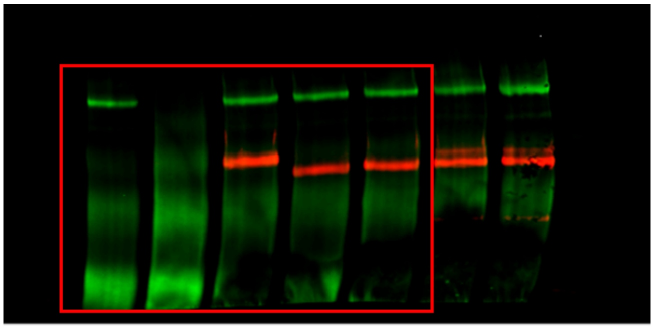

Fig. S22a

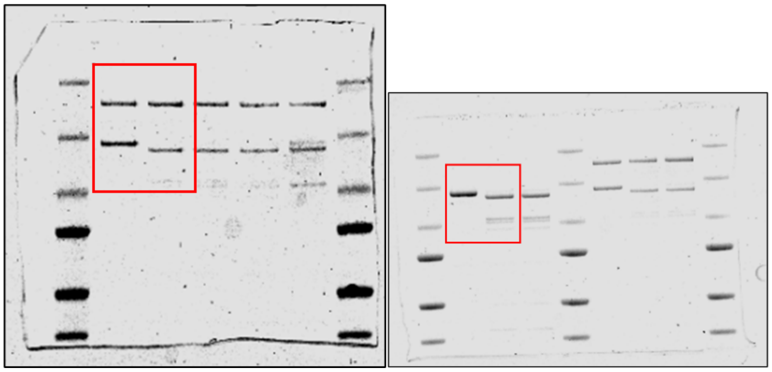

Fig. S22b

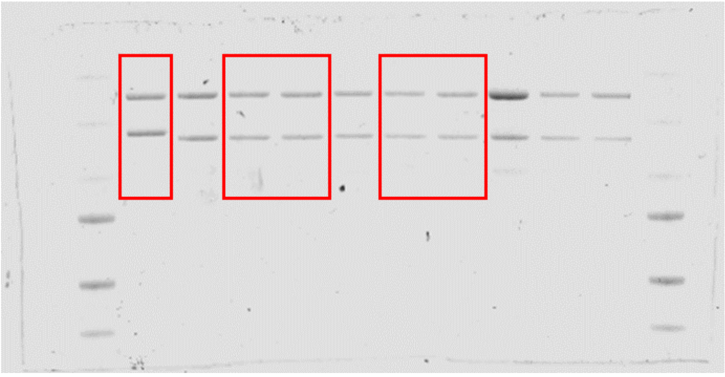

Fig. S22c

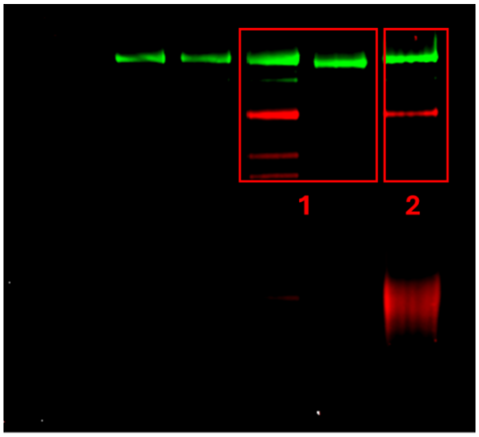

1. Fig. S22c
2. Fig. S22a

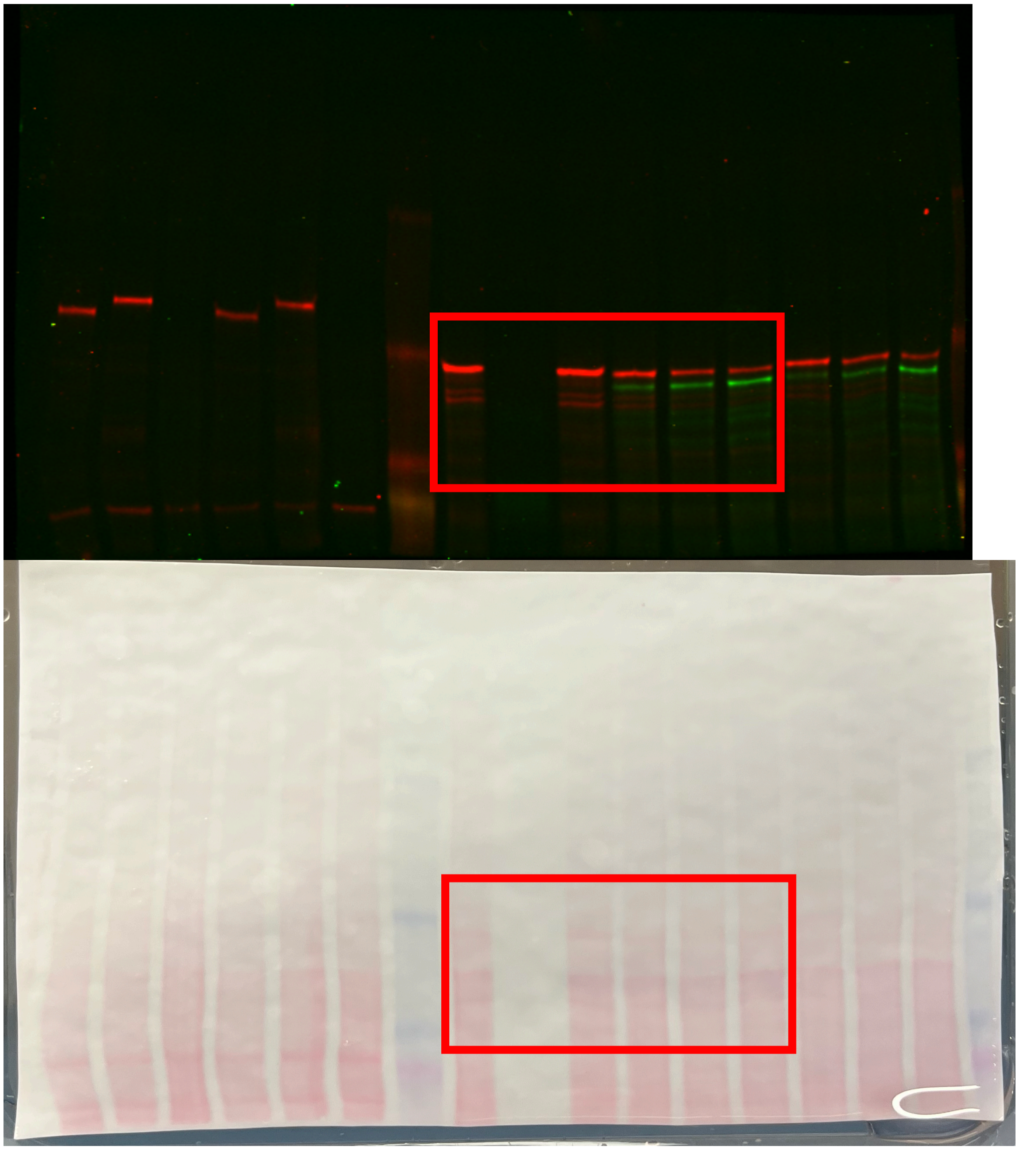
Figs. 4F and S35

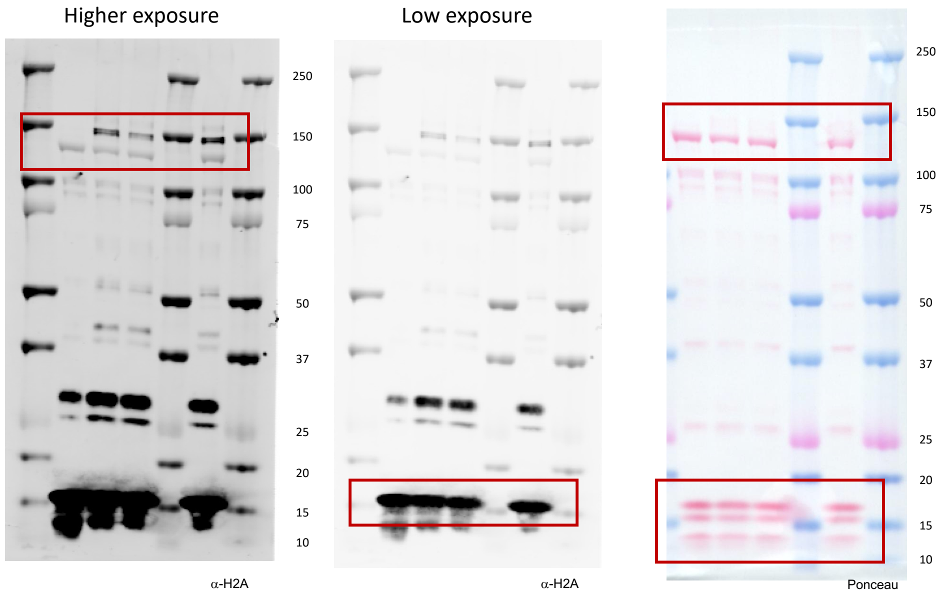


Fig. 4 and Fig. S26

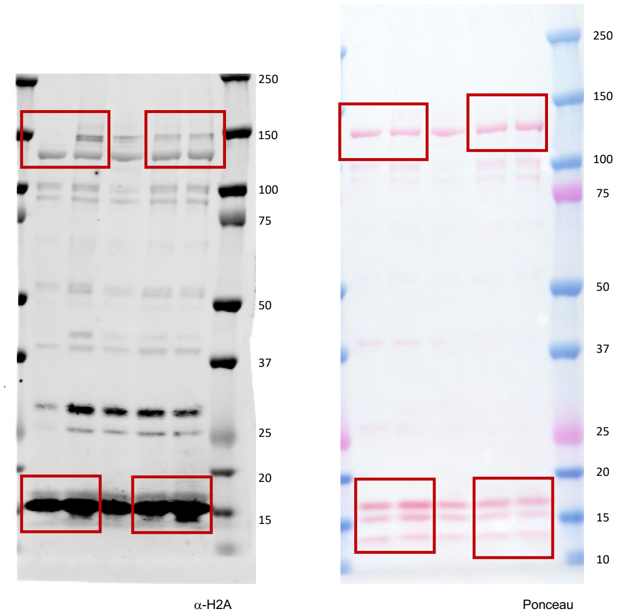


Fig. S26

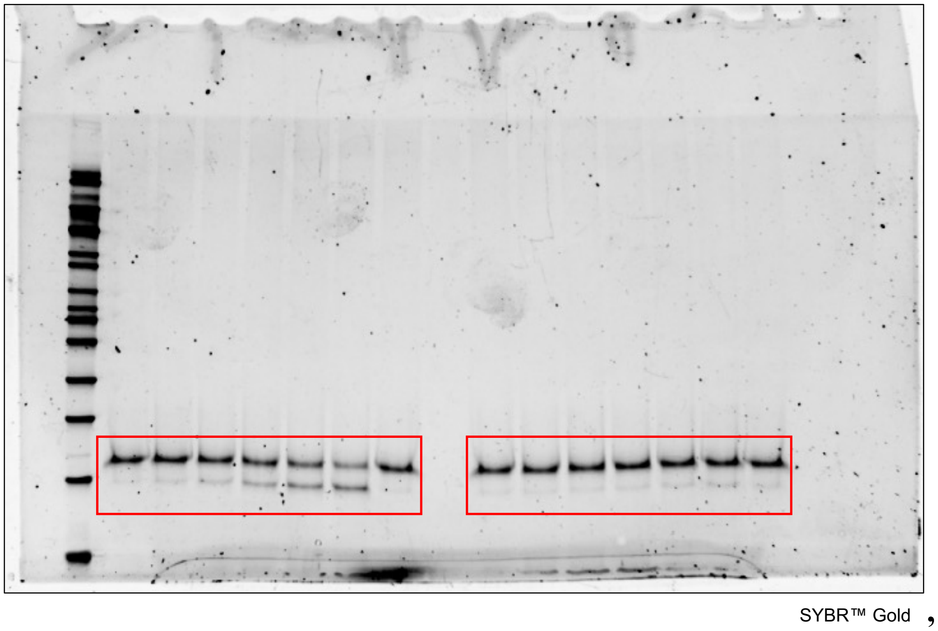


Fig. S31

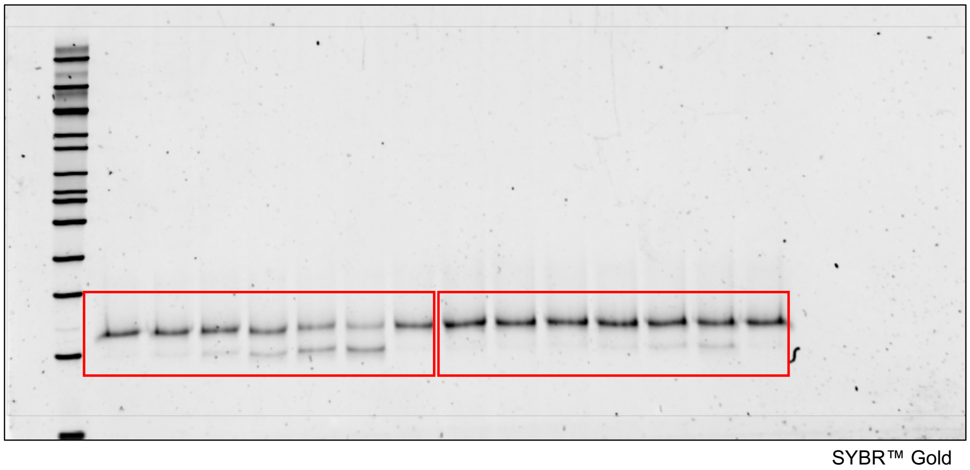


Fig. S31

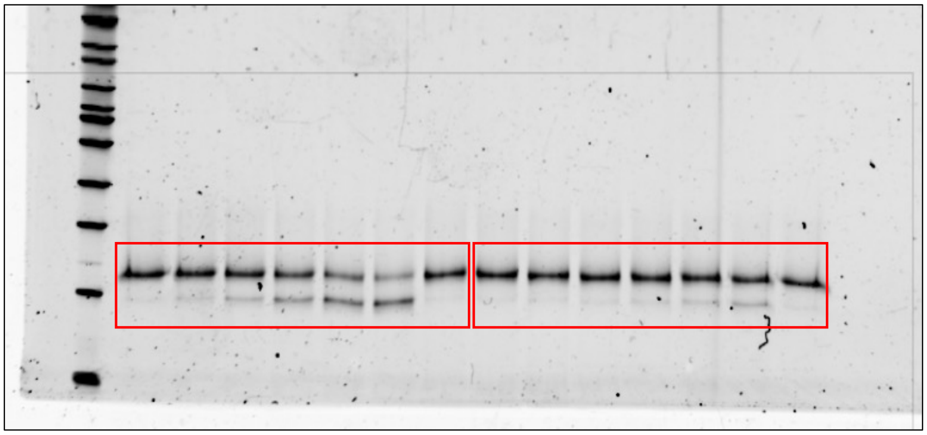


Fig. S31

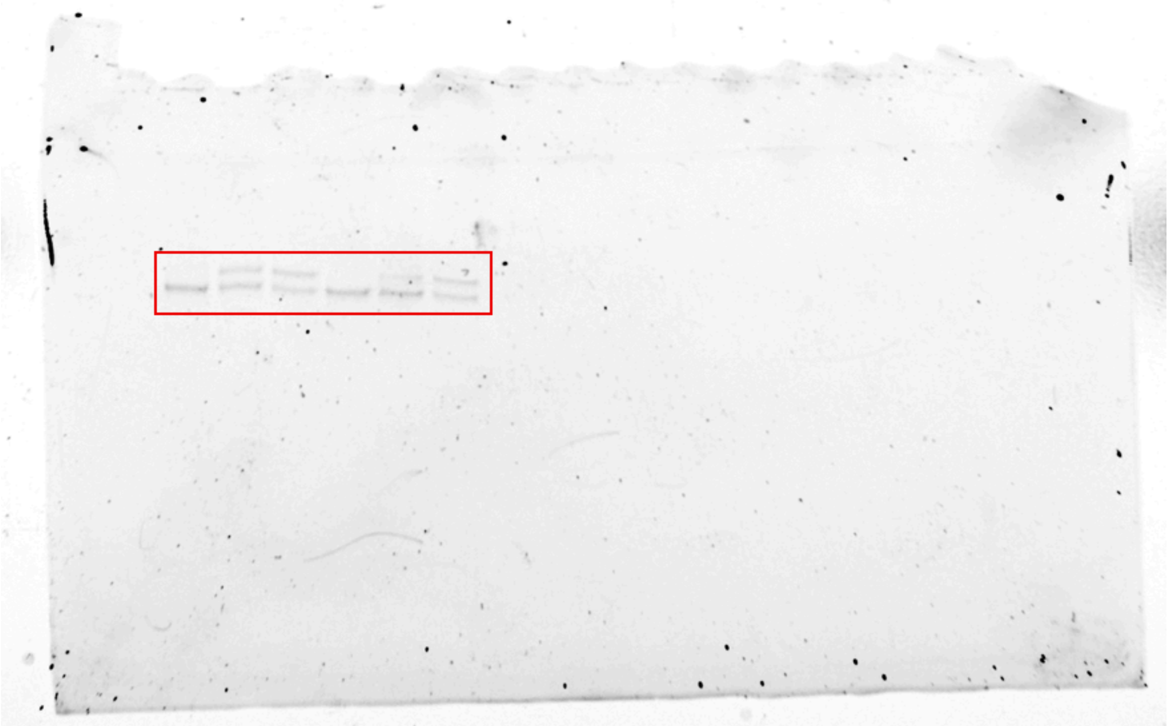

Fig. S32

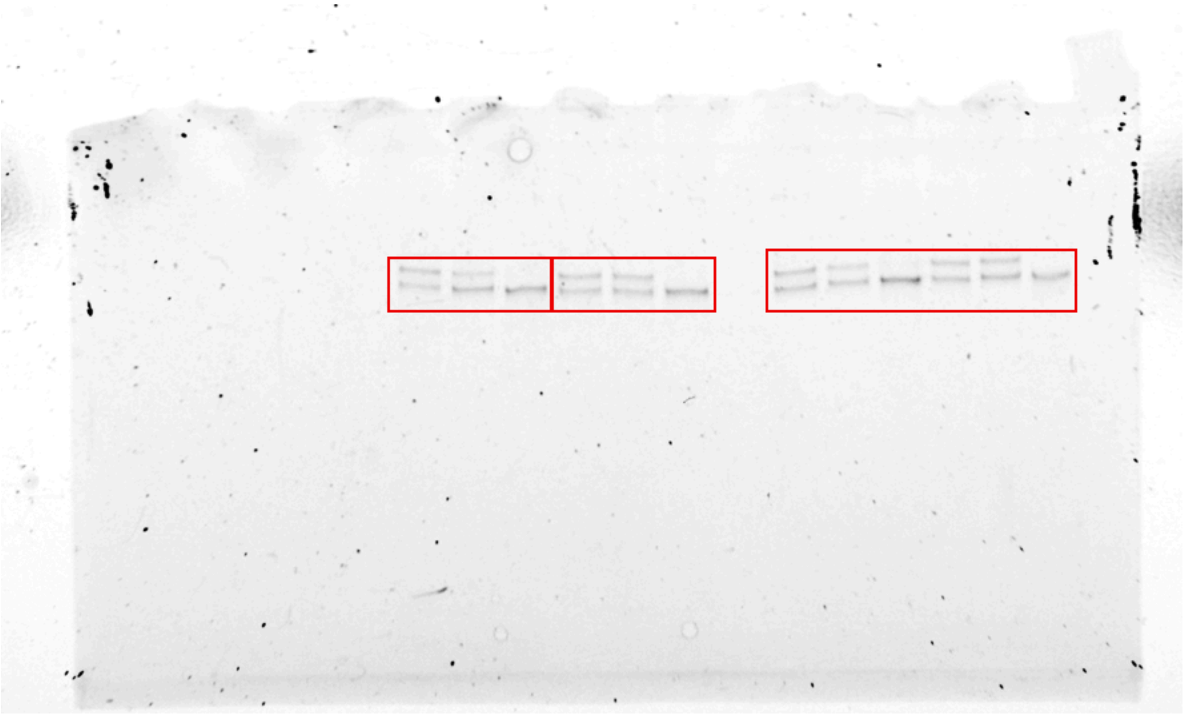

Fig. 4d and Fig. S32

### Full Ion Mass List for LC-MS/MS Experiments

#### Recipient = dCas9, Transposon = HA

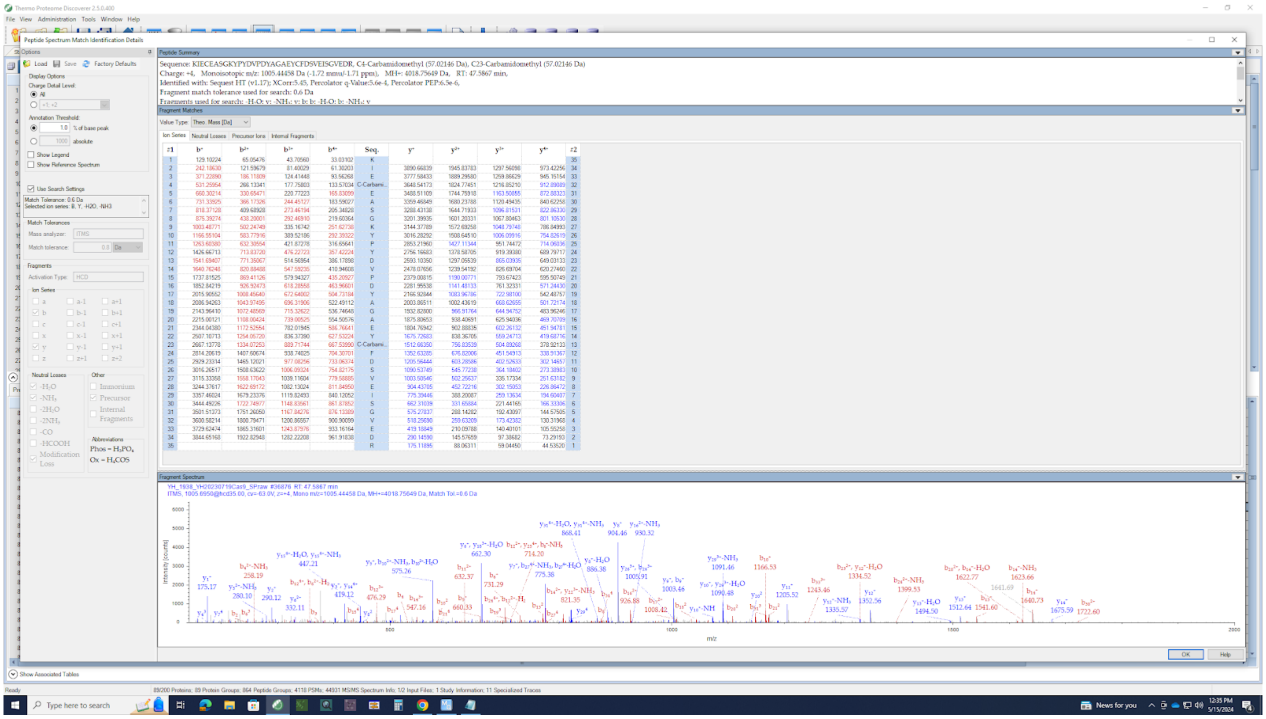


#### Recipient = SMARCA5, Transposon = HA

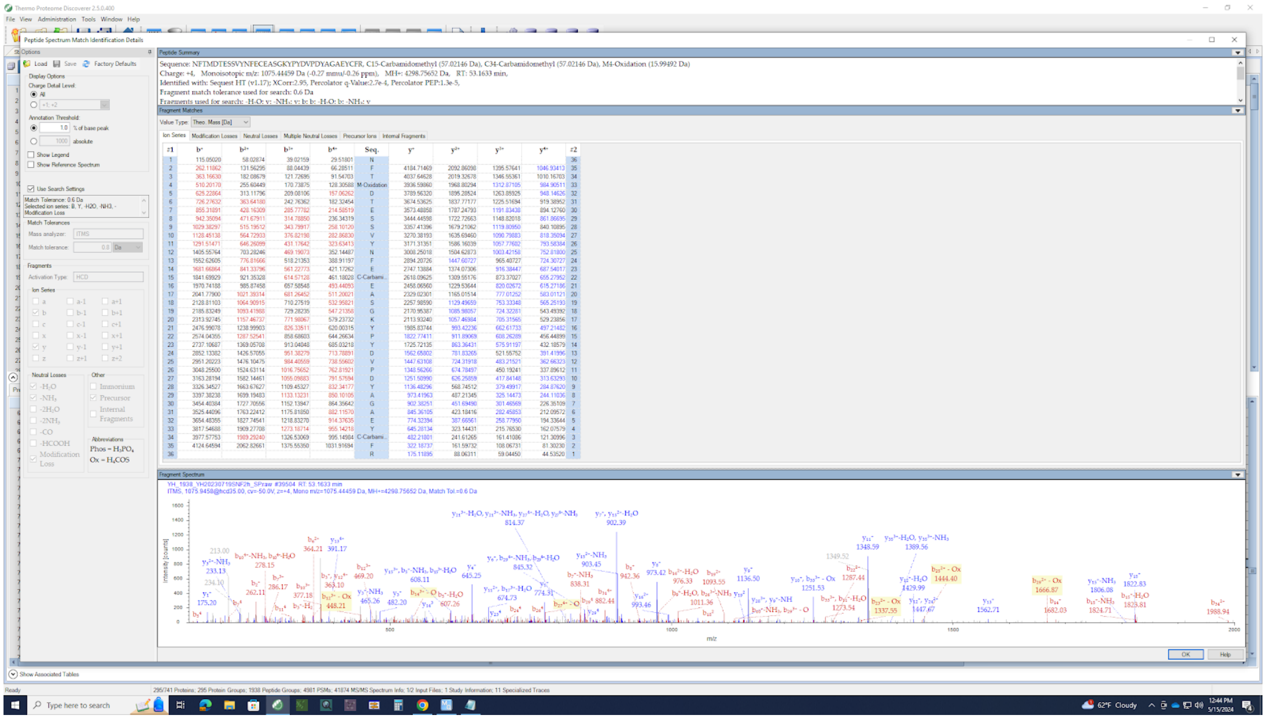
